# Direct neural perturbations reveal a dynamical mechanism for robust computation

**DOI:** 10.1101/2022.12.16.520768

**Authors:** Daniel J. O’Shea, Lea Duncker, Werapong Goo, Xulu Sun, Saurabh Vyas, Eric M. Trautmann, Ilka Diester, Charu Ramakrishnan, Karl Deisseroth, Maneesh Sahani, Krishna V. Shenoy

## Abstract

The rich repertoire of skilled mammalian behavior is the product of neural circuits that generate robust and flexible patterns of activity distributed across populations of neurons. Decades of associative studies have linked many behaviors to specific patterns of population activity, but association alone cannot reveal the dynamical mechanisms that shape those patterns. Are local neural circuits high-dimensional dynamical reservoirs able to generate arbitrary superpositions of patterns with appropriate excitation? Or might circuit dynamics be shaped in response to behavioral context so as to generate only the low-dimensional patterns needed for the task at hand? Here, we address these questions within primate motor cortex by delivering optogenetic and electrical microstimulation perturbations during reaching behavior. We develop a novel analytic approach that relates measured activity to theoretically tractable, dynamical models of excitatory and inhibitory neurons. This computational model captures the dynamical effects of these perturbations and demonstrates that motor cortical activity during reaching is shaped by a self-contained, low-dimensional dynamical system. The subspace containing task-relevant dynamics proves to be oriented so as to be robust to strong non-normal amplification within cortical circuits. This task dynamics space exhibits a privileged causal relationship with behavior, in that stimulation in motor cortex perturb reach kinematics only to the extent that it alters neural states within this subspace. Our results resolve long-standing questions about the dynamical structure of cortical activity associated with movement, and illuminate the dynamical perturbation experiments needed to understand how neural circuits throughout the brain generate complex behavior.

---

Complex behaviors rely on neural computations that are dynamical in nature, in that the brain must maintain and transform the values of internal variables relevant to the task at hand, as well as generate time-varying signals that produce goal-directed movements. The transformations by which recurrently connected circuits shape the evolution of neural population activity—and the task-related signals therein—may be characterized as a dynamical system.^1,2^ This framework of *computation through neural dynamics* maps theories of neural computation onto dynamical motifs that guide neural population activity. Importantly, specific computational hypotheses may be evaluated as falsifiable predictions regarding the evolution of neural states over time.

This perspective has been particularly influential in studies of the motor cortex,^1,3,4^ which serves as an important nexus in the control of skilled arm and hand movements.^5–8^ Temporally complex, heterogeneous firing patterns in individual cortical neurons exhibit lawful dynamics at the population level,^9,10^ generating patterned outputs that help guide movement via lower motor centers^11–13^ (Fig. 1a). Across numerous studies in primates and rodents, three distinct classes of hypotheses have emerged to describe the operation of motor cortex as a dynamical system. First, motor cortex could implement a *reservoir network* (**H1**), supporting a highly expressive collection of basis patterns that are flexibly combined to create complex downstream readouts.^14–22^ Under this view, although neural activity following from any single initial condition might engage only a few dimensions in course of one movement, the circuit retains the concurrent potential to generate activity patterns in many additional dimensions, which would be revealed by increasing task demands^23^ (Fig. 1b). Second, motor cortex could establish a *subspace-structured network* (**H2**), in which activity that lies within a small set of dimensions in the full neural space shapes the evolution of task-related signals (Fig. 1c). Such low-dimensional subspace structure is present in model networks with low-rank connectivity^24–27^ and has motivated the development of methods to estimate low-dimensional dynamical systems in a latent variable space directly from neural population recordings.^9,10,28–31^ Low-dimensional subspace-structure can also emerge in task-optimized artificial systems when appropriately regularized.^32–34^ Lastly, motor cortex may be governed by *path-following dynamics* (**H3**), in which the neural state is constrained to move along an externally configured path. In the context of motor control, previous work has proposed that the motor cortex might serve to activate specific motor programs implemented by recurrent circuitry in the spinal cord.^35^ Under this hypothesis, motor cortical activity would be pushed along specific trajectories dictated by evolving movement instructions,^36,37^ sensory feedback,^38^ and predictive internal models within the cerebellum,^39^ conveyed from other cortical regions and via the motor thalamus.^4,40,41^

**Fig. 1.**
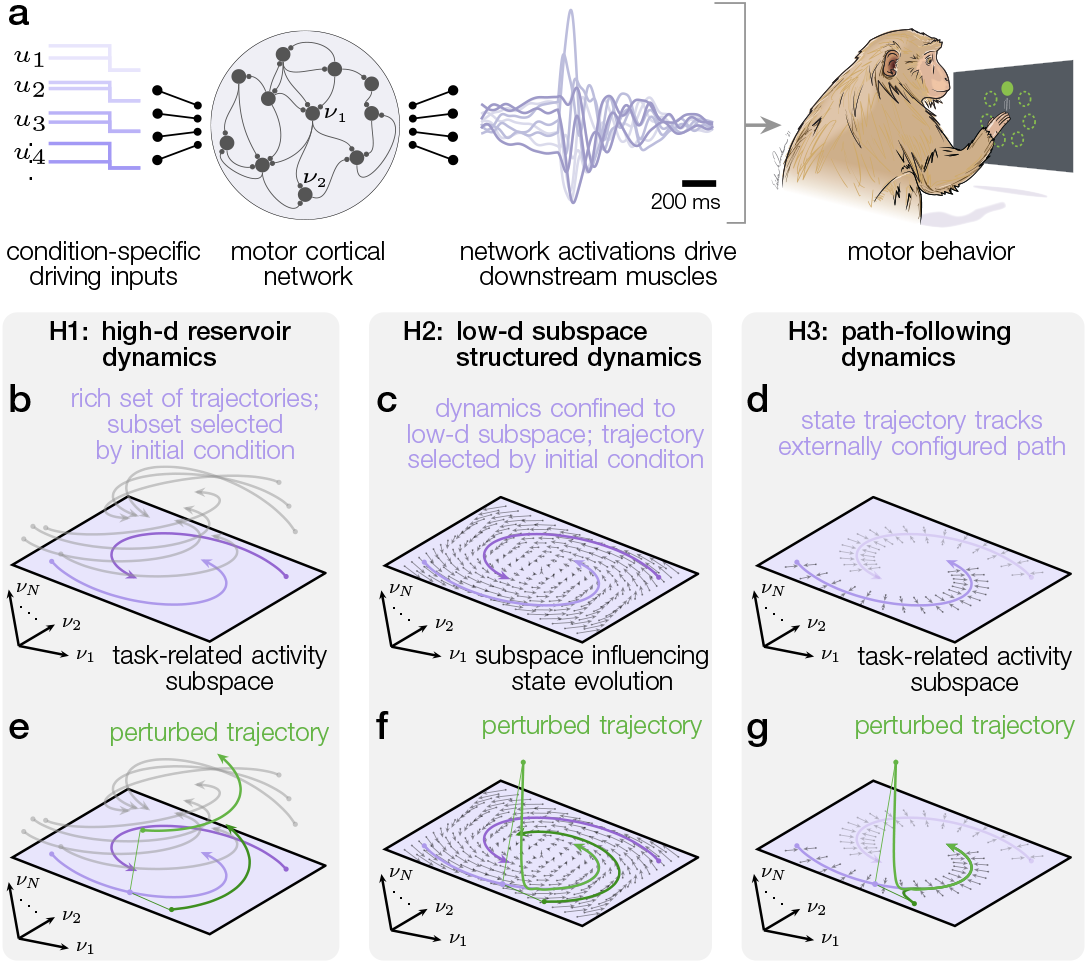
Hypotheses for the dynamical systems underlying the control of movement. **a:** A dynamical system creates heterogeneous activity patterns in response to initial conditions and inputs. The activity patterns drive downstream muscle activations, which ultimately result in the desired movement. **b,e: H1**. The network dynamics are a high-dimensional reservoir creating the activity patterns needed to produce complex movements. For simple tasks and inputs, observed activity patterns may only explore a lower-dimensional subspace (colored plane). For other inputs, the network has the ability to explore additional dimensions (grey trajectories), reflecting the underlying high-dimensionality of network dynamics (**b**). Since the network is sensitive to a high-dimensional space of inputs and initial conditions, a perturbation is likely to unveil this structure and result in long-lasting effects (**e**). **c,f: H2**. The network dynamics implement specific computations within lower-dimensional subspaces. Inputs are targeted towards this space and there is only a limited set of inputs and initial conditions that will affect the future state evolution of the dynamical system (**c**). Unless perturbations are specifically targeted towards this low-dimensional space, they are unlikely to engage with the low-dimensional structure of the dynamical system (**f**). **d,g: H3**. Network dynamics are configured to follow a single trajectory. All directions in network space are set up to decay rapidly towards this path (**d**). As a consequence, any perturbation to the local neural state is expected to decay back towards the established trajectory (**g**). ↑ Go back.

Each of these proposals is consistent with the low-dimensional neural activity patterns that have been observed during reaching tasks.^9,14,32^ However, they make different predictions about how experimental perturbations of the neural state would engage with local cortical dynamics and alter the future evolution of neural population activity. In reservoir networks, which retain the concurrent potential to express basis patterns in many different dimensions, perturbations of neural state should engage new dynamical modes and thus evoke complex, long-lasting transients (Fig. 1e). In subspace structured networks, where activity is driven by the state within a low-dimensional subspace, only perturbations that affect this subspace should elicit complex, long-lasting effects, while perturbations along all other dimensions should fail to engage with the circuit dynamics (Fig. 1f). Lastly, in a path-following network, all perturbations away from the externally-configured trajectory will decay back rapidly (Fig. 1g).

To select among these competing hypotheses, we performed targeted circuit perturbations using optogenetic stimulation and electrical intracortical microstimulation (ICMS) in dorsal premotor and primary motor cortices of rhesus macaque monkeys engaged in an instructed delay center-out reaching task. Optogenetic stimulation created a large displacement in neural population state that decayed rapidly and did not alter reach kinematics. These optogenetic stimulation responses were inconsistent with the predictions of high-dimensional reservoir dynamics (**H1**). In contrast, ICMS produced heterogeneous effects on reaching kinematics and exhibited more temporally complex interactions with the local dynamics; these long-lasting responses provided evidence against path-following dynamics (**H3**). Statistical estimates of a local dynamical system with underlying low-dimensional structure embedded in a high-dimensional network of excitatory and inhibitory cells were predictive of the time-courses of recovery from both perturbation modalities, supporting a view of motor cortex as a dynamical system with low-dimensional, subspace-structured dynamics (**H2**). Additionally, we found that ICMS distorted the underlying geometry of task-relevant latent neural states to an extent correlated with the resultant behavioral effects. This non-additive mechanism revealed how these electrical perturbations—not targeted to specific neural dimensions—influenced the low-dimensional subspace that governs task dynamics. Collectively, these findings significantly constrain the space of hypotheses about the exact nature of the dynamical system that shapes motor cortical activity, reveal new insights into how distinct stimulation modalities engage with cortical dynamics, and suggest a general mechanism by which subspace-structured low-dimensional dynamics support robust computation in neural circuits.

## Optogenetic stimulation during reaching

In two rhesus macaques (Q and O), we used AAV5-CaMKIIα to target the red-shifted, excitatory opsin C1V1_⊤⊤_ to excitatory neurons^42,43^ throughout the gyral motor cortex, comprising dorsal premotor (PMd) and primary motor (M1) areas. We inserted a primate-optimized coaxial optrode^44^ at one of several discrete locations in these areas, along with several independently positioned microelectrodes (Fig. 2a). This strategy delivered repeatable optogenetic perturbations across experimental sessions while simultaneously recording the responses of the local neural population. As the monkeys performed a center-out reaching task with an instructed delay period, we delivered 200 ms continuous pulses of optogenetic stimulation on randomly interleaved trials at a discrete set of salient timepoints within the task (Fig. 2b). Consistent with previous reports,^45–47^ optogenetic excitation of motor cortex failed to alter reaching kinematics at any time during the trial (Fig. 2c, Extended Data Fig. 1a,b). We observed a modest slowing of reaction time when stimulation was delivered to PMd coincident with the visual go cue, indicating a disruption of motor preparation (Extended Data Fig. 1c).

**Fig. 2.**
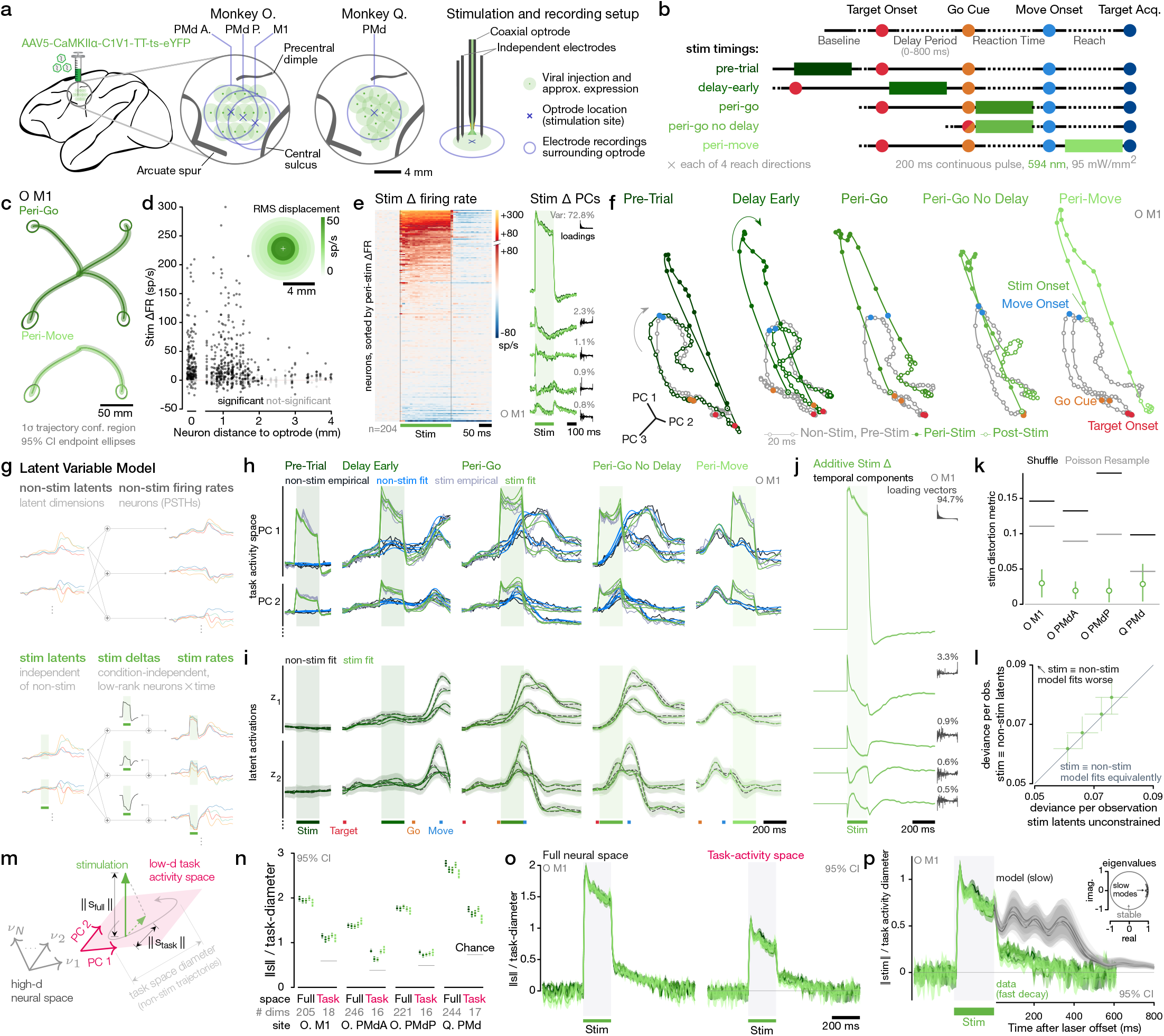
Optogenetic perturbations of motor cortex additively displace task activity but decay quickly. **a:** Schematic of optogenetic stimulation setup with simultaneous electrode recording of surrounding neurons. Population responses were collected at four optrode sites in two monkeys. Anatomical landmarks are approximate. **b:** Diagram of stimulation timings within instructed-delay reaching task. **c:** Peri-move optogenetic stimulation does alter reach kinematics. **d:** Stimulation-evoked change in firing rate (Δ FR) vs. neuron distance from stimulation. Neurons at 0 were recorded on the coaxial optrode. Inset: r.m.s. firing rate displacement of neural state by stimulation within population of neurons recorded in annuli of increasing distance from optrode. **e:** Left: Pre-trial stimulation responses of all recorded neurons at O. M1 site, sorted by mean ΔFR during stimulation period. Right: Principal components of stimulation-evoked ΔFR. Timecourses colored by stimulation timing as in (**b**); black stem plots show PC loading vectors over neurons, sorted by ΔPC 1. **f:** State space visualization of stimulation evoked displacements of neural trajectories within the task-activity space (PCs of non-stimulated trial-averages) at site O. M1. **g:** Schematic illustration of latent variable model to distinguish additive effects of stimulation from changes in underlying task-related latent variables. **h:** Empirical and LVM-fitted trial-averaged firing rates projected into leading two task activity space dimensions. Stimulation timings are separated into columns, with each trace corresponding to a reach direction. **i:** Fitted latent activations for the leading two latent dimensions. **j:** Timecourses of the additive delta term which reflects a condition-invariant stimulation induced change in firing rates. Inset: loading vectors mapping each component into neural space together with the percentage of variance each component explains in the empirical stimulation-delta. **k:** Shape similarity metric quantifying distortion between non-stimulated and stimulated latents. Shuffle and Poisson resample lines indicate significance thresholds (*α* = 0.05) under two null hypotheses, see Methods §8.3. **l:** LVM goodness of fit (as Poisson deviance) with stimulation latents fit independently of non-stimulation latents (horizontal), or constrained equal (vertical). Filled circles indicate statistically significant differences (*α* = 0.05); error bars indicate jackknife standard error. **m,n:** Empirical stimulation vector length for each stimulation timing, in the full neural space and projected into task activity space, normalized by task-space diameter. Numbers in **m:** denote dimensionality of each space. **o:** Timecourses of normalized stimulation vector length for O. M1 in the full neural space (left) and projected into the task activity space (right). Each trace represents a stimulation timing colored as in (**b**). **p:** LDS model predictions (grayscale) vs empirical decay of stimulation vector in task-activity space (green, same as (**o**), right). Inset: eigenvalues of LDS dynamics matrix; circle outlines stable region. ↑ Go back

In the local neural population, optogenetic stimulation rapidly evoked strong responses in a large region extending 2 mm of the optrode source (Fig. 2d,e), presumably reflecting both directly stimulated and transsynaptically driven neurons. At the population level, the perturbed firing rates exhibited large deflections along the leading principal components of non-stimulated activity.

We sought to characterize the impact of optogenetic stimulation using a latent variable model (LVM). Firing rates were modeled as linear functions of a set of lower-dimensional time-varying latent variables with condition-dependent trajectories, combined with an additive term during and after stimulation that was invariant across reach directions and stimulation times (*stimulation* Δ, Fig. 2g, Extended Data Fig. 2a-d; see Methods §8). The LVM accurately captured neural responses for both non-stimulated and stimulated conditions (Fig. 2h, Extended Data Fig. 2e,f). Under stimulation, the inferred latent variable trajectories were indistinguishable from the trajectories in the corresponding non-stimulation condition (Fig. 2i). Remarkably, stimulation’s effects on firing rates were captured entirely within the additive stimulation delta term (Fig. 2j). We verified that the latent trajectories were not distorted under optogenetic stimulation using a shape similarity metric^48^ (not significant against either a permutation test or a null distribution sampled from synthetic firing rates computed with identical stimulation and non-stimulation latents, Fig. 2k shuffle and Poisson resample, respectively; see Methods §8.3). Furthermore, the LVM’s goodness of fit was not significantly reduced when fit with a constraint enforcing identical stimulation and non-stimulation latents (Fig. 2l). Collectively, these results demonstrate that optogenetic stimulation resulted in a purely additive effect, translating neural states in parallel without distorting the underlying geometry of task related activity.

The magnitude of the stimulation-induced displacement vector was large, exceeding the task diameter, which we defined as the largest distance between any two neural states on the non-stimulated neural trajectories (Fig. 2m,n, Full space). We then identified the low-dimensional task activity space which captured the majority of task-related variance on non-stimulated trials using PCA. The projection of the stimulation vector within the task activity space was also larger than expected by chance for a randomly oriented perturbation vector (Fig. 2n, Task space). The displacement distance increased immediately at laser onset and decayed rapidly after the offset of stimulation (Fig. 2o).

## Rapid decay of optogenetic stimulation constrains hypothesis space

This rapid decay of the perturbation vector indicates that optogenetic stimulation failed to evoke complex, long-lasting transients in motor cortex. This is an unlikely outcome under the reservoir network hypothesis (**H1**) where dynamics are dominated by slow and structured responses along most input dimensions. In a representative network model from the reservoir network class,^14^ random additive perturbations to excitatory cells triggered long-lasting effects. These predicted responses were qualitatively different from the observed neural responses (Extended Data Fig. 3), arguing against the presence of high-dimensional reservoir dynamics in the motor cortex (Fig. 1b,e, **H1**).

In a network governed by subspace-structured dynamics (**H2**), dynamically-potent dimensions are confined to a low-dimensional *task dynamics* subspace. If the optogenetic stimulation were misaligned with this task dynamics subspace, a fast decaying transient would result, which could be consistent with the observed neural responses. However, although the optogenetic stimulation decayed rapidly, it exhibited a large projection into the task activity space where neural population activity displayed variance during reaching. It is typically assumed that the task activity space—where task-related activity exhibits high variance— is identical to the the task dynamics space—where activity is shaped by slow, task-related dynamics. Consequently, we would expect that optogenetic stimulation would elicit slowly-decaying transients that match the autocorrelation timescales of the observed task-related activity (e.g., Fig. 2h, black traces). Consistent with this prediction, a latent linear dynamical system (LDS) model fit to the non-stimulated firing rates learned to produce slow task-related dynamics and consequently predicted slowly decaying transients following stimulation offset (Fig. 2p, Extended Data Fig. 4a-c)

This mismatch in timescales requires that if motor cortex is shaped by subspace-structured dynamics (**H2**), then its task dynamics space active for reaching movements must be oriented in neural space differently from the task activity space. Such a misalignment is characteristic of non-normal dynamical systems governed by a dynamics matrix whose eigenvectors are not orthogonal to each other.^49,50^ Indeed, when we fit the LDS model to neural activity from both non-stimulated and stimulated trials, these models reproduced both the slow evolution of task-related neural activity and the rapid decay of the empirical stimulation via a non-normal dynamical mechanism (Extended Data Fig. 4d,f,h).

The LDS solution demonstrates that the observed data may be *consistent* with subspace-structured dynamics (**H2**, Fig. 1c), yet, these non-normal dynamics did not arise without explicitly fitting to stimulation data (Fig. 2p). When we shuffled the elements of the stimulation vector used to perturb the model, simulating stochasticity in opsin expression over neurons, the LDS model failed to generalize, predicting incorrect slow decay times following perturbation (Extended Data Fig. 4e,g). Moreover, the fragile solution learned by the LDS models provided little mechanistic insight into *why* such non-normal structure would arise in the motor cortical network and *how* it orchestrates the rapid decay of the experimental perturbation.

Consistency with a low-dimensional dynamical system is also insufficient to rule out the alternative hypothesis of path-following dynamics (**H3**, Fig. 1d). Under **H3**, all perturbed neural states would rapidly return to the imposed path, which would provide a trivial explanation of the observed fast decaying transient. We next examined how and why non-normality and stimulation robustness could arise in motor cortical networks, before evaluating the remaining hypotheses with a second set of perturbation experiments using electrical intracortical microstimulation (ICMS).

## Subspace-structured dynamics in a balanced E/I network

Non-normality arises naturally from interactions between excitatory and inhibitory (E/I) cells in networks whose connectivity obeys Dale’s law.^51^ We wondered whether such E/I-based structure might explain aspects of the measured dynamics, and particularly the interplay of those dynamics with cell-type-specific perturbations. To explore this possibility, we developed a novel modeling framework to capture the low-dimensional dynamical activity observed in random samples of motor cortical neurons within an underlying E/I circuit.

The model comprised a high-dimensional (N=1500–2500 units), linear network representing the putative motor cortical circuit (Fig. 3a). Network connections were constrained to satisfy two principles of neurobiological connectivity: Dale’s law^52^ and E/I balance.^53–55^ The network was optimized under these constraints to reproduce the low-dimensional, subspace-structured activity patterns representative of motor cortical dynamics. In particular, we selected a low-dimensional subspace of size K=N/100 at random within the high-dimensional network space, which we designated as the task dynamics subspace, and required network dynamics to be *self-contained* within this subspace. Self-containment ensures that the future evolution of activity within the task dynamics space depends only on the projection of the network state into that subspace. As such, it forms a minimal constraint ensuring that the resulting network evolves with Markovian dynamics within a low-dimensional subspace. More stringent constraints on the network dynamics, such as requiring the connectivity to be of low rank, automatically satisfy this constraint (Extended Data Fig. 5d). We then optimized the E/I network connectivity matrix to maximize the likelihood that a linear readout from this task dynamics space matched the measured neural responses. This approach ensured that the model used only the slow and structured activity within the low-dimensional task dynamics space to produce the activity patterns contained in the observed data, while the rest of the network dynamics largely maintained random structure of the initialization (Extended Data Fig. 5f). The low-dimensional structure was evident in the activity of random samples of network units, which could all be used to reproduce the measured neural responses used to fit the model (Extended Data Fig. 5e). Thus, the model recapitulated the observation that motor cortical responses during simple center-out reaching movements are generic, in the sense that population activity exhibits highly similar structure across experimental sessions, recording sites, and animals and from study to study^23,56,57^ (Extended Data Fig. 5a).

**Fig. 3.**
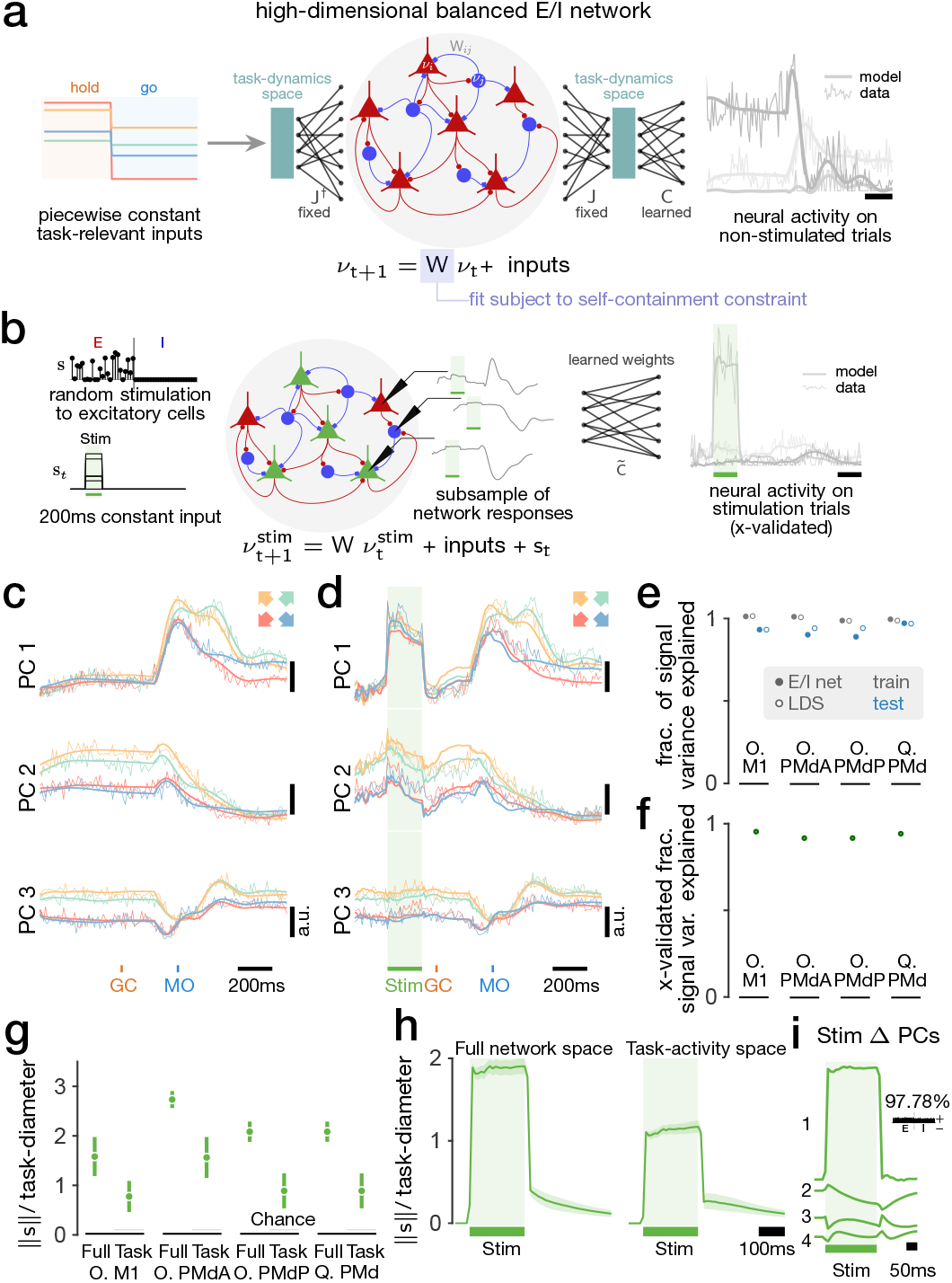
E/I network model recapitulates stimulation responses in recorded data. **a:** A high-dimensional balanced E/I network model of motor cortical activity. The network receives constant target-specific inputs throughout hold/go and is constrained to exhibit self-contained low-dimensional dynamics in a randomly chosen subspace (task-dynamics space). A readout from activity in this subspace is used to reconstruct PSTHs recorded in the absence of stimulation. **b:** After fitting, the network responses to optogenetic stimulation are evaluated. Stimulation is modeled as a constant input targeting a random subset of excitatory cells in the network with randomly chosen amplitudes. A subsample of the network units is used to linearly reconstruct the stimulation responses of recorded units in a leave-one-stimulation-timing-out cross-validated manner. **c:** Example model evaluation predicting forward from learned initial condition on training data for Monkey O. PMdP. **d:** Example cross-validated model reconstructions of stimulation responses for Monkey O. PMdP. **e:** The fraction of signal variance explained for training and held out test data for the E/I network model and an analogous low-dimensional LDS model. **f:** The cross-validated fraction of signal variance explained by the E/I model reconstructions (leave-one-condition-out). **g:** The mean task-diameter normalized stimulation distance during stimulation. Error bars indicate ± one standard deviation across 100 random stimulation patterns for each of 20 networks trained under different random initializations. **h:** The task-diameter normalized mean stimulation distance vs. time in the full network space, and projected into the task-activity space of network units for an example network trained to reproduce Monkey O. M1 PSTHs, averaged across 100 random stimulation patterns. **i:** The top four principal components of example network responses to the stimulation input to excitatory cells. The inset shows the loading weights and fraction of variance explained of the leading principal component of the stimulation response. ↑ Go back

The E/I model was fit exclusively to data from trials without stimulation. After fitting, the network’s responses reproduced the recorded data closely (Fig. 3b,d). Even though the E/I network model was much higher dimensional than the recorded data, it did not overfit, generalized well to held-out test data, and performed similarly to a low-dimensional LDS model in terms of the fraction of signal variance explained (Fig. 3e). Full details on the model description and fitting procedure are provided in Methods §10.

## E/I network model reproduces optogenetic stimulation response features

We probed the network’s responses to cell-type specific perturbations that resemble the optogenetic stimulation of our experiment. Once the network model had been fit, optogenetic stimulation was modeled with additive noisy, positive input patterns targeting random subsets of excitatory cells in the E/I network (Fig. 3b). Even though they had not been used to fit the network parameters, the specific patterns of stimulation response of the network closely matched those seen in the measured data. To demonstrate this, we reconstructed the measured PSTHs on stimulation trials linearly from random sparse subsamples of the network variables at the same time as the task-related PSTHs (Fig. 3b). The reconstructions were quantitatively robust, generalizing to held-out combinations of reach target and optogenetic stimulation timing (Fig. 3d,f). The network response to stimulation inputs reproduced the qualitative features of the recorded data without any further adjustment of the dynamics (Fig. 3g-i). As in the recorded data, the network stimulation responses had a much greater projection into the leading dimensions of task-related variance than would be expected by chance (Fig. 3g,h). The perturbation decayed rapidly after the stimulation input ended, and there was little effect on the future evolution of network activity, despite the apparent overlap with task-related variance (Fig. 3h,i). Furthermore, the stimulation response pattern qualitatively matched what was observed empirically in neural activity (Fig. 2f, Fig. 3h).

## E/I network model establishes mechanism underlying stimulation responses

The effectively parameter-free agreement between the network behavior and the cortical measurements suggested that the observed response to optogenetic activation may be fundamental to balanced E/I networks that implement low-dimensional dynamical computations. To understand how and why these observed response features arise, we isolated the behavior of different input components in the linear network. We first considered how a single stimulation input pattern interacts with the linear network dynamics (Fig. 4a). The dynamics initially amplify the stimulation input into a response where both E and I cells are activated. This E/I co-activation is the dominant response pattern. Any deviations around it can be viewed as random, since these depend on randomness in the stimulation input (e.g., due to opsin expression or laser activation) as well as the network connectivity itself. The E/I co-activation component of the stimulation response then decays rapidly due to E/I balance. The remainder of the response depends on how the random component of the stimulation interacts with the dynamics. The low-dimensional subspace-structured dynamics of the network mean that only inputs within the K-dimensional dynamically potent space (K ≪ N) will produce sustained and structured responses in the network. The expected length of the projection of a random unit-norm input into this space is 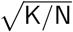. It thus vanishes for large networks that implement low-dimensional dynamical processes through subspace-structured dynamics. Indeed, we find that most of the stimulation responses lie outside of the task dynamics space of the network and that the response component within the task dynamics space is at chance level (Fig. 4a).

**Fig. 4.**
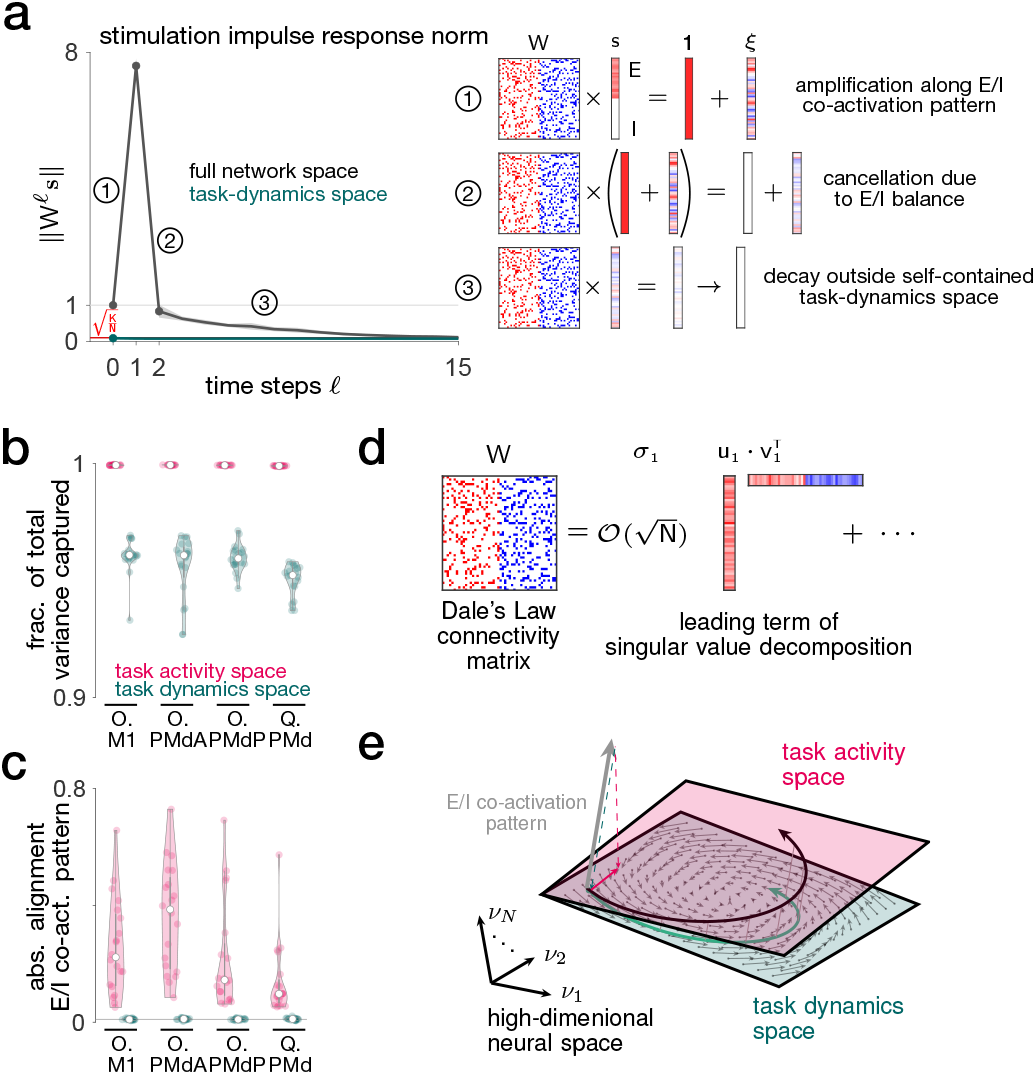
E/I structure and low-dimensional subspace-structure explain stimulation responses. **a:** Left: The mean impulse response norm to 100 randomly chosen stimulation vector patterns in the full network space (black) and projected into the self-contained task-dynamics space of the network (green) together with ±2 standard deviation error tubes. Grey line indicates unit norm, red line indicates chance level for projecting an *N* dimensional random vector into a *K* dimensional subspace. Right: The stimulation impulse response can be understood in terms of three steps: 1) based on the sign structure of the connectivity matrix, positive inputs to E cells get amplified along an E/I co-activation pattern. 2) In the next step of the network evolution, equal activation of E and I cells cancel, leaving only a smaller residual response vector. 3) The residual response vector only has a small by-chance projection into the task-dynamics space of the network and decays with unstructured dynamics. **b:** The total fraction of variance in the network responses captured by the task-dynamics and task-activity space of the the network, for 20 networks trained starting from different random initializations. The task-activity space dimensionality is chosen to match that of the task-dynamics space for each dataset. All networks produced variance in dimensions outside of the task-dynamics space. **c:** Absolute projection of the E/I co-activation pattern (all-ones vector) into the task activity and task-dynamics space. Grey lines indicate chance level. The task-activity space is aligned with the E/I co-activation pattern, while any alignment with the task dynamics space is at chance level. **d:** Schematic of the dominant direction of non-normal amplification in a network obeying Dale’s law. Inputs aligning with the difference pattern (top right singular vector, *v*_1_) are amplified by a factor of 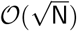 along a co-activation pattern (top left singular vector, *u*_1_). **e:** Schematic illustration of the misalignment across the task-dynamics and task-activity space of the network, where the task-activity space has an alignment with the E/I co-activation pattern and the task-dynamics space does not. Non-normal amplification leads to extra variance being produced along the E/I co-activation pattern. ↑ Go back

The network responses to stimulation input were approximately orthogonal to the task dynamics space, yet the model reproduced the alignment of the stimulation response with the task activity space that was observed in neural data. This response feature underlies general properties of dynamical systems with E/I structure. The dominant pattern of activity in E/I networks represents an amplification from a differential pattern (where E cells are activated above, and I cells below their baseline firing rate) to a co-activation pattern (where E and I cells are both activated above their baseline firing rate).^51^ This can be understood in terms of the singular value decomposition of matrices with Dale’s law sign constraints: The leading singular value of any such connectivity matrix scales as 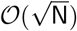, and the leading left and right singular vectors take the form of a co-activation and differential activation pattern, respectively (Fig. 4c). Any input or activity that aligns with the differential pattern will thus be strongly amplified along the co-activation pattern, solely due to the Dale’s law constraints of the connectivity matrix. Indeed, in our trained networks we found that network activity spanned extra dimensions outside of the task dynamics space (Fig. 4b), which aligned with the E/I co-activation pattern (Fig. 4c). As a result, the task activity space of the networks was aligned with the E/I co-activation pattern, even when the task dynamics space was not (Fig. 4e).

Overall, the E/I network model demonstrated that the observed stimulation responses are consistent with the hypothesis that motor cortical dynamics are subspace structured (Fig. 1c, **H2**). Furthermore, the model provided theoretical insight into the circuit mechanisms underlying our empirical measurements. Robustness to additive perturbations arose naturally when low-dimensional dynamical structure was embedded in a high-dimensional network, while non-normal amplification due to E/I cell-type constraints explained the alignment of the stimulation responses with the task activity space.

## Observing intracortical electrical microstimulation during reaching

Having observed that responses to optogenetic perturbation were inconsistent with the reservoir hypothesis (**H1**) but could be reproduced by E/I network models with self-contained subspace dynamics (**H2**), we sought to evaluate the third possibility of path-following dynamics (**H3**). In principle, **H2** could be distinguished from **H3** by examining the effects of perturbation *within* the task dynamics space. Under subspace-structured dynamics, such perturbations would trigger long-lasting neural effects that are shaped by the same dynamical system as the one that governs the evolution of task-related neural activity (Fig. 1f). In contrast, under path-following dynamics, all local perturbations should decay rapidly back towards the externally imposed trajectory (Fig. 1g). However, according to our theoretical analyses, additive optogenetic stimulation directed at local excitatory neurons could be expected to evoke only vanishingly small, random perturbations in the task dynamics space of networks with subspace-structured dynamics. Therefore, **H2** and **H3** cannot be distinguished based on the results from these optogenetic perturbations alone.

We reasoned that the ability to influence task-relevant behavioral outcomes might indicate that an experimental intervention is able to induce perturbations within the task dynamics space. Thus, we turned to intracortical electrical microstimulation (ICMS), which has long been used to probe causal relationships between brain regions and behavior^58,59^ and, when delivered to motor cortex, is known to disrupt motor preparation^60^ and to evoke movements readily.^61^

We trained two additional macaques (monkeys P and V) in the same reaching task, and delivered high-frequency ICMS through a stimulating electrode while recording nearby spiking activity with a Neuropixels probe (Fig. 5a, Extended Data Fig. 6a-c). Like optogenetic stimulation and consistent with prior studies,^60,62^ ICMS in PMd slowed reaction times (Extended Data Fig. 9a). In some sessions, primarily when stimulating in M1, we observed visible deflections of the hand path when ICMS was delivered mid-reach (Peri-Move stimulation, Fig. 5b,c), confirming that ICMS can alter reach kinematics.

**Fig. 5.**
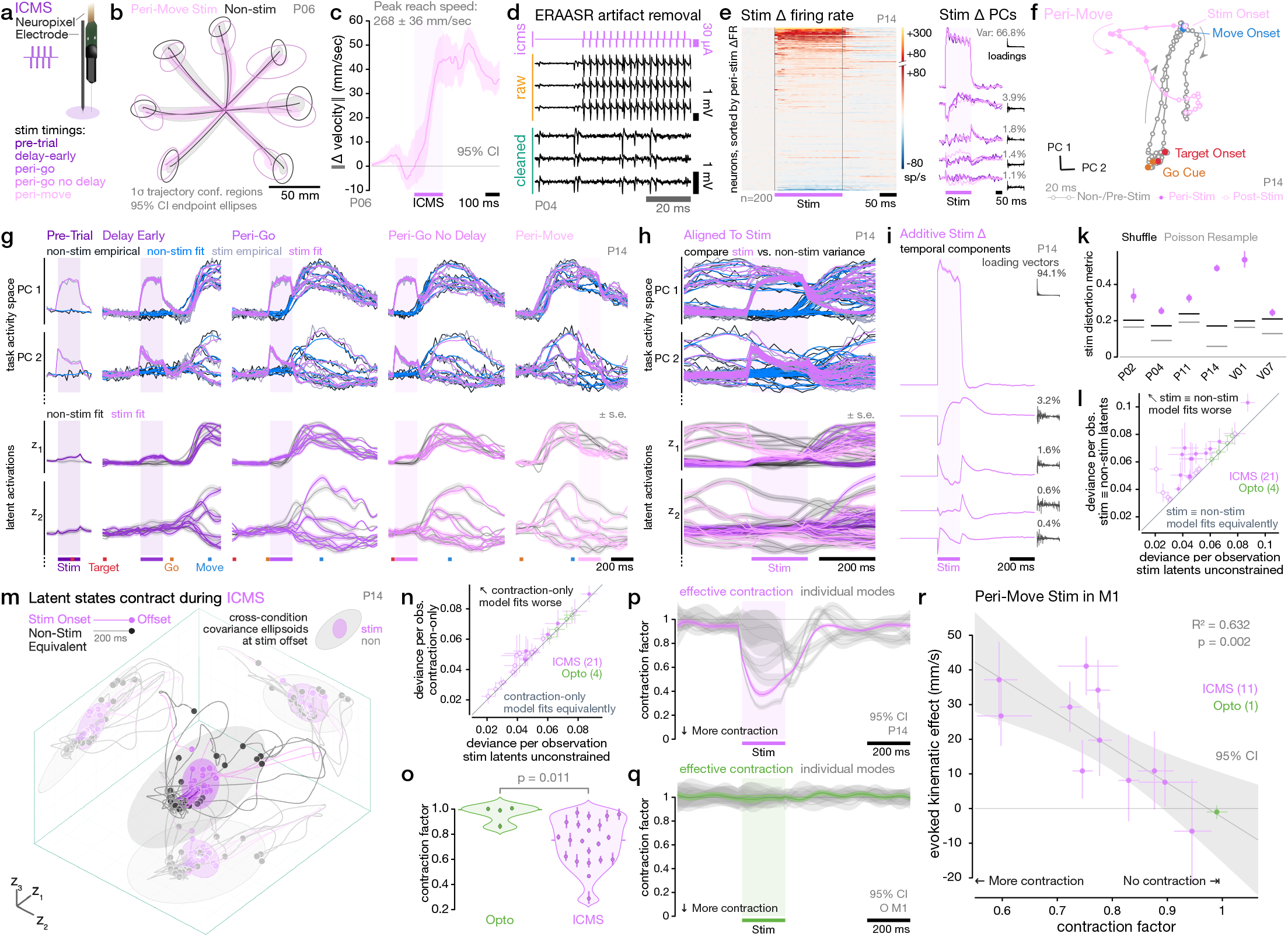
ICMS alters reach kinematics and distorts task-related activity. **a:** Biphasic electrical stimulation was delivered near a Neuropixel electrode. **b:** Reach kinematics for non-stimulated and peri-move stimulated reaches. **c:** Norm of change in hand velocity vector evoked by peri-move stimulation vs. time. **d:** Example of ERAASR_2_ artifact removal on three example channels with visible neural spiking. **e:** Left: Stimulation responses of all recorded neurons at P14 site, sorted by mean change in firing rate (ΔFR) during stimulation period. Right: Principal components of stimulation evoked ΔFR. Timecourses colored by stimulation timing as in (**a**); black stem plots show PC loading vectors over neurons, sorted by ΔPC 1. **f:** State space visualization of stimulation evoked displacements of neural state within the task-activity space (PCs of non-stimulated trial-averages) at site O. M1. **g:** Top: Empirical and LVM-fitted trial-averaged firing rates projected into leading two task activity space dimensions. Stimulation timings are separated into columns, with each trace corresponding to a reach direction. Bottom: Fitted latent activations for the leading two latent dimensions. **h:** Same as (g), aligned to stimulation onset. **i:** Timecourses of the additive delta term which reflects a condition-invariant stimulation induced change in firing rates. Inset: loading vectors mapping each component into neural space together with the percentage of variance each component explains in the empirical stimulation-delta. **k:** Shape similarity metric quantifying distortion between non-stimulated and stimulated latents for six example sessions. Shuffle and Poisson resample lines indicate significance thresholds (*α* = 0.05) under two null hypotheses, see Methods §8.3. **l:** LVM goodness of fit (as Poisson deviance) with stimulation latents fit independently of non-stimulation latnents (horizontal) or constrained to be equal (vertical). Filled circles indicate statistically significant differences (*α* = 0.05); error bars indicate jacknife standard error. **m:** Inferred trajectories of leading three latent dimensions. Circles indicate states at end of stimulation (or equivalent times in non-stimulated conditions). Ellipsoids indicate 90% confidence covariance ellipses across conditions. **n:** LVM goodness of fit (as Poisson deviance) with contraction-only model vs. independent stimulation latents, presented as in (**l**). **o:** Distribution of stimulation-averaged effective contraction factors in each session (shown with 95% CIs) vs. stimulation modality. **p,q:** Contraction factor timecourses for the effective latent variance (colored traces) and for the individual contraction modes in latent space (gray traces). **r:** Relationship across M1 stimulation sessions between evoked kinematic effect, i.e. the integral under curve in (**c**) vs. stimulation-averaged effective contraction factor. Shading shows 95% CIs for a linear fit. ↑ Go back

Observing the neural population response under ICMS presents two unique experimental challenges. First, the effects of ICMS are sensitive to the precise location of the stimulating electrode tip, likely due to the spatially-restricted recruitment of axons.^63^ This precludes a strategy of accumulating responses of the local neural population over multiple sessions with a consistent perturbation. We addressed this by recording with Neuropixels probes,^64^ permitting dense sampling of local population responses on individual experimental sessions. Second, ICMS resulted in very large stimulation artifacts on the Neuropixels probe, which obscured neural spiking activity. We addressed this challenge by developing an artifact removal technique (ERAASR_2_, see Methods §3.2), which exploits differences in the covariance structure of spontaneous neural activity and stimulation artifact to recover spiking activity during ICMS^65^ (Fig. 5d, Extended Data Fig. 6). On some experimental sessions, the electrical artifact transiently saturated the recording amplifier (for *<*1 ms during each biphasic pulse), resulting in undetected spike times during the saturation window. However, spikes in the ∼ 2 ms windows in between ICMS pulses could be accurately detected (Extended Data Fig. 7). Moreover, the pattern of evoked neural firing after each pulse was indistinguishable in sessions with and without saturation (Extended Data Fig. 7g,h). For both non-saturated and saturated sessions, we validated that ERAASR_2_ cleaned neural signals could be used to detect spike times by developing a synthetic stimulation pipeline (Extended Data Fig. 8, Methods §3.6). The synthetic stimulation results confirmed that our approach accurately estimated neural states during and immediately following ICMS and allowed us to compensate conservatively for unobserved spikes during the saturation time window accompanying each stimulation pulse (see Methods §3.7).

## ICMS distorts task-related activity

Like optogenetic excitation, ICMS resulted in large increases in local motor cortical firing rates (Fig. 5e) and displaced neural states within the task activity space (Fig. 5f, Extended Data Fig. 9b-d). Unlike neural responses under optogenetic stimulation, which were fully characterized by an additive response component common across conditions, a non-additive effect of ICMS was visible in the leading principal components comprising the task activity space. Neural firing rates along these dimensions converged closer together (Fig. 5g,h, top two rows, shaded peri-stim regions), resulting in reduced variance across conditions (reach directions × stimulation timings). We applied the same latent variable model (LVM) to quantify this interaction between ICMS and task-related neural activity (Fig. 5g). The LVM successfully captured neural responses during non-stimulated and stimulated conditions. Across all sessions, a large component of neural responses under ICMS could be characterized by an additive transient that is constant across conditions, and that closely resembled the additive component observed in response to optogenetic stimulation (Fig. 5i). However, in some sessions, the LVM revealed that ICMS also altered the underlying latent variables (Fig. 5g,h, bottom two rows), resulting in a significant distortion in the geometry of these states (Fig. 5k, Extended Data Fig. 9e). Constraining paired non-stimulated and stimulated latent variables to take identical values resulted in significantly reduced goodness-of-fit for many sessions (Fig. 5l).

Visualizing the trajectories of the inferred latent variables revealed that ICMS-driven neural states converged towards a common origin across reach directions and stimulation timings during and after stimulation (Fig. 5m). This convergence was also visible in inferred single trial latents (Extended Data Fig. 9f) and in the structure of single trial variability around the condition means (Extended Data Fig. 9g-j). We therefore asked whether this convergence could be described as a strict contraction of the latent states. We fit a modified *contraction-only* version of the LVM in which the latents under stimulation were linked to their non-stimulated counterparts through a pure contraction towards a fitted centroid (Extended Data Fig. 9k-l). Despite having many fewer degrees of freedom, the contraction-only version of the LVM fit the data equivalently well for the majority of the ICMS datasets (Fig. 5n). This demonstrates that the effects of ICMS on task-related activity were well described by a contraction of neural states towards a specific induced state in neural space.

For each session, we computed a contraction factor, defined as the ratio of variance in the stimulated vs. non-stimulated neural states, with one corresponding to no contraction, and zero indicating complete contraction to a point. We validated that the contraction factor could be estimated accurately using simulated transformations of each dataset (Extended Data Fig. 9m, Methods §8.5). The contraction factors across the ICMS sessions were heterogeneous, but were significantly lower on average than the optogenetic stimulation sites (Fig. 5o). The contraction of neural states induced by ICMS was maximal during stimulation, and persisted for hundreds of milliseconds thereafter (Fig. 5p). In contrast, the contraction factor remained near one during and after optogenetic stimulation, consistent with a purely additive effect on neural states (Fig. 5q).

Lastly, we sought to understand the link between evoked changes in neural states and the observed effects on reaching behavior. Additive optogenetic stimulation would likely fail by chance to significantly affect neural states within the low-dimensional task dynamics space in a high dimensional network. In contrast, ICMS could affect the task dynamics space by distorting the underlying geometry of task-related neural states through contraction. If this low-dimensional subspace mediated the effects of ICMS on behavior, this would predict a relationship between the degree of contraction evoked in a particular session and the size of the evoked behavioral effect. Consistent with this prediction, for M1 stimulation sites, we observed a strong negative correlation between the computed contraction factor and the magnitude of the evoked change in hand velocity (Fig. 5r; PMd stimulation sites exhibited a non-significant trend with reduced slope, Extended Data Fig. 9n). This correlation was also evident when contraction factors were computed from the latent states inferred with the original, unconstrained LVM. (Extended Data Fig. 9o,p). In contrast, the length of the perturbation induced displacement vector was not predictive of behavioral impact (Extended Data Fig. 9q).

## Recovery from ICMS follows task dynamics

The neural effects of optogenetic stimulation and ICMS were dominated by a condition-invariant translation of neural states (Fig. 6a, red arrow). In addition to this translation, ICMS also resulted in a contraction of neural states towards a common origin across reach directions and stimulation timings (Fig. 6a, right). If condition-specific differences in neural states are shaped and maintained by low-dimensional subspace-structured dynamics (**H2**), then the ICMS-induced contraction would result in an effective perturbation inside the task dynamics subspace (Fig. 6a, teal arrow). Furthermore, the evolution of neural states within the task dynamics subspace following ICMS should be governed by these same dynamics and therefore be predictable based on estimates of the population dynamics obtained using neural data recorded during normal reaches without stimulation.

**Fig. 6.**
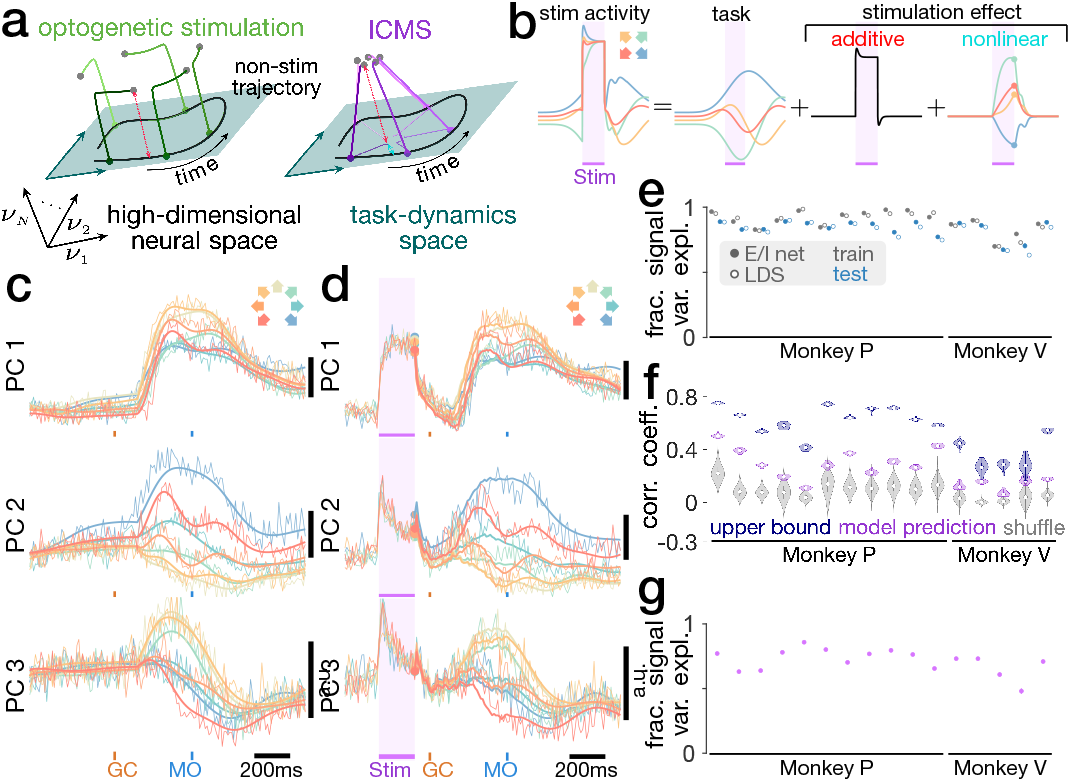
E/I network model predicts population recovery from ICMS. **a:** Schematic illustration of the differences in effects across optogenetic stimulation and ICMS. Optogenetic stimulation can be described as an additive displacement orthogonal to the task dynamics space (red arrow), whereas the distortion of the task-geometry under ICMS additionally results a perturbation inside the task dynamics space (teal arrow). **b:** Schematic illustration of modeling approach for ICMS population responses. PSTHs on stimulation trials are modeled as a combination of normal task-related activity, an additive stimulation response component and a residual stimulation response component. The value of the residual responses component at the end of stimulation is used as an initial condition for modeling the state evolution using the dynamics of the E/I network model (colored dots). **c:** Example model evaluation predicting forward from learned initial condition on training data for session P14. **d:** Example cross-validated model reconstructions of stimulation responses for P14. **e:** The fraction of signal variance explained for training and held out test data for the E/I network model and an analogous low-dimensional LDS model. **f:** An estimate of the state of the residual Δ within the task-dynamics space is obtained by projecting the binned spike times in the last 10 ms of stimulation into the task-dynamics space using the mapping that was learning from non-stimulated data. This projection is used as an initial condition for the dynamics. We show the correlation coefficient between the evolution of residual Δ projected into the task-dynamics space, and the model predictions using the model dynamics for the first 200 ms following stimulation offset, together with the correlation coefficients obtained when predicting the evolution using a version of the dynamics where eigenvalue, eigenvector pairs are shuffled (grey) as well as the theoretical upper bound for a correlation coefficient based on the signal-to-noise ratio of the data. **g:** The cross-validated fraction of signal variance explained by the E/I model reconstructions. Reconstructions are obtained as a sum of the non-stimulated initial conditions and inputs, a random additive perturbation to the E/I network (related to neural recordings via a learned mapping using leave-one-condition-out cross-validation), and using the network dynamics that were learning on non-stimulated data and an inferred initial condition of the residual stimulation to predict the evolution of the residual from stimulation-offset forward. ↑ Go back

To test this, we directly modeled the empirical stimulation effects using the balanced E/I network model with self-contained low-dimensional subspace structure. The stimulation effects can be described as a combination of a condition-invariant, additive component, and a condition-dependent, nonlinear component (Fig. 6b). The additive component represents the condition-invariant translation in neural state space, while the nonlinear component reflects the distortion of task-related structure due to the ICMS-induced contraction of neural states (Fig. 6a,b).

The E/I network model was again fit exclusively to neural activity recorded in the absence of stimulation. The fitted model was able to capture underlying population activity patterns (Fig. 6c) and explained a large fraction of signal variance on non-stimulated trials (Fig. 6e). The additive component of the stimulation effect was modeled as the fitted E/I network’s response to a constant, additive input vector. The elements of this vector were drawn at random from a Gaussian distribution, since ICMS was not targeted to specific cell-types. By chance, this input vector was always approximately orthogonal to the task dynamics space of the network, and thus elicited a rapidly decaying transient response in the network. Next, a random subsample of the network responses to this additive input was used to reconstruct the empirical stimulation effect. The network responses to the additive perturbation were unable to capture the condition-dependent, nonlinear component of the stimulation responses. Thus, the nonlinear stimulation effects could be estimated by subtracting estimates of both task-related activity and additive effects from the recorded neural firing rates (Fig. 6b).

We projected the state of the nonlinear stimulation effect at the last time-point of ICMS into the task dynamics space of our E/I model. Taking this projected state as the initial condition, we then predicted forward in time using the learned dynamics within the task dynamics space and evaluated Pearson’s correlation coefficient between the model prediction and empirical nonlinear stimulation effect throughout the 200ms following the end of stimulation. Strikingly, the model predictions showed a strong correlation with the empirical effect in the majority of datasets (Fig. 6f). The model also performed better than predictions generated by shuffling the eigenvalues and eigenvectors within the task dynamics space, scrambling the fitted dynamics while preserving the characteristic timescales. This demonstrates that the stimulation indeed engaged the task dynamics space in a way that is predictable based on normal task-related dynamical structure, without any further adjustment of model parameters. Lastly, modeling the ICMS responses as a combination of (1) the underlying task-related population responses without stimulation, (2) a response component due to the random additive perturbation input, and (3) the model-predicted evolution of neural states within the task dynamics space starting from an inferred initial condition at the end of stimulation explained a large fraction of signal variance in the stimulated PSTHs (Fig. 6d,g). More details are provided in the Methods §15.

Overall, our analyses demonstrated that population responses to ICMS also reflect structure that is expected in balanced E/I networks with self-contained, low-dimensional subspace structure. Since ICMS induced a contraction of neural states towards a common induced state, this resulted in an effective perturbation inside the task dynamics space (Fig. 6a). The E/I network model fit exclusively to neural activity from trials without stimulation was able to predict the trajectory of perturbed neural states within the task dynamics space. This indicates that these fitted task dynamics act upon neural states at locations far from the trajectories followed during non-stimulated trials. The observation of long-lasting, predictable effects following ICMS constitutes strong evidence against path-following dynamics (**H3**). Instead, neural population responses to both optogenetic stimulation and ICMS were indicative of subspace structured dynamics embedded within an E/I network (**H2**).

### Discussion

The dynamical systems perspective has played a crucial role in understanding the complex neural activity patterns that underlie skilled behavior.^1,3^ However, the precise nature of the dynamical rules which govern the evolution of neural population activity have remained elusive. In this work, we delivered optogenetic and electrical stimulation to perturb the motor cortex while recording the responses of the local neural population. We demonstrated that within the motor cortex, task-related dynamical sensitivity is constrained to a self-contained, low-dimensional subspace of the ambient high-dimensional neural circuit. Self-contained structure allows task-related dynamics to generate low-dimensional patterns of activity, while remaining robust to irrelevant signals or noise outside of the low-dimensional subspace. Moreover, multiple mutually orthogonal self-contained subspaces may be maintained to implement different tasks without interference,^66,67^ or engaged concomitantly to compose multiple dynamical motifs to address task demands.^68^ Thus, the dynamical structure we identified in the context of the cortical control of arm movement may reflect a more general principle of robust cortical computation in the presence of noise or multiple tasks.

Motor cortex receives rich inputs from thalamus and other cortical regions and projects outputs that shape the activity of lower motor centers and coordinate predictive control via the cerebellum.^69^ Ultimately, it is the collective action of this distributed network that is responsible for preparing and executing movement. As a critical node within these circuits, the population dynamics that shape motor cortical activity reflect these recurrent influences. These inputs both shape the dynamical landscape within motor cortex and convey online feedback about the state of the body and the environment.^70–72^ Our work here speaks to the structure of the functional dynamical system that shapes motor cortical activity supporting reaching movements. In particular, the dynamics we observe and perturb likely reflect the functional interplay of motor cortex and strongly bidirectionally connected motor thalamus.^4,73^ Recent studies have begun to probe the role of specific, inter-areal projections in establishing goal-directed movement.^74–76^ This research aims to provide a circuit-level of understanding of the distributed dynamical system—how population activity is shaped both by local synaptic interactions and inputs via specific projections within the network.

The low-dimensional dynamical structure we observed has important implications for probing causal links between brain circuits and behavior.^77,78^ For motor cortical stimulation, evoked changes in reach kinematics were observed only when stimulation affected neural states within the task dynamics space. This was achieved in varying degrees across ICMS sessions, although never with optogenetic excitation. We found that optogenetic stimulation responses were well described as additive, simply translating neural states while preserving their relative geometry. Optogenetic perturbations thereby failed to alter ongoing neural activity and reach kinematics due to the low probability of aligning with the task dynamics space by chance. In contrast, although neural activity during ICMS also contained this additive component, neural states were contracted together during electrical stimulation. This contraction necessarily altered neural states within the task dynamics space, despite not being specifically targeted along those dimensions. The observed contraction towards an induced neural state also explains prior behavioral findings that ICMS supplants rather than adds with task-related muscle commands^79^ and drives the arm towards a consistent final posture.^61,80,81^ During high-frequency ICMS, this replacement effect might stem from collisions within activated axons between evoked, antidromic spikes and task-related signals.^82^ This effective engagement with motor cortical state may also reflect more effective recruitment of thalamocortical projection axons.

It is important to note that many parameters may influence the effect of optogenetic stimulation; driving purely additive effects on firing rates is likely a special case. Our proposed dynamical mechanism is consistent with observed behavioral effects across a broad range of stimulation modalities and experimental contexts. Optogenetic and pharmacological inhibition contract neural states towards the quiescent state.^83–85^ Alternatively, strong optogenetic activation^86^ may entrain neurons near the upper limit of their dynamic range, thereby contracting firing rates towards a distinct induced state. Moreover, targeted stimulation methods might alter behavior by directly impacting the task dynamics space. Perturbations might preferentially engage the task dynamics space when targeting specific projection pathways,^87,88^ cell types,^89,90^ or cell ensembles^91^ with important computational roles in the circuit.^26^ In some behavioral contexts, such as a sensory detection task, untargeted stimulation might alter behavior via broadly sensitized downstream circuitry.^92,93^

The non-normal dynamical structure of E/I networks results in different input signals driving transiently amplified responses along the same E/I co-activation mode of the network (see Methods §14). Variance along this co-activation mode thus arises easily in the presence of external inputs or intrinsic noise. Yet, when excitation and inhibition are balanced, this co-activation mode is dynamically short-lived. Our network models supported the hypothesis that slow and structured dynamics may be embedded in a low-dimensional subspace oriented approximately orthogonal to the E/I co-activation mode and yet contributing variance along this mode. This arrangement provides a mechanism for robust computation in the presence of transient amplification of inputs, noise, or other perturbations in the network. Consequently, computing with self-contained, subspace structured dynamics may be a computationally advantageous strategy employed by other cortical circuits as well. The apparent non-normal structure of cortical circuits also has implications for the analysis and interpretation of neural activity patterns. For example, a high-variance, condition-invariant signal (CIS) that generally aligns with the co-activation mode emerges in motor cortical populations immediately preceding movements, and has been hypothesized to trigger a transition from preparatory to movement-related neural activity.^32,94^ Based on our theoretical analyses, we propose a revised hypothesis: when external inputs arrive to motor cortex to initiate movement, the CIS emerges as a mixture of this dynamical transition into movement and the highly amplified response that emerges along the co-activation mode in tandem. In this view, the CIS itself does not reflect only the trigger signal, but also variance created in response to it. Consistent with this hypothesis, optogenetic perturbations (and some ICMS perturbations) drove neural responses similar to the CIS but failed to trigger movement. Similar considerations apply for any approach that seeks to optimize explained neural variance, including targeted dimensionality reduction.

Non-normal dynamics induce a misalignment between task activity spaces with high variance and task dynamics spaces with long-lived effects. In light of this distinction, experimental perturbations, internal perturbations in the form of trial-to-trial variability,^95^ and dynamical models constrained by neural circuit architecture will be crucial to establish the precise mechanisms of neural computation.

## Author contributions

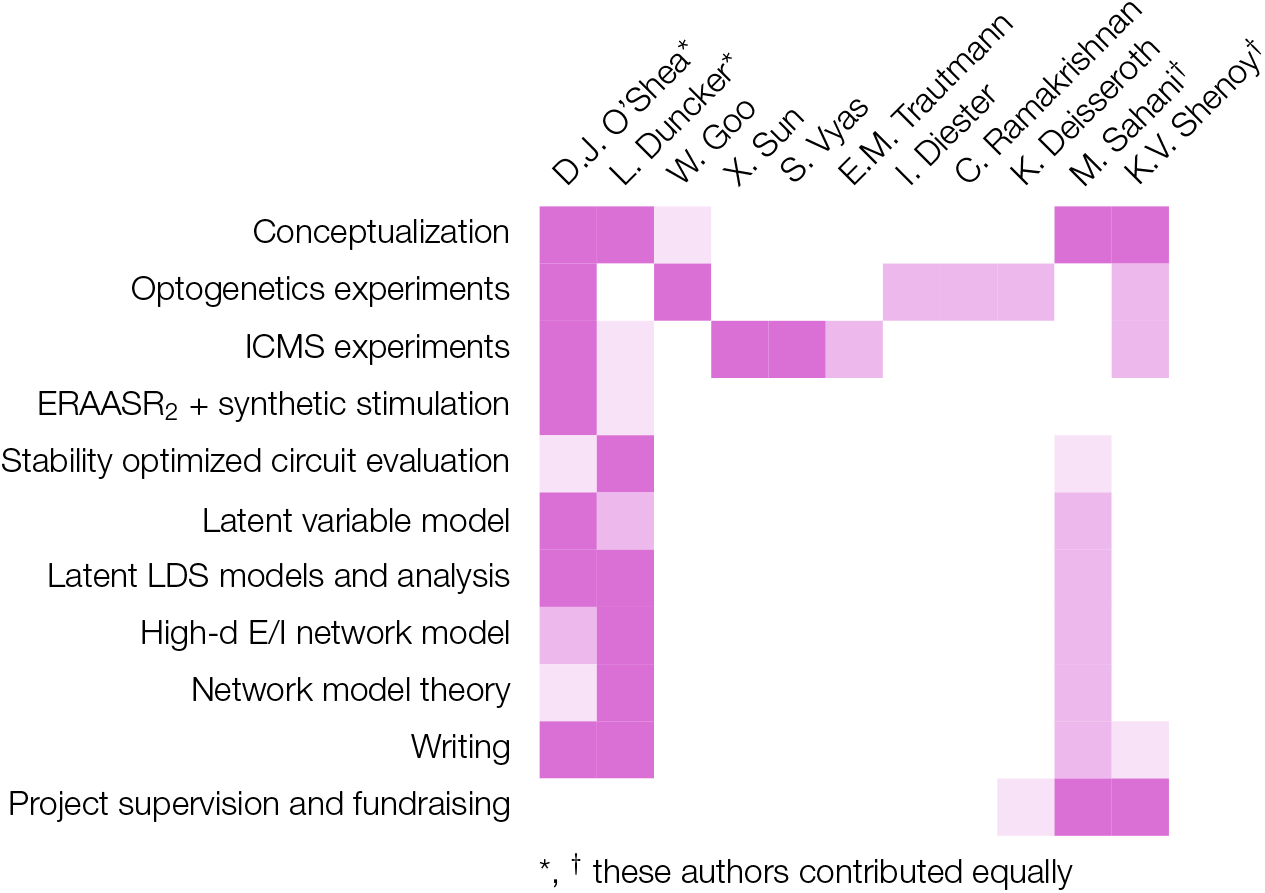

## Acknowledgements

We thank M. Kaufman and C. Brody for formative discussions that profoundly shaped the direction of this research. We thank L. Driscoll and other members of the Shenoy laboratory at Stanford University for comments on the manuscript and discussions on the methods and results; A. Nurmikko, I. Ozden, and J. Wang for graciously providing the coaxial optrodes that were essential to the optognetics experiments; S. Ryu for surgical expertise and advice; M. Risch, M. Wechsler, R. Reeder, J. Aguayo, L. Yates, R. Steinbach, S. Smith, C. Sherman, E. Morgan, W. Kalkus, S. Kang, C. Pacharinsak, and M. Huss for expert veterinary care; and B. Davis and E. Castaneda for administrative support.

This work was supported by a National Science Foundation graduate research fellowship (D.J.O.) ^[2009079313]^; the Dr. Regina Casper Stanford Graduate Fellowship (D.J.O.); the Gatsby Charitable Foundation (L.D., M.S.); the Simons Foundation Collaboration on the Global Brain (D.J.O., L.D., M.S., K.V.S.) ^[325380, 543045]^; the Human Frontier Science Program fellowship (I.D.); the German Academic Exchange Service Award (I.D.); the Defense Advanced Research Projects Agency (DARPA) Biological Technology Office (BTO) REPAIR award (K.D., K.V.S.) ^[N66001-10-C-2010]^ and NeuroFAST award (K.D., K.V.S.) ^[W911NF-14-2-0013]^; the NIH Director’s Pioneer Award (K.V.S.) ^[8DP1HD075623]^; the Office of Naval Research (K.V.S.) ^[N000141812158]^; the Hong Seh and Vivian W. M. Lim endowed chair (K.V.S.); and the Howard Hughes Medical Institute at Stanford University (K.D., K.V.S.).

## Methods

### 1 Experimental model and subject details

All surgical and animal care procedures were performed in accordance with National Institutes of Health guidelines and were approved by the Stanford University Institutional Animal Care and Use Committee. The subjects were four adult male macaque monkeys (*Macaca mulatta*): O (13 kg, 7 years old), Q (10 kg, 5 years old) P (16 kg, 11 years old), V (9 kg, 9 years old). After initial training, we performed a sterile surgery during which each macaque was implanted with a head restraint and either a 19 mm circular recording cylinder (O, Q, and V; Crist Instruments) or a 30 mm long oval recording cylinder (P; NAN Instruments). The recording chamber was located over left, caudal, dorsal premotor cortex (PMd) and primary motor cortex (M1), placed surface normal to the skull, and secured with methyl methacrylate. In monkey Q, a craniotomy was performed which exposed dura within the full extent of the recording chamber. In monkeys O, P, and V, a thin layer of methyl was deposited atop the intact, exposed skull within the chamber. and a set of small (3 mm diameter) craniotomies was performed near the center of the chamber to facilitate injections and/or electrical stimulation and recording.

We confirmed targeting of M1 / PMd by testing for neural modulation during movements and palpation of the upper arm. We also confirmed that suprathreshold electrical stimulation evoked twitches of the shoulder and arm musculature.

### 2 Optogenetic experiments

#### 2.1 Injections

↪ c.f. Fig. 2a

In monkey O, two injections of AAV5-CaMKII*α*::C1V1(E122T/E162T)-ts-EYFP (3 × 10^12^ viral genomes/mL) were made within each of seven mini-craniotomies. These injections spanned the arm regions of both M1 and PMd. In Q, a similar grid pattern of 13 injections were made approximately 2.5 mm apart spanning M1 and PMd. All viral vectors were pressure injected with an injection assembly as reported previously.^45^ At each injection site, we injected 1 μL of the viral vector at depths located 1 mm apart spanning cortex. The infusion rate was 0.1 μL/min. In monkey Q, we confirmed histologically that strong opsin expression was present throughout all cortical layers in the injected regions.

#### 2.2 Stimulation and recording

↪ c.f. Fig. 2a,b

In early experiments with Q, we delivered optical stimulation with a 200 μm flat cut fiber optic, secured along its length to a tungsten microelectrode (FHC), as described previously.^45^ In later experiments with Q and all experiments with O, we used a custom-manufactured coaxial optrode in which a central light guide was surrounded by a gold recording surface and insulating shell.^44^ Both optrode types enabled simultaneous light delivery and electrical recording, and were lowered into position using a motorized microdrive (NAN Instruments). Adjacent to the optrode, we also lowered 1–3 tungsten microelectrodes (typically 3), at lateral distances ranging from 500 μm to 6 mm. Electrodes were lowered via independent microdrives in parallel with the optrode. This arrangement allowed us to sample many neurons in the local neighborhood around the optrode both within and across days while keeping the optrode (the site of stimulation) in the same location. This strategy served both to minimize damage to the cortex due to the optrode, as well as to provide a constant perturbation that could be sampled by the individual electrodes in the surrounding neural population. Defined by the position of the optrode, we stimulated at three unique sites in monkey O, in gyral M1 (O. M1), in posterior PMd (O. PMdP), anterior PMd (O. PMdA) and at one site in PMd in monkey Q (Q. PMd). We recorded the locations of the electrode and optrode and validated their relative positions by analyzing photographs of the guide tube ends taken from directly below the microdrive assembly after each experimental session. We used these coordinates to compute the distance from the optrode for each unit recorded on a surrounding electrode.

Optrode and electrode voltages were buffered and digitally amplified at a head-stage, and recorded with the Cerebus neural signal processor (Blackrock Microsystems). Broadband signals were recorded on each channel and filtered at the amplifier (0.3 Hz one pole high-pass filter, 7.5 kHz three pole low-pass filter). The signals were also digitized to 16-bit resolution over ±8.196 mV (resolution = 0.25 μV) and sampled at 30 kHz. Each channel was differentially amplified relative to a local reference, typically a guide tube flush with the dural surface. To aid with noise rejection, the electrodes and head stage were surrounded with a flexible electromagnetic shielding mesh fabric (McMaster Carr), which was also shorted to the guide tube. This same shielding also served to block light emitted from the optrode from illuminating the surrounding rig. Voltage signals were band-pass filtered (250 Hz–7.5 KHz) and thresholded at -3.5 times the root-mean-square voltage. Spiking waveforms were later hand-sorted (Plexon Offline Sorter). We collected 205 neurons at O. M1, 246 at O. PMdA, 221 at O. PMdP, and 244 at Q. PMd.

Optical stimulation was computer controlled by the real time experiment controller running TEMPO (Reflective Computing), which triggered a Master 8 pulse generator (A.M.P.I.) and in turn controlled a green DPSS laser (561 nm, CrystaLasers) whose power output at the optrode was manually adjusted to 3 mW output at the start of each experimental session. Stimulation pulses were 200 ms continuous pulses at 95 mW/mm^2^ (3 mW total power), which we had previously observed to generate the largest increases in firing rates at the optrode, relative to high-frequency light pulse trains, in a separate set of experiments.

Each experimental day consisted of multiple blocks, each consisting of 300-600 trials. Within each block, the electrodes remained stationary and only units and multi-units which could be reliably isolated for the duration of the block were analyzed. Between blocks, we advanced the electrodes independently to isolate new units before resuming stimulation.

### 3 ICMS experiments

#### 3.1 Stimulation and recording

↪ c.f. Fig. 5a, Extended Data Fig. 6a

In monkeys P and V, several mini-craniotomies were made within the recording chamber. The stimulation electrode was a single tungsten microelectrode with approximately 100 kΩ impedance at 1 kHz (Configuration #UEWLGC-SECN1E, Frederick Haer Company). Neural population responses were recorded using a Neuropixels phase 3a probe (IMEC) with 374 recording channels and an external reference. The stimulation electrode, the Neuropixels recording probe, and a third blunt guide tube (used for dural surface stabliization) were secured to independently controllable, motorized micromanipulators (NAN Instruments). To insert the Neuropixels probe, we made a short linear incision through the dura. On some sessions, we also inserted a tungsten microelectrode a short distance beyond the pial surface to create a pilot hole to facilitate the Neuropixels probe insertion. The Neuropixels probe was inserted as close as possible to the stimulation electrode while preventing collision with the headstage, typically between 2–6 mm away.

The stimulation was performed using a StimPulse electrical stimulation system used as a combined function generator and isolated current source (Frederick Haer Company). Microstimulation trains consisted of biphasic pulses delivered at 333 Hz (150 μs cathodic, 100 μs pause, 150 μs anodic). Stimulation duration was most commonly 200 ms to match the optogenetic parameters, although some sessions 60 ms or 800 ms stimulation durations were used. For 800 ms stimulation sessions, subsequent analyses focused on the initial 200 ms of stimulation. Stimulation amplitude was varied between 20–160 μA across sessions.

We recorded continuously through stimulation using the Neuropixels. On each session, we balanced the competing demands of eliciting a behavioral effect with sufficient stimulation amplitude, avoiding saturation, and using AP-band gain sufficiently large to detect neural spiking, and recording near the stimulation site. Variation in the choices of stimulation amplitude, stimulating electrode and Neuropixels probe location, and AP gain led to heterogeneity in evoked behavioral effects across sessions. We collected Neuropixels data at 11 unique recording sites, and then defined an experimental session as contiguous group of trials with a unique set of stimulation electrode locations and stimulation amplitude and duration. We collected data for 6 sessions without saturation in the recorded Neuropixels signals and 15 sessions with partial or complete saturation during each individual current pulse.

#### 3.2 Electrical artifact removal with ERAASR_2_

↪ c.f. Fig. 5d, Extended Data Fig. 6d-f, Extended Data Fig. 7a-b

First, individual AP band data files were digitally concatenated after scaling the files to a common AP gain. Individual recording channels were excluded from further analysis if they exhibited atypical RMS (outside of 3–100 μV) or exhibited a rapidly toggling baseline (resulting from a process flaw in the phase 3a probes) which resulted in a bimodal voltage histogram. Every channel was subsequently manually reviewed.

We then scanned each stimulation period to detect saturation for each individual channel. On the phase 3a probes, the saturation limits were not matched to the limits of the quantization (e.g., ± 2^9^ samples). Consequently, the saturation limits were determined automatically for each channel (and if multiple files were concatenated, for each AP gain setting) individually. We identified the minimum and maximum value of each channel over the recording (with constant AP gain) as well as the baseline offset of each ADC group. The range of sample values, relative to baseline, observed on each ADC group was then used to define conservative signal ranges for each channel. Computation of these ranges was aided by one or more short duration, test stimulation periods recorded before the start of the experiment conducted with higher amplitudes than those used in the actual experiment. Timepoints where the recorded voltage approached the limits of these per-channel signal ranges were marked as possibly saturated in subsequent signal processing.

To remove the artifact, we exploited the difference in covariance structure between neural activity, which is spatially localized on the probe, and artifact, which is spatially broad and relatively homogeneous. We refer to the following algorithm as ERAASR_2_, a modified version of the original algorithm published in O’Shea and Shenoy [65]. For each trial independently, we first computed the spontaneous covariance matrix across channels Σ^spont^ ∈ ℝ^*C*×*C*^, where *C* is the number of usable recording channels (typically 350–374). For this, we used 1 second of data without any stimulation immediately prior to stimulation. During the subsequent stimulation artifact period, we computed the artifact covariance matrix as Σ^art^ = X^art^(X^art^)^⊤^, where X^art^ ∈ ℝ^*C*×*T*^ is the centered data matrix (channels by time) during the artifact, excluding samples marked as possibly saturated.

We then identified a low-dimensional subspace which explained maximal variance during the artifact period, while capturing minimal spontaneous neural variance, by solving the optimization problem:

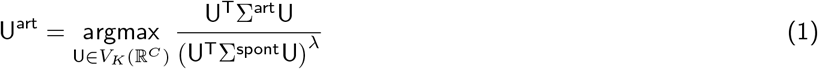

where *V*_*k*_(ℝ^*C*^) is the Stiefel manifold of *K* dimensional subspaces in *C* dimensions, and *λ* is a hyperparameter that controls the tradeoff between capturing the artifact variance (the numerator) and avoiding spontaneous neural variance (the denominator). With *λ* = 0, this reduces to PCA on the artifact directly. We used *λ* = 0.05.

With this subspace, we then proceeded as in the original ERAASR algorithm with a version of principal components regression, using U^art^ instead of the principal component dimensions. We projected the artifact data into this subspace to obtain a set of *K* artifact components as A = (U^art^)^⊤^X^art^ ∈ ℝ^*K*×*T*^. Then, for each channel individually, we regressed that channel’s data onto the artifact components (again excluding possibly saturated samples in determining the coefficients). Lastly, we subtracted the predicted artifact on this channel from the recorded signal to obtain the cleaned channel data. For each channel *c*, we have

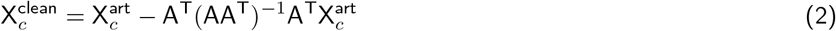

When many channels were marked as possibly saturated (more than 60%), we considered all channels to be saturated, and cleared the values of every channel to zero during this peri-pulse saturation window. Otherwise, for each sample which was marked possibly saturated, we checked whether the absolute value of the predicted artifact exceeds the absolute value of the recorded data, i.e. if 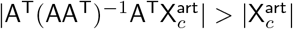, and mark these timepoints as actually saturated, and clear their values to zero. Unlike in the original ERAASR algorithm, we found it unnecessary to clean the resulting data across pulses within the train or across trials.

We reviewed the cleaned neural signals manually for each artifact. The entire data file was then common averaged reference for each ADC group separately, as described in Siegle et al. [96], excluding samples marked as saturated, which remained set to zero.

#### 3.3 Location estimation from artifact amplitude

↪ c.f. Extended Data Fig. 6b

Having estimated the artifact for each stimulation event, we computed the peak to peak artifact amplitude on each channel. We removed outliers using a moving median method, then resampled the artifact amplitude estimates to a vertical column at the center of the probe using natural neighbor interpolation. We then fit these amplitude measurements collected across all sessions to a model relating amplitude to the distance of each site to the stimulation source. We defined *x*_*s*_ as the distance measured laterally from the probe to the stimulating electrode on session *s*, and *y*_*s*_ as the vertical distance along the penetration path from the most superficial electrode to the tip of the stimulating electrode. Then each location *e* where the artifact amplitude is resampled is located vertically at *y*_*e,s*_ = *y*_*s*_ + *eδ*_*y*_, where *δ*_*y*_ = 50 μm is the spacing between samples. This resample location is then located at a distance

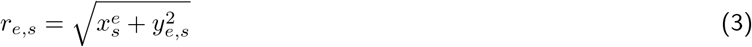

from the stimulation source, defined to lie at the origin. The electric field falls off with distance, such that the amplitude of the artifact at each resampled location, given stimulation amplitude *I*_*s*_ follows:

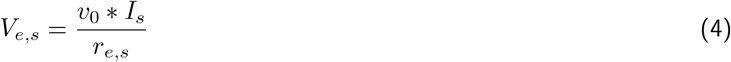

where *v*_0_ is a fitted parameter with units V mm/μA. We fit this model for all sessions simultaneosly using a constrained nonlinear curve fitting algorithm to minimize the squared error between empirical and predicted artifact amplitudes. At each recording site, we use the minimal stimulation amplitude used (which may be lower than that used for the subsequent experiment). The model was first fit with non-saturated sessions to fit *v*_0_, and then subsequently fit again with the saturated sesssions with fixed *v*_0_. For sessions where the artifact saturated the majority of the channels, the fitted distance *x*_*s*_ constitutes an upper bound on the distance—if the Neuropixels probe were further away than the modeled distance, the artifacts would be smaller and therefore not saturate the amplifier.

#### 3.4 Spike sorting

Cleaned AP band data were automatically spike sorted with a modified version of Kilosort2 spike sorting software.^97^ The modifications do not affect the core operation of the algorithm; instead, these changes facilitate subsequent processing for the synthetic stimulation pipeline. First, we made several changes to ensure the outputs of Kilosort2 are fully reproducible. This was accomplished by preventing race conditions in the GPU kernels by using atomic functions where appropriate, sorting certain arrays to ensure determinstic ordering, and rounding floating point operations at sufficient levels of precision to prevent issues with non-associative addition. Second, Kilosort2 processes data in time batches and allows waveform templates to drift over time. We track and persist these batchwise parameters, allowing us to resort individual batches of data using the identical parameters and templates used as in the original full-dataset sorting process. Lastly, we ignore any time windows during stimulation in specific algorithm steps where RMS or covariance is computed and where the waveform templates are updated over batches. This reproducible version of Kilosort2 is available at https://github.com/djoshea/Kilosort2. Every sorted neuron was manually reviewed using a custom GUI written in Matlab App Designer (Mathworks).

#### 3.5 Multichannel spike waveform evaluation

↪ c.f. Extended Data Fig. 6g-j, Extended Data Fig. 7c-f

For each sorted neuron, we assembled the set of spiking waveforms during spontaneous, non-stimulated periods, during the stimulation window, and in the 300 ms period immediately post-stimulation, and computed the average spiking waveform for each group. We identified the channel with the largest spiking waveform for each neuron and the six immediately adjacent channels. To quantify the amount of distortion present in the sorted waveforms that remained after artifact removal, we computed the correlation coefficient between the waveforms on these seven channels, comparing spontaneous with peri-stim and spontaneous with post-stim.

#### 3.6 Synthetic stimulation pipeline

↪ c.f. Extended Data Fig. 8

For both non-saturated and saturated sessions, we validated that ERAASR_2_ cleaned neural signals could be used to detect spike times by developing a synthetic stimulation pipeline (Extended Data Fig. 8). Here, we corrupted neural signals from paired non-stimulated trials with stimulation artifacts estimated from true stimulation trials before applying the entire ERAASR_2_ and spike sorting pipeline. This synthetic stimulation pipeline enabled us to validate our approach against known ground truth data.

For each stimulation condition (reach direction × stimulation time), we identified otherwise equivalent time periods in matched non-stimulated trials for use as ground truth. We extracted the raw Neuropixel AP band voltage traces from these nonstimulated trials and added to them the stimulation artifact estimated on the corresponding stimulation trials, yielding what we define as synthetic stimulation traces. Where a given channel was marked as saturated, we replicated this saturation by clipping the synthetic stimulation traces at the known saturation limits of each channel. We reinserted these synthetic stimulation traces into the AP band data. As with the true stimulation data, we estimated and removed the artifact using ERAASR_2_. We then compared the ground truth non-stimulated voltage traces against the synthetically-corrupted and cleaned traces by computing the correlation coefficient.

We then used our modified Kilosort2 to resort these batches of data, restarting the algorithm using the exact state and batchwise waveform templates that were used to sort these batches of data during the original sorting pass. We subsequently reapplied the same cluster thresholding, splitting, and merging steps performed by both the algorithm and during manual curation. This allowed us to compare the synthetic stimulation spike trains against the ground truth extracted from these non-stimulated traces during the original spike sorting procedure. For neuron and for each true non-stimulation spike, we marked the spike as *recovered* if a corresponding synthetic stimulation spike was marked within 500 μs (true positive) or marked as *missed* if no corresponding synthetic spike was found (false negative). Similarly, for each detected synthetic stimulation spike, we marked the spike as *fabricated* if no corresponding ground truth spike was found for that neuron within 500 μs (false positive). The fraction of recovered ground truth spikes (ideally 1) and of fabricated detected spikes (ideally 0) served as a stringent end-to-end evaluation of the artifact removal pipeline in terms of actual detected spiking activity.

#### 3.7 Firing rate gain correction

↪ c.f. Extended Data Fig. 7g-h, Extended Data Fig. 8a, i–k, n–p

For each neuron *d*, timepoint *t*, we next computed the average firing rate within the stimulation period for synthetic stimulation 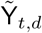 versus the ground truth rate Y_*t,d*_, averaged over all trials across synthetic stimulation conditions. For both saturated and non-saturated sessions, these firing rates were tightly correlated Extended Data Fig. 8j,o, indicating that neural states were estimated accurately during (and immediately following) ICMS. For saturated sessions, the recovered synthetic firing rates were proportionally reduced relative to ground truth, due to the loss of spiking signals during the saturation window. We computed this recovery gain as the element-wise ratio 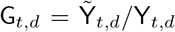. We denoised this gain matrix by computing a rank one approximation of the gain matrix around unity, i.e. 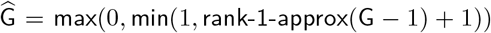. This denoised gain matrix represents the estimated fraction of true spikes detected for each neuron, in each time bin, due to saturation or distortion resulting from the ERAASR_2_ cleaning pipeline. We corrected for this distortion by dividing out the respective gain, i.e. 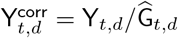.

Therefore, the synthetic stimulation pipeline allowed us to compute and compensate for the unobserved spikes during the saturated timepoints accompanying each stimulation pulse. We applied this gain correction to both the stimulated firing rates observed during ICMS and to the paired non-stimulated firing rates, which were taken from the output of the synthetic stimulation pipeline with the identical gain correction applied. Consequently, for all sessions, for all comparisons between non-stimulated and stimulated firing rates, an identical sequence of processing steps (ERAASR_2_ artifact removal, spike sorting, firing rate gain correction) was applied. This ensures that any distortion created by the artifact removal process itself would affect non-stimulated and stimulated firing rates equally. Critically, the gain correction step applies identically to both nonstimulated and stimulated firing rates and therefore does not affect the latent variable model and contraction analyses described in §8.

With respect to the gain correction, we note that this synthetic stimulation pipeline is also conservative with respect to the observed contraction of neural states. Our reasoning is as follows:

> Suppose that we consider a single ICMS pulse, and consider two time bins, where the first time bin is fully saturated over channels, whereas no channels are saturated in the second time bin. Consider if we apply the synthetic stimulation pipeline to non-stimulated data where a given neuron fires 10 spikes/sec in both time bins, i.e. {10, 10}. Due to the saturation, suppose we recover 0 spikes/sec in the first bin, and 10 spikes/sec in the second time bin, i.e. {0, 10}, for an average rate of 5 spikes/sec. We then compute a gain of 5 / 10 = 0.5, which then applied to the recovered rate of 5, yields the correct 10 spikes/sec. In the true stimulation condition, consider that ICMS raises firing rates to 20 spikes/sec in the first bin and 30 spikes/sec in the second bin, i.e. {20, 30} for a true average rate of 25 spikes/sec. Due to saturation, we observe {0, 30} for an average of 15 spikes/sec, which is then corrected to 15 / 0.5 = 30 spikes/sec, leading us to overestimate the true firing rate (25 spikes/sec) averaged over the full stimulation period.

The gain correction implicitly assumes that spikes are nearly uniformly distributed throughout the stimulation period, such that the firing rate can simply be linearly scaled to compensate for the fraction of samples where spikes are lost to saturation. However, in the non-saturated sessions, we observed that ICMS generally evokes more spikes in the period of time between the pulses (Extended Data Fig. 7g). In assuming that the firing rate during the saturated window during each ICMS pulse matches the observed rates immediately following each pulse, the gain correction multiplier is likely too large for the true stimulation rates. This in turn tends to *underestimate* any contraction of neural states during ICMS when saturation is present.

We also developed a more complex gain correction mechanism that used regression to incorporate fine temporal dynamics of the ICMS evoked firing rates, at the resolution of the raw sampling rate (30 kHz). These estimates (data not shown) corrected for the non-uniformity of evoked spiking with respect to each pulse, and yielded nearly identical results in all subsequent analyses, including those related to the observed contraction.

### 4 Task design

During training and experiments, monkeys sat in a customized primate chair (Crist Instruments) with an opening to allow the arms to move freely. The head was restrained via an implant and the left arm was held comfortably in place using a tube and cloth sling. Stimuli were presented on a screen in the frontoparallel plane located approximately 25 cm from the eyes. A photodiode was used to record the timing of stimulus presentation on screen with 1 ms resolution. A clear, acrylic, removable visor shielded the neural recording system and prevented the monkey from bringing the reflector bead to his mouth. A flexible tube connected to a fluid-flow valve solenoid and attached to the visor dispensed juice rewards.

Monkeys were trained to perform a center-out delayed reaching task. Each experimental session consisted of several thousands of trials, ending with a juice reward if successful and a short (typically 500-1200 ms) time penalty if unsuccessful. Each trial begins when the monkey touches a central touch target presented at approximately eye level on the screen. After a short, variable hold time (200-600 ms), a reach target is presented (Target Onset) at a distance of 10 cm from the central target in one of a set of discrete radial directions. On delay trials, the target initially appeared hollow and its position jittered slightly around the actual location. Cessation of the target jittering, filling of the target, and disappearance of the central hold target provided the Go Cue signaling that the monkey could initiate his reaching movement. The Go Cue followed Target Onset after 0-800 ms delay period. On zero delay trials, the target was presented as a stationary filled shape at the same time that the central target disappeared, which signaled the simultaneous Target Onset and Go Cue. Both the timing of Target Onset and Go Cue were recorded using the photodiode. Online movement onset was detected using position boundaries and speed thresholds, with the trial terminating if the online reaction time was too rapid suggesting pre-empting of the Go Cue (e.g., under 150 ms) or too slow (e.g., longer than 500 ms). Reaches were rewarded if they were brisk and terminated accurately within the target.

#### Optogenetics experiments

The reaching task was performed by touching and holding targets on a screen. Hand position was measured at 60 samples/sec by tracking the position of a reflector bead taped to the middle finger of the right hand using an infrared stereo tracking system (Polaris; Northern Digital, Waterloo, Ontario, Canada). Four reach directions were used (45, 135, 225, and 315 degrees from the positive x-axis).

#### Electrical stimulation experiments

We trained the monkey to use his right hand to grasp and translate a custom 3D printed handle (Shapeways, Inc.) attached to a haptic manipulandum (Delta.3, Force Dimension, Inc.). The other arm was comfortably restrained at the monkey’s side.Seven reach directions were used (0, 45, 90, 135, 180, 225, and 315 degrees counter-clockwise from the positive x-axis).

The device reports its position in 3D space via optical encoders located on its 3 motor axes and can render 3D forces by applying torques to these motors in the appropriate pattern. The haptic device was controlled via a 4 kHz feedback loop implemented in custom software (https://github.com/djoshea/haptic-control) written in C++ atop Chai3D (http://chai3d.org). The weight of the device was compensated by upward force precisely applied by the device’s motors, such that the motion of the device felt nearly effortless because the device’s mechanical components were lightweight and had low inertia. The device endpoint with the attached monkey handle was constrained via software control to translate freely in the frontoparallel plane. Aside from the stiff, planar spring-like forces that kept the endpoint within this plane, no forces were applied to the device during the task.

The handle was custom 3D printed and contained a beam break detector which indicated whether the monkey was gripping the handle. The handle also contained a 6 axis force / torque transducing load cell (ATI Instruments). Hand position was recorded at 1 kHz, and the X/Y position of the device was used to update the position of a white circular cursor at the refresh rate of 144 Hz with a latency of 2-8 ms (verified via photodiode) displayed on an LCD screen located in front of the monkey and above the haptic device (such that the cursor was approximately at eye level when located at the center of the workspace). A plastic visor was used to mask the monkey’s visual field such that he could see the screen but not his hand or the haptic device handle.

##### 4.1 Trial structure and stimulation conditions

: ↪ c.f. Fig. 2b, Fig. 6c

To facilitate trial-averaging, a discrete set of conditions was used. Non-stimulated trials consisted of delay periods of 0, 300, 400, 600, or 800 ms. Stimulation conditions were structured as follows:

- *pre-trial*: stimulation at least 250 ms before Target Onset, 300 ms delay,
- *delay early*: stimulation at 320 ms after Target Onset, 600 ms delay,
- *peri-go*: stimulation at 20 ms after Go Cue, equivalently 320 ms after Target Onset, 300 ms delay,
- *peri-go no delay*: stimulation at 20 ms after Go Cue, 0 ms delay,
- *peri-move*: stimulation at 50 ms after online movement onset, 300 ms delay.

This condition structure was designed to ensure that (a) each stimulation condition could be paired with an equivalent nonstimulation trial and (b) stimulation would not be predictive of upcoming task events, particularly the Go Cue. The 800 ms delay non-stimulation condition, not included in subsequent analysis, also ensured that monkeys would not make anticipatory movements for the 600 ms delay condition. These conditions were replicated and randomly interleaved for each of the four reach directions, and non-stimulated and stimulated trials were randomly interleaved within each block such that stimulation was present on 40–45% of trials.

### 5 Behavioral analysis

↪ c.f. Extended Data Fig. 1, Fig. 5b,c, Extended Data Fig. 9a

Hand positions were zero-phase low-pass filtered with a 4th order Butterworth filter with 25 Hz corner frequency. Velocities were computed using a smoothing, differentiating Savitzky-Golay filter (2nd order polynomial, 21 ms smoothing). Hand trajectory confidence intervals were calculated using Teetool,^98^ which models the 2d trajectories as a Gaussian process, producing an area that encompasses the 1*Σ* covariance around the mean path. Reach endpoints were labeled where the hand speed fell below 50 mm/sec, and the 95% covariance ellipses were computed.

We computed reach reaction times as the elapsed time from the visual display of the go cue to the time at which the hand speed exceeded a threshold of 5% of the peak reach speed in each trial. We discarded “false-start” trials where the hand speed rose above a higher threshold of 15% of peak, and again fell below 5% before the actual reaching movement. Reaction time effects were computed as the difference in mean RT between stimulated trials and otherwise equivalent non-stimulated trials, i.e. having the same delay period duration. Statistical significance was assessed using the Mann-Whitney *U* test on the sets of non-stimulated and stimulated RTs.

For each stimulation condition (defined by reach direction and stimulation timing), we compared reach velocities at each time between the stimulation condition and otherwise equivalent non-stimulation condition (i.e., with the same delay period, aligned to the same time when stimulation would have occurred). We then computed an unbiased estimate of the norm of the difference in the instantaneous velocity vectors at that time (technically, an unbiased estimate of the squared norm, passed through a square root operation). This technique is from Willett et al. [99] for the unbalanced case (differing trial counts), though we will extend it below in other sections. This approach obtains conservative estimates of distance between trial-averaged quantities, reducing the bias that arises due to estimation noise in the averages by using cross-validation. For each condition pair *c* with *M*_*c*_ non-stimulated and *N*_*c*_ stimulated trials, we form *F*_*c*_ = min(*M*_*c*_, *N*_*c*_) cross-validation folds, and denote the average of the observations (here, 3d instantaneous velocity vectors) in the *i*th large fold as *A*_*i*_, *B*_*i*_ ∈ ℝ^3^ and the average of the small fold *a*_*i*_, *b*_*i*_ ∈ ℝ^3^, for non-stimulated and stimulated trials, respectively. The unbiased estimate for squared norm difference in velocity is given by:

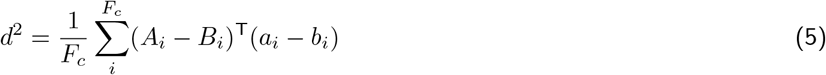

for each conditition pair and at each timepoint.

For each pair of conditions, we compute this distance at each timepoint relative to stimulation onset, and average across conditions. We also summarize stimulation evoked effects on kinematics by averaging the unbiased distance estimate over all Peri-Move conditions and over the time window from 50 ms to 300 ms after stimulation, as in the y-axis of Fig. 5r.

### 6 Task activity space

↪ c.f. Fig. 2f,m-p, Fig. 5f, Extended Data Fig. 9b-d

We define the task activity space as the low dimensional subspace which captures the majority of the variance of trial-averaged neural activity during non-stimulated reaches. We identify the dimensions of this space using principal components analysis (PCA) on non-stimulated activity, and then determine the dimensionality of this subspace by thresholding at 95% of the explainable “signal variance”, defined as in Kobak et al. [100]. As originally noted in Machens [101], the estimated PSTHs 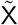 differ from some true underlying firing rates due to noise, and when we estimate the variance explained along each successive PC, some of this variance is due to this residual noise. We proceed by assembling a matrix X_noise_ of scaled single trial residuals with size *RT* × *N* (total tRials, timepoints × neurons). We subtract from each single trial firing rate the corresponding condition mean, and scale each by 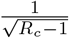 where *R*_*c*_ is the number of trials sampled for condition *c*, set to the minimum trial count recorded for that condition for all neurons in the dataset. We then scale the full matrix by 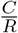, where *C* is the number of conditions and 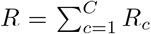 is the total number of sampled trials. This scaling ensures that the total variance of X_noise_ matches the estimated total noise variance of our estimated PSTHs 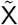. We then compute the singular values *s* and *s*^noise^ of each matrix, and take 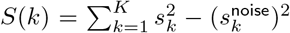 as an estimate of the signal variance captured in the leading *K* principal components. In practice, we find that selecting the dimensionality to capture 95% of the signal variance captures at least 90% of the total variance, a more traditional threshold.

Trial averaged neural trajectories were smoothed with a 20 ms Gaussian kernel for plotting only, and projected into the task activity space.

### 7 Neural effect size analysis

↪ c.f. Fig. 2d,e, Fig. 5e

For each recorded unit, we computed the average change in firing rate (Δ FR) during the stimulation window for the pre-trial condition, relative to equivalent non-stimulated trials. Responses were assessed for significance using a Wilcoxon rank-sum test, with the false discovery rate controlled at 0.05 using the Bejnamini and Hochberg procedure. To compute the stimulation deltas vs. time (Stim Δ), we performed causal 10 ms binning of spike counts aligned with stimulation and averaged across trials within each condition and computing the difference of stimulated and non-stimulated rates. At each of the four stimulation sites, we arranged the Stim Δ as a *T* × *N* matrix (conditions, timepoints × neurons) and performed PCA over neurons. The resulting component scores reveal temporal patterns, which we then averaged across reach directions within each stimulation timing.

To compute the perturbation induced displacement distance, we again used an unbiased estimator of distance between trial-averaged firing rates. We use this estimator to compute the distance between the stimulated neural state and the neural state at the same timepoint on an otherwise equivalent non-stimulation.

#### 7.1 Optogenetic perturbation distance

↪ c.f. Fig. 2m-p, Extended Data Fig. 9c

For the optogenetic stimulation datasets, neurons were recorded sequentially, so the trial averaged firing rates for each neuron were estimated from distinct sets of trials. For each neuron *d* and condition pair *c* (over reach directions and stimulation timings), we have *M*_*c,d*_ non-stimulated and *N*_*c,d*_ stimulated trials, and we form *F*_*c,d*_ = min(*M*_*c,d*_, *N*_*c,d*_) cross-validation folds over trials. Dropping the condition and timepoint subscripts *c* and *t*, we denote the trial-averaged firing rate for neuron *d* within the *i*th large fold as 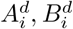 (non-stimulated, stimulated) and the average of the small folds as 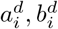. Let the difference between stimulation and non-stimulation for the large and small folds be 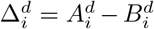 and 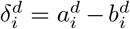. Because the neural recordings are sequential, we sum across the individual unbiased estimates of squared distance along each dimension (neuron) separately, to arrive at an unbiased estimate of squared distance in the high-dimensional neural space.

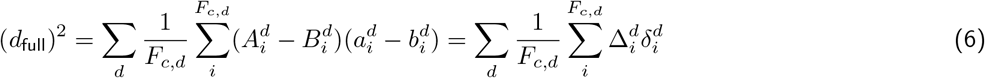

for each condition pair and at each timepoint.

We computed this distance at each timepoint for each pair of stimulation and non-stimulation conditions. We then computed the length of the perturbation vector projected into the task activity space. This estimator cannot sum across squared distance estimates for individual dimensions because of the dimensions are linearly combined through the projection matrix. Instead, to compute the projected distance in a *K* dimensional space defined by the projection matrix W = [*w*_1_ *w*_2_ … *w*_*K*_] ∈ ℝ^*D*×*K*^, we use a strategy of summing over all possible combinations of folds on each dimension. We will implicitly compute the squared distance along projected dimension *k* as (dropping the condition subscript as well):

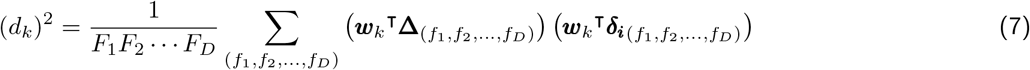

Where each 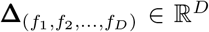 and 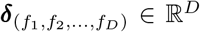 are the assembled vectors of large fold and small fold differences, taking the *f*_1_th fold for the first dimension, the *f*_2_th fold for the second dimension, etc. Fortunately, this expression simplifies so that the full combinatorial sum need not be computed explicitly. Expanding this out, and dropping the *k* superscript for brevity, we have

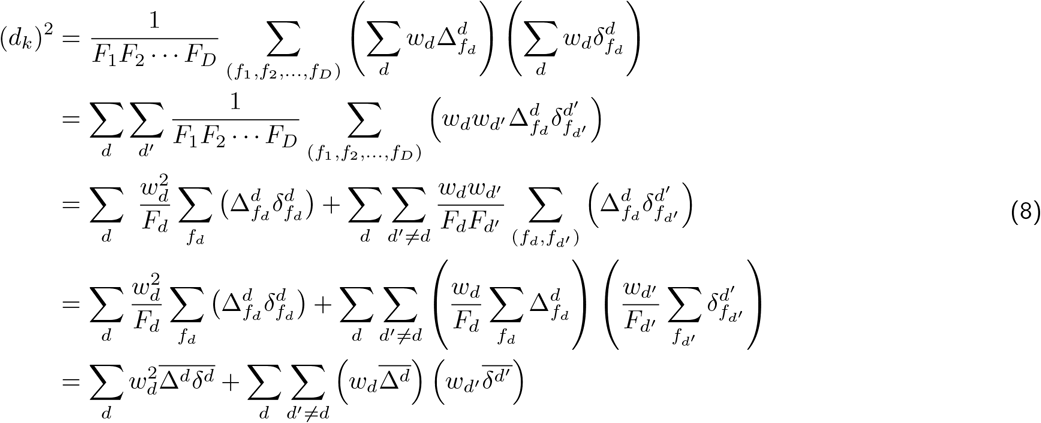

Defining matrix A ∈ ℝ^*D*×*D*^ as:

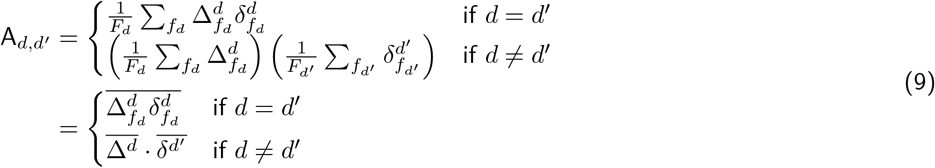

Finally, we have:

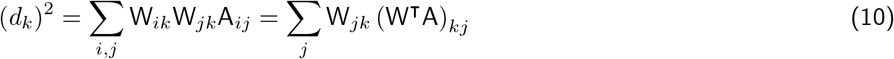

Summing over dimensions *k* in the task activity space, we arrive at an unbiased estimated of the projected (squared) distance:

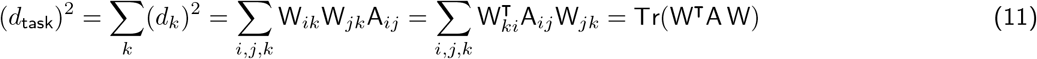

for each condition pair, and at each timepoint.

#### 7.2 ICMS perturbation distance

↪ c.f. Extended Data Fig. 9b-c

For the ICMS datasets, neurons were recorded simultaneously on the Neuropixels probe, so the unbiased distance estimators are simpler. The unbiased estimate of the (squared) perturbation distance in the full neural space is:

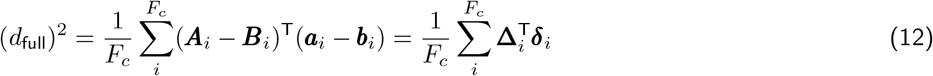

for each condition pair and timepoint. The perturbation distance projected into the task activity space via projection matrix W is estimated as

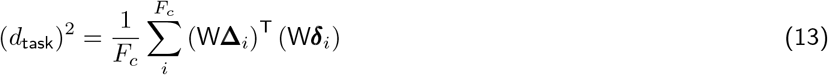

##### Task space diameter normalization

At each stimulation site, we normalize these distances by a measure of the scale of non-stimulated neural activity. We define the task space diameter as the maximum distance between any two points on the non-stimulated neural tractories, for any two conditions and timepoints, as measured using the unbiased estimator of distance projected into the task activity space.

### 8 Latent variable model

In this section, we discuss the design and implementation of the latent variable model used to fit non-stimulated and stimulated neural responses. Our approach here is based off of a simplified version of GPFA,^102^ in that task-related signals in neural firing rates are treated as linear readouts of a low-dimensional collection of latent variables modeled as Gaussian processes. Unlike in GPFA, we fix the common autocorrelation timescale and the linear readout (using standard factor analysis). We also include an low-rank additive term during stimulation, and a rectified Poisson likelihood model, which deals gracefully with neurons whose firing is suppressed as a result of stimulation.

Let 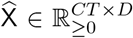 be trial-averaged firing rates for non-stimulated conditions, where *C* is the number of conditions, *T* is the number of time bins, and *D* is the number of recorded neurons. Let N_*c,t,d*_ denote be the number of trials averaged in each corresponding entry. Similarly, let 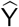 be the trial-averaged rates for the corresponding stimulation conditions, with the same size, where each entry is average over M_*c,t,d*_ trials. In all versions of the model that follow, these spike counts are treated as observations of each neuron’s underlying firing rate, given by X and Y with the same sizes. We take spike counts in non-overlapping 20 ms bins, and assume that all rates are normalized as spikes/bin in the derivations that follow.

#### 8.1 Unconstrained latents version

↪ c.f. Fig. 2g-j, Extended Data Fig. 2a,e,f, Fig. 5g-i,m

First, we fit a version of the model where firing rates during non-stimulated and stimulated conditions are linear readouts of independent latent variables, *z* and 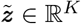, as illustrated in the graphical model show in Extended Data Fig. 2a. This version infers task-related modulation of the non-stimulated and stimulated data independently. For each condition and timepoint, these firing rates are determined by

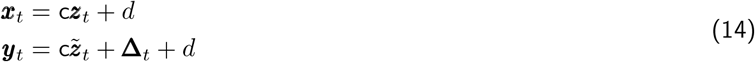

Here, *C* ∈ ℝ^*D*×*K*^ is the fixed readout mapping latents to neurons, initialized using standard factor analysis, and ordered such that the leading latents explain the most variance in neural space. *d* ∈ ℝ^*D*^ is a vector of baseline rates. To ensure temporal smoothness, *z* and 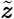 have independent Gaussian process priors with a squared exponential kernel of the form 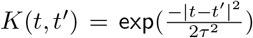 with characteristic timescale *τ* = 100 ms. We also break the autocorrelation before and after the onset and offset of stimulation for 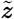 only, to allow for abrupt changes in task related modulation due to stimulation.

Δ ∈ ℝ^*T* ×*D*^ is a matrix of purely additive influences of stimulation on neural firing rates. We constrain Δ to be low rank to avoid overfitting (rank 5, although our results were not sensitive to this choice). This corresponds to the assumption that the additive influences can be captured by a mixture of several timecourses applied to different dimensions of neural activity.

#### 8.2 Contraction-only version

↪ c.f. Extended Data Fig. 9k-m

We also fit a second version of the model in which stimulation latents 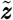 were determined directly by the non-stimulated latents *z* through a linear contraction operator around a fitted centroid (*z*^cent^ ∈ ℝ^*K*^).

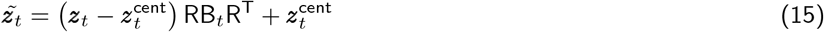

Here 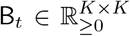 is fitted diagonal matrix whose entries contract (or expand, if greater than unity) individual latent axes, *R* ∈ ℝ^*K*×*K*^ is a fitted rotation matrix that orients the modes of the contraction as needed with the latent dimensions. The individual entries of B_*t*_ have a Gaussian process prior as well, along with the fitted centroid *z*^cent^.

#### 8.3 Shuffled and Poisson Null versions

↪ c.f. Fig. 2k, Fig. 5k, Extended Data Fig. 9e

We also fit two versions of the model to provide distributions of latents under the null hypothesis that stimulated and non-stimulated latents were identical (i.e.,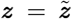). In the first version, *Shuffle*, we assembled single trial spike counts from paired non-stimulated and stimulated conditions and shuffled the labels. This yields trial-averaged firing rates under the null hypothesis, such that differences between non-stimulated and stimulated firing rates result from differing amounts the additive Δ (which is now mixed into both trial averages) and noise. We tracked the fraction of true-stimulation trials contributing to the shuffled condition averages for non-stim and stim (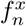 and 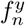 for neuron *n*) and fit the model where both *x* and *y* incorporated a scaled contribution from the additive Δ term to match:

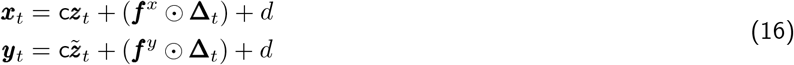

In the second version, *Poisson Null*, we took the fitted non-stimulated latents *z* and the additive Δ from the unconstrained model fit, and regenerated synthetic spike count data under the null hypothesis. Specifically, we set 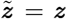, computed the predicted firing rates, and then generated spike counts for each trial as Poisson-distributed samples from these rates. We then fit the model to these synthetic data as before, fitting *z* and 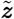 (and Δ) independently.

Both versions were used to compute a distortion metric (see §8.6) of distortion under the null hypothesis that stimulation does not distort the underlying latent task variables.

#### 8.4 Rectified Poisson likelihood model

↪ c.f. Extended Data Fig. 2b-d

We treat the observed single trial spike counts as individual observations of inhomogenous Poisson process with rates given by matrices *X* and *Y*, with the same size as 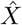, *Ŷ*. For an individual entry 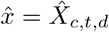 is computed by averaging *N*_*c,t,d*_ trials, where the *i*th trial, the bin contained 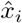 observed spikes. The modeled rate is *x* = *X*_*c,t,d*_, which is normalized to spikes per time bin. Then the total log-likelihood is:

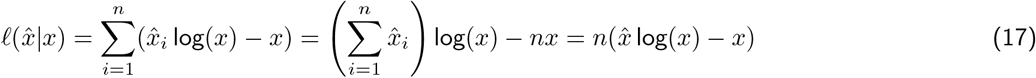

Therefore, due to the additivity of the Poisson distribution, we can work only with the trial-averaged rates and trial counts.

Firing rates in X and Y can be negative, particularly if the additive term leads to suppression of individual neurons. We account for this in the likelihood model by passing the rates through a soft-plus rectifiying nonlinearity with fixed hyperparameters *κ* and *E*.

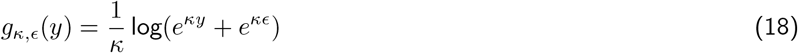

Here, *κ* controls the sharpness, with *κ* ≫ 1 approaching ReLU. *E* determines the minimum value as *y* → − ∞. We used *κ* = 10 and *ϵ* = 0.5 spikes/sec, for which *g*(·) is plotted in Extended Data Fig. 2c. Note that we will use *y* and *ŷ* in the description that follows, but the same likelihood is used for the non-stimulated fitted rates *x* and data 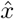 as well.

Unfortunately, we run into a few issues if we implement the likelihood simply as (ignoring a scalar trial count factor *m*):

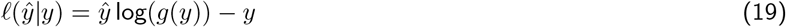

First, since *g*(*y*) *> y* everywhere, there is always pressure for *y* to be slightly larger than *ŷ* to compensate, especially when *ŷ* is closer to *ϵ*. This we address by wrapping *ŷ* in *g*(·) as well:

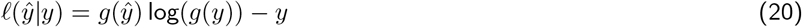

Second, the presence of the softplus rectification distorts the likelihood landscape when *y* or *ŷ* are close to *ϵ* or when *y* is negative. Taking the derivative with respect to *y*, we have:

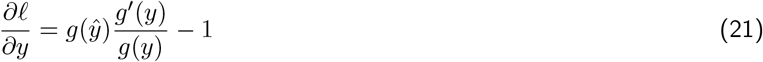

which will be zero when:

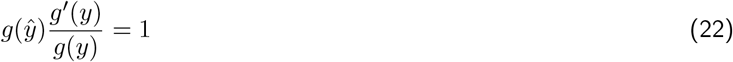

For *ŷ, y* ≫ *ϵ*, we have *g*^′^(*y*) ≈ 1 and *g*(*y*) ≈ *y*, so the maximum likelihood is correctly located at *y* = *ŷ*. However, as *ŷ* approaches *ϵ* from above, this maximum disappears. To address this, we introduced a transformation function *h*_*ŷ*_ (*y*), to be specified below, on the second log likelihood term as well, which acts to transform *y* but whose form depends on *ŷ*.

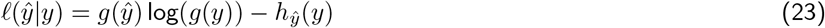

Our goal now is to design *h* so as to shape the log likelihood surface in *y, ŷ* so as to address these distortions caused by the rectification function when *y* or *ŷ* are close to ϵ or when *y* is negative. When *y* and *ŷ* are greater than *E*, we want the log likelihood to match a standard Poisson observation model. Also, high spike counts (*ŷ* ≫ 0) should be discouraged when the rate is zero or negative (*y* ≤ 0), but negative firing rates are allowed when near-zero spikes are observed, essentially treating the spike counts as a censored observations of the rates. To accomplish this, we design the log likelihood surface piecewise in regions of *y, ŷ*. First, we let *f*_0_ be the gradient with respect to *y* of the original Poisson likelihood with *g*(·) applied to the inputs:

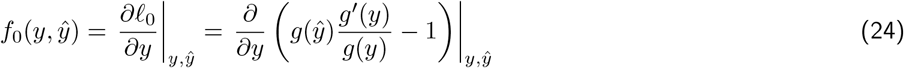

We also define *U* (*ŷ*) = argmax_*y*_ *f*_0_(*y, ŷ*) for *ŷ > ϵ*. We then design a revised log likelihood surface, such that its gradient *f* has the following region-specific properties:

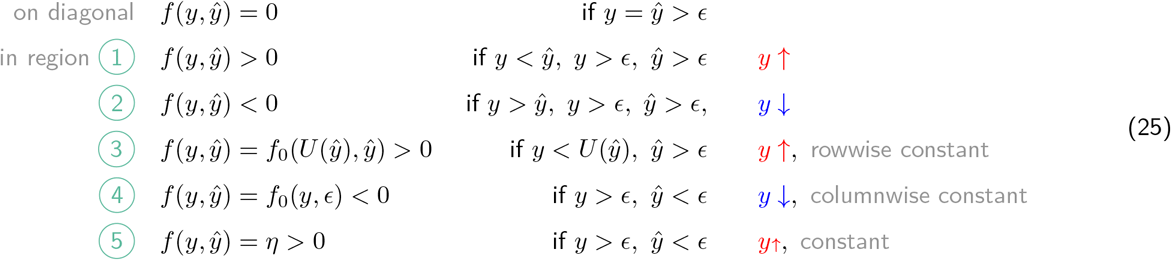

where the circled region numbers refer to the schematic in Extended Data Fig. 2b. We also introduce a third likelihood hyperparameter *η* = 0.01 (in addition to *κ* and *ϵ* that parameterize *g*(·)), which acts to gently penalize unnecessarily large, negative rates even when the observed spike count is very low (region ➄).

To satisfy these requirements for *f*, we define *h* implicitly by its derivative with respect to *y*:

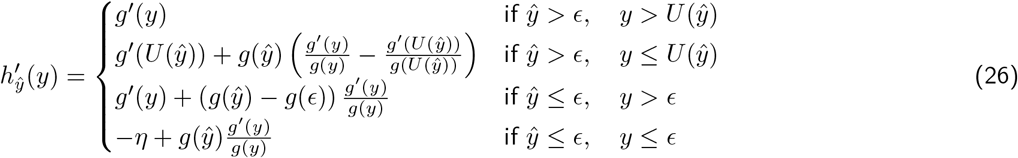

Although this formulation is complex, the resulting log likelihood surface is straightforward (plotted in Extended Data Fig. 2d). Overall, this approach is an intuitive extension of the Poisson observation model allowing for negative firing rates (i.e., suppression via inhibition), while remaining diffentiable and straighforwardly optimized.

#### 8.5 Model fitting and effective contraction factor

↪ c.f. Fig. 5o-r, Extended Data Fig. 9m-p

All versions of the model were fit using Manopt, a toolbox for optimization on manifolds^103^ using the Riemannian trust-regions solver. This allowed up to optimize the log-likelihood over the fitted parameters while incorporating non-negative and low-rank constraints on those parameters. We computed approximate confidence intervals for fitted parameters using the inverse of the observed Fisher information matrix (the Hessian of the negative log-likelihood function).

From the fitted parameters, we could compute an effective contraction factor *β*_*t*_. In all versions of the model, this was defined in terms of the sum of cross-condition variance over neurons for the stimulated firing rates relative to the non-stimulated firing rates. We define 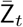 and 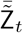 as the *K* × *C* (latents × conditions) matrices of centered latents at time *t*, having subtracted the centroid across conditions, and Var_*c*_ as the variance across conditions, such that the squared contraction factor equals:

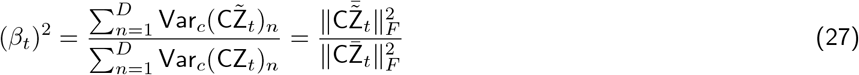

where 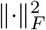 denotes the squared Frobenius norm of the matrix.

For the unconstrained model, the contraction factor was computed directly from the fitted latents, and confidence intervals were estimated by resampling Z and 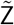 under the posterior to sample a distribution for the contraction factor.

For the contraction-only model, this contraction factor can be written directly in terms of the other fitted parameters. Here, we define 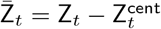 as the *K* × *C* (latents × conditions) matrices of contraction centroid-centered non-stimulated latents. Noting that B is diagonal, we have:

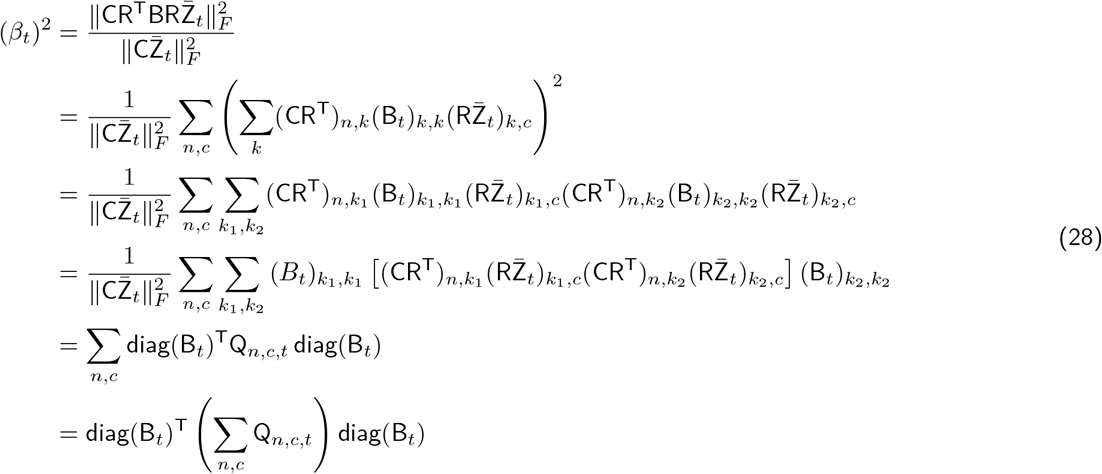

where we define the inner matrices Q_*n,c,t*_ ∈ ℝ^*K*×*K*^ as:

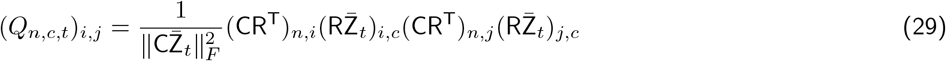

Consequently (*β*_*t*_)^2^ is a quadratic form of normally distributed variables (diag(B_*t*_) under the posterior, computed from the Fisher information matrix). Therefore, (*β*_*t*_)^2^ follows a generalized χ^2^ distribution.^104^ From this distribution we compute confidence intervals for *β*_*t*_.

We also summarize the contraction factor in Fig. 5o,r and Extended Data Fig. 9n-p as the average *β*_*t*_ over the stimulation interval (truncated at 200 ms if longer).

Finally, we verified that the contraction factor could be estimated accurately by using simulated transformations of real data. We began with the fitted non-stimulated latents for each dataset and applied a known contraction factor uniformly across all latent dimensions. This provided the simulated latent states for a given contraction factor. We computed neural firing rates in each condition by projecting out to neural space, sampled spike counts from the corresponding Poisson distribution, and then used these simulated datasets to refit the latent variable model. For all datasets and across the full range of contraction factors from zero to one, the estimated contraction factor tracked the true contraction factor well (Extended Data Fig. 9m).

#### 8.6 Stimulation distortion metric

↪ c.f. Fig. 2k, Fig. 5k, Extended Data Fig. 9b

We defined a metric to quantify the amount nonlinear distortion in the underlying latent variables induced by stimulation. This metric is based on the linear, Euclidean shape similarity metric defined in Williams et al. [48]. For each comparison, we first performed a Procrustes superposition allowing translation, reflection, and rotation but *not* scaling between the full non-stimulated and stimulated latent trajectories (Z and 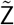). We define 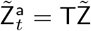 be the *K* x *CT* (latent dimensions x conditions, timepoints) matrix of stimulated latents, aligned to the non-stimulation latents via the Procrustes transformation T ∈ ℝ^*K*×*K*^ with T^⊤^T = I. Given these aligned latents, we project out to neural firing rate space and compute the Euclidean distance, summed over conditions and timepoints within the peri-stimulation period.

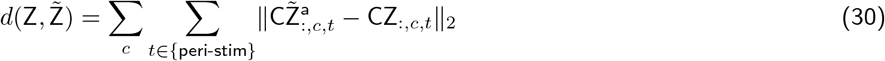

We computed this distortion metric for the fitted latents in the uncontrained model, and we computed confidence intervals by resampling from the posterior of the latents and recomputing the distortion. We also computed the distortion metric for the Shuffled and Poisson-null latents and took the 95th percentile of the distortion metric under these null hypotheses as the significance threshold.

#### 8.7 Goodness of fit

↪ c.f. Fig. 2l, Fig. 5l,n

We computed the final goodness of fit of each model using the Poisson deviance. With observed spike counts 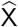, fitted firing rates (in spikes per bin) X, the total deviance, summed over conditions, timepoints, and neurons is given by:

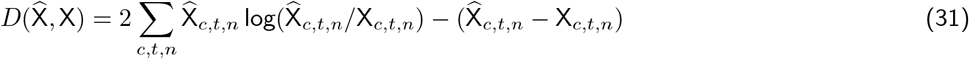

We normalize the deviance by number of observations (conditions × timepoints × neurons), and compute the jackknifed standard error over neurons.

#### 8.8 Single trial analysis

↪ c.f. Extended Data Fig. 9f-j

Lastly, we fit a single trial version of the model, where an independent latent variable trajectory was fit for each non-stimulated and stimulated trial. This version is equivalent to the unconstrained latents model in §8.1 if each trial is considered as an individual condition.

Given the fitted single trial latent states, we focused on the states at the final timepoint of stimulation (or the equivalent time points on otherwise equivalent non-stimulation trials). We centered the single trial states around their respective within-condition means and computed the covariance of the single trial residuals of non-stim and all stim trials separately as Σ_*n*_ and Σ_*s*_. Separately, we computed the covariance of the condition means around the grand mean as Σ_*N*_ and Σ_*S*_. Each of these covariances are *K* × *K* where *K* is the dimensionality of the latent space.

We first compared the covariance of the residuals Σ_*n*_ and Σ_*s*_ directly using the correlation matrix distance (CMD),^105^ defined as:

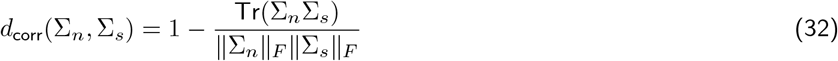

This quantity is zero when the correlation matrices are equal up to a scaling factor and its maximum is one. We evaluated the CMD statistic against the null hypothesis that the residual covariances are identical, using a permutation test by shuffling the non-stim and stim residuals before computing the covariances.

We also considered whether the change in covariance structure from non-stimulation to stimulation that acted on the condition means was similar to the change in covariance in the single trial residuals. To measure this change in covariance, we computed non-stim whitened versions of the stimulation covariances for both the condition averages and the single trial residuals as:

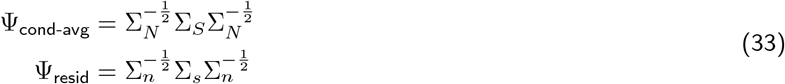

Ψ_cond-avg_ and Ψ_resid_ ∈ ℝ^*K*×*K*^ describe the change in the covariance of latent states from non-stimulation to stimulation for the condition averages and residuals, respectively.

We then the correlation matrix distance (CMD) between these matrices as:

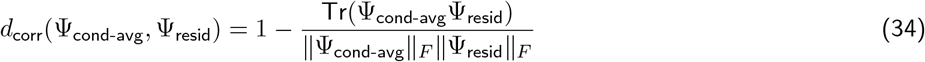

If this distance is small, it implies that the transformation that applies to the covariance structure of the condition averages is similarly to the transformation that reshapes the single trial variability around the condition averages.

To establish statistical significance, we consider the null hypothesis in which the shape of Ψ_resid_ is uncorrelated with Ψ_cond-avg_. To assess this, we compute the *d*_corr_(Ψ_cond-avg_, Ψ_resid_) statistic under random orientations of the stimulation residuals. Multiplying the stimulation residuals by a random rotation matrix Q has the effect of transforming the covariance matrix to 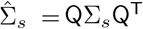.

### 9 Latent linear dynamical system model

↪ c.f. Extended Data Fig. 4a,b

We fit latent linear dynamical system (LDS) models as a standard first step in describing the observed neural population dynamics. These models describe the evolution of a low dimensional latent variable in terms of linear dynamics and piecewise-constant inputs as:

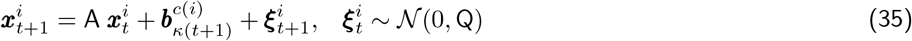

Here, *i* = 1, …, *M* indexes the non-stimulated conditions (reach directions, delay periods), *c*(*i*) returns the reach direction index corresponding to that condition, and *κ*(*t*) indicates the time epoch, establishing the time where the piecewise constant inputs transition from ‘hold’ to ‘go’.

Observed PSTHs are modelled as a linear combination of latents with a constant offset.

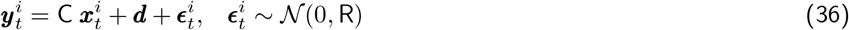

The model parameters 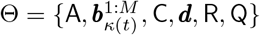 are fit using Expectation Maximisation. Trial averaged firing rates were binned at 10 ms in causal, non-overlapping bins. The dimensionality K and time points at which inputs transition from ‘hold’ to ‘go’ are determined via a gridsearch, selecting the values that achieve lowest mean-squared prediction error on held out validation data. For this purpose, we evaluate on data prior to stimulation onset on stimulation trials is used as the validation set. This ensures that the test data used in Fig. 3m-p is truly unseen across all model classes. These optimal hyperparameter values are fixed across all versions of the LDS model and also used in specifying the dimensionality and transition from ‘hold’ to ‘go’ in the high-dimensional model described below in section 10.

We generated 1000 resampled model fits by resampling non-stimulation trials with replacement from each condition, computing trial averages and fitting the LDS model. From these resampled fits, we computed a density estimate of the eigenvalues of the dynamics matrix A.

#### 9.1 Model predictions following stimulation

↪ c.f. Extended Data Fig. 4b-e,g, Fig. 2p

Having fit the model using only non-stimulated data, we then predict the LDS model’s responses following optogenetic perturbation by using the firing rates at the last time bin during stimulation as the initial condition, 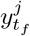 for stimulation condition *j*. We estimate the initial condition for the latents from the empirical firing rates at stimulation end:

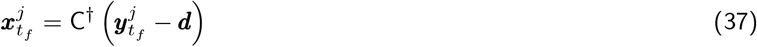

We then run the latent dynamics forward from this initial condition according to Equation (35) (but without injecting innovations noise). We compute the Euclidean distance in neural space between the stimulated and non-stimulated model predictions as a function of time. We computed confidence intervals for these decay curves by resampling stimulation trials to re-estimate the initial condition for the latents, and then evaluating model predictions forward using the resampled model fits. We computed the evoked energy as a summary statistic of these decay curves, defined as the integral under this perturbation distance over time, again normalized by the task space diameter.

We also fit a second version of the LDS model using both non-stimulation and all stimulation conditions for model fitting, and evaluted the predictions of this model to stimulation in the same manner. To simulate stochasticity (e.g., of opsin expression) in the stimulation initial condition, we computed the difference in firing rate (stim minus non-stim) at the final time bin of stimulation, randomly permuted these offsets, and then added the shuffled offsets onto the non-stimulated firing rates. We then evaluated the model’s predictions from this shuffled stimulation initial condition.

#### 9.2 LDS model analysis, Schur decomposition

↪ c.f. Extended Data Fig. 4f,h

To better understand the differences between the LDS model fit only to non-stim and the model fit to both non-stim and stim, we computed the Schur decomposition of the dynamics matrix.^106,107^ The real Schur decomposition expresses a square matrix as

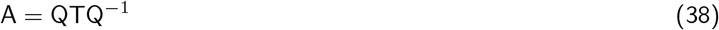

where Q is an orthogonal matrix and T is lower quasi-triangular (possibly with 2 × 2 blocks along the diagonal). ^a^ Here, Q defines an orthonormal basis in *K*-dimensional latent space. T, called the real Schur form of A comprises the eigenvalues of A along the diagonal (with 2 × 2 blocks for complex eigenvalue pairs), as well as possibly nonzero entries below the diagonal. If A were a normal matrix, such that A^⊤^A = AA^⊤^, T is be strictly quasi-diagonal, and the Schur decomposition reduces to a diagonalization by eigenvalues. If A is non-normal, then a lower triangular entries at T_*i,j*_ indicates a functional feedforward interactions from dimension *i* onto *j* in the latent basis of U.

The Schur decomposition is not unique; specifically, any ordering of the eigenvalues along the diagonal of T is valid. We sorted the eigenvalues (or complex eigenvalue pairs) in increasing order of how strongly the stimulation vector projected along the corresponding eigenvector (or plane spanned by the corresponding eigenvector pair). This allowed us to reveal functionally feedforward interactions from latent modes largely unaffected by stimulation onto modes strongly affected by stimulation. In visualizing the real Schur form, we also computed the total feedforward input by adding in quadrature the entries in the corresponding row of T.

Lastly, we quantified the non-normality of the dynamics matrix A across the different model variants using the normalized Henrici’s departure from normality measure,^108^ defined as:

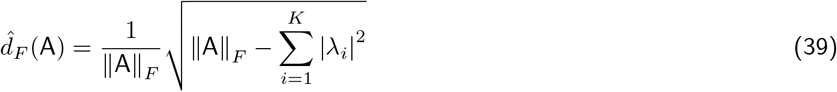

This measure ranges from 0 (normal) to 1 (maximally non-normal).

### 10 An E/I network model with low-dimensional subspace-structure

#### 10.1 Model description

↪ c.f. Fig. 2a, Extended Data Fig. 5

We develop a dynamical model of a putative subnetwork of motor cortex able to produce the slowly-varying, low-dimensional activity patterns observed in motor cortical data, while having a direct interpretation in terms of circuit-level properties. To do this, we describe the temporal evolution of a set of network variables 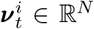 on experimental condition *i* = 1, …, *M* via

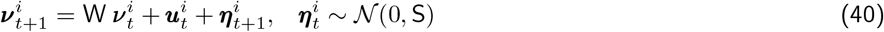

Subsets of ***ν*** reflect difference-from-baseline firing rates (and are hence allowed to go negative) of a pool of excitatory and inhibitory neurons. The connection matrix W is constrained by Dale’s Law and by a requirement that E and I input strength to each unit be balanced. This results in constraints on the dynamics of the form

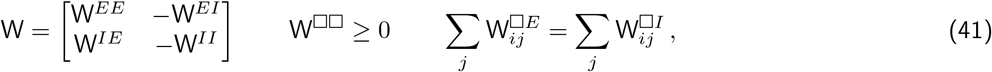

where W^□□^ are matrices with non-negative entries that specify the connection strengths within (*EE, II*) or across (*EI, IE*) the pools of excitatory and inhibitory cells. The reach target and the time of the “go” signal are supplied by additive external inputs (***u***_*t*_), which were constant during each of the hold and movement periods, such that the evolution of ***ν***_*t*_ during each phase was determined by the autonomous dynamics of the network. Recorded neurons are modelled as driven by combinations of network units, so that deviations in PSTHs about their respective background levels (***d***) are given by linear combinations of elements of ***ν***_*t*_ according to a loading matrix (C). The network variables are hence related to recorded firing rates 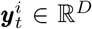 via

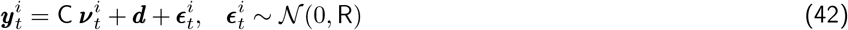

Under this specification, the model network contains more units than recorded neurons in the dataset. Therefore, many different patterns of dynamics in the high-dimensional system could reproduce the recorded PSTHs. However, these solutions may generally be more complex dynamically than the recorded data, with the agreement in PSTH reconstruction depending on precise cancellation of high-variance signals by the specific linear weights in C. In these cases, small changes in C would generate apparent dynamics that differed appreciably from the measured patterns. This contrasts with the widely observed finding that motor cortical activity associated with controlled reaching movements exhibits systematic low-dimensional dynamics that can be consistently reconstructed from many different samples, or mixtures, of recorded neurons.^3,23,56,57,109^

Specifically, we would like the high-dimensional system to match the complexity of the recorded neural activity in two ways: First, the observed shared structure of population activity during simple center-out reaching movements is generic, in the sense that it varies little across recording sessions, sites, animals, or from study to study.^23,56,57^ This was the case here, with principal component projections displaying similar trajectories at all recording sites in two monkeys (Extended Data Fig. 5a). If the underlying patterns of activity in the full cortical circuit were of significantly greater complexity, then different recorded populations, reflecting different linear projections, would instead exhibit different projected structure (Extended Data Fig. 5b). Second, for activity during phases of the trial that are internally paced – including during execution of the movement – a low-dimensional projection of the recorded activity has been reported to be sufficient to accurately predict its future evolution.^3,28,29,110,111^ This property was also reflected in our recordings by the ability of a low-dimensional linear dynamical system (LDS) model to accurately recapitulate population activity (Fig. 3e).

To ensure that these characteristics were also typical of the model dynamics, we regularise the high-dimensional system as follows. We randomly selected a *K* × *N* rectangular projection matrix J, where *N* is the number of network units and *K* is a dimension that suffices to capture the majority of variance in the recordings. The row space of J defined a *K*-dimensional subspace of network activity, which will be referred to as the *task-dynamics space*. We constrained the evolution of the network state projected into this subspace to be unaffected by the orthogonal component of the state. That is, we required

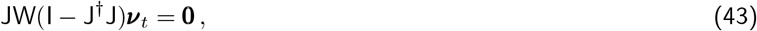

where J^†^ = J^⊤^(JJ^⊤^)^−1^ denotes the pseudo-inverse. Under this constraint, the evolution of the network state projected into the subspace defined via J, can be described as

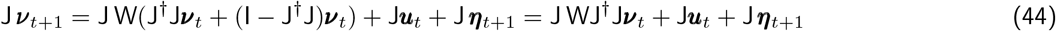

which shows that the state J ***ν***_*t*+1_ only depends on activity within J at the previous time point. Letting ***x***_*t*_ = J***ν***_*t*_, the dynamical evolution within the this low-dimensional space can be described via low-dimensional, linear dynamics A = J WJ^†^. Thus, under the constraint in Equation (43), dynamics in the J-defined subspace are *self-contained*, with properties similar to those observed in motor cortical data.

To ensure that the model dynamics matched those observed in the recordings, we also required that the reconstruction of the measured PSTHs be based only on the projection of the network state into this subspace. That is, we set

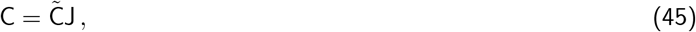

where 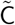 is a *D* × *K* matrix that can be learnt from data. Furthermore, we assume that the inputs are low-dimensional and arrive within the J-defined subspace. We thus assume the additional input structure

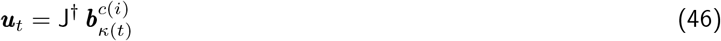

*κ*(*t*) ∈ {1, 2} indicates the indices for the constant inputs supplied during the hold, or movement period and *c*(*i*) indicates the target direction of the *i*th PSTH. In addition to the constraints in Equations (43) and (41), we also impose sparsity in the connectivity matrix W by constraining 75% of its entries to be equal to zero.

#### 10.2 Model fitting

We fit this model by maximum likelihood learning using the Expectation Maximization algorithm (EM).^112^ Since the data reconstruction only depends on ***x***_*t*_ = J***ν***_*t*_ when the dynamical evolution is self-contained, we can write the objective function in terms of ***x***_*t*_, instead of representing the full state according to ***ν***_*t*_.

The optimisation objective of the algorithm is a lower-bound to the marginal log-likelihood (with hidden-states integrated out). For the model we consider here, it is given by

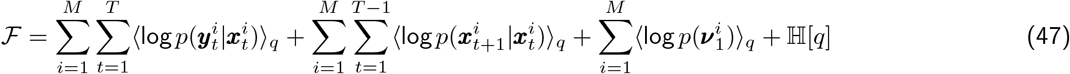

Angled brackets ⟨ · ⟩_*q*_ denote expectations with respect to a distribution 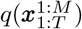 over hidden network states within the task-dynamics space defined by J and ℍ[*q*] is the entropy of 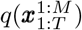. The constraint in Equation (43) is enforced via the addition of a penalty term to the EM objective function. The penalty term takes the form

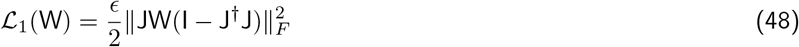

Similarly, we encourage the E/I balance constraint in Equation (41) via a quadratic penalty of the form

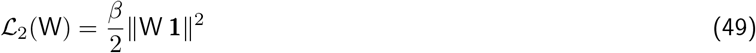

The parameters *ϵ* and *β* are determining the trade-off between fitting the data and satisfying the constraints and are increased gradually throughout the optimisation. While it is possible to use constrained optimization approaches to implement the linear equality constraints in Equation (43) and (41) (for example interior-point methods), these quickly become too computationally costly for larger networks. The quadratic penalty approach used here is more computationally efficient for large network sizes, since gradient descent with upper- and lower-bound constraints can be used to learn W.

Our final optimization objective can be written as

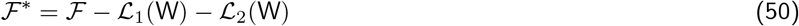

We maximise this objective with respect to the distribution over hidden states 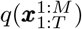 and the model parameters 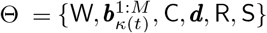. The task-dynamics space mapping J is sampled at random with 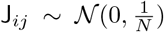 and is held fixed throughout model fitting. In order to fit the model parameters Θ via EM, we exclusively use recorded PSTHs from experimental conditions without optogenetic stimulation. At no time throughout the model fitting procedure is the model instructed to reproduce PSTHs recorded under optogenetic stimulation.

To fit the model, the network connections in W are initialised randomly and independently according to the weight distribution used in Hennequin, Vogels, and Gerstner [106] using a spectral radius of 0.85 to ensure stability. In a second initialisation step we take the random weight matrix and minimize the norm between the dynamics of an analogous fitted LDS model and JWJ^†^, while penalizing violations of the constrains in Equations (43) and (41). The resulting W matrix is then further adjusted according to the EM algorithm (along with the model parameters Θ) to maximize the probability of the measured responses, while maintaining Dale’s Law, E/I balance, sparsity, and the subspace constraints. The latent dimensionality *K* and change time-point for the piece-wise constant inputs were determined via cross-validation in an LDS model with no connection to a high-dimensional E/I network, but otherwise analogous model structure.

#### 10.3 Hidden-state inference

The optimal distribution 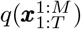 is equal to the posterior distribution of hidden-states 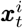 given the observed PSTHs. Computing this posterior distribution over 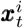 is analogous to Kalman smoothing with low-dimensional linear dynamics JWJ^†^ in the presence of inputs.^113,114^ The output of the Kalman smoother will be the set of posterior means 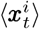, posterior second moments 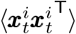 and time-shifted posterior second moments 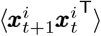. The angled brackets indicate expectations with respect to 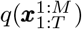.

#### 10.4 Parameter learning

Most of the required parameter updates are available in closed-form, and only involve slight modifications from the solution for the classic linear dynamical system^115,116^ due to the introduction of the task-space mapping J. The update equations are as follows:

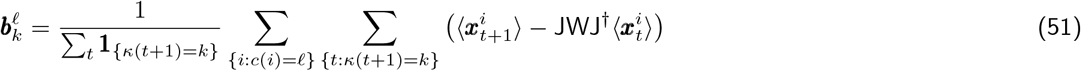

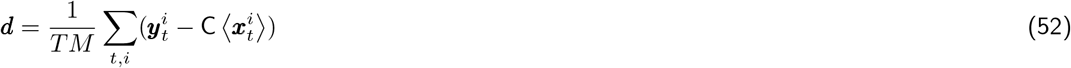

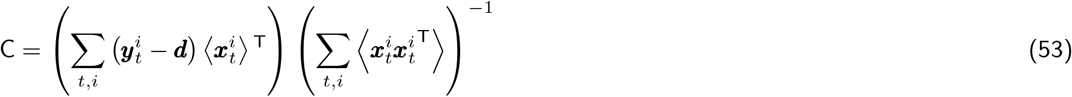

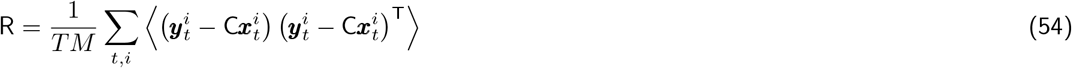

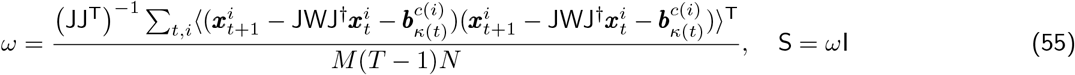

Learning W is done by gradient descent on ℱ*. The E/I sign constraints are included as upper and lower bounds on each entry W_*ij*_ during the optimisation. The relevant gradient for W is given by

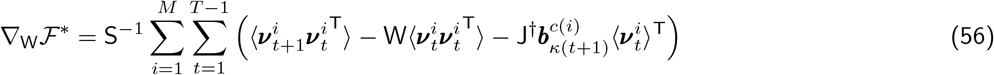

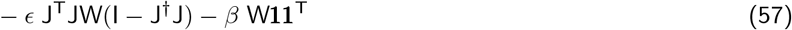

A given level of sparsity is enforced by pre-selecting the number of active, non-zero connections in W at random (but always including the elements on the diagonal), and only including these as parameters in the optimisation.

#### 10.5 Initial state distribution

Initial state distributions are again chosen to maximize ℱ*. We assume that the initial hidden state is drawn from a distribution with 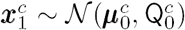. The optimal update can be shown to take the form

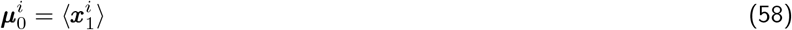

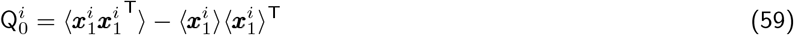

### 11 Evaluating model responses to optogenetic perturbations

↪ c.f. Fig. 2b-d, g–i

We seek to test the model’s responses to inputs that resemble the perturbation patterns of the experiment. To do this, we consider a general class of input patterns that target the excitatory sub-population. We consider stimulation patterns that target different fractions of excitatory cells and with added noise in the stimulation-vector value for each affected cell, representing noise in opsin expression levels. Stimulation vectors used to perturb the network model are generated as

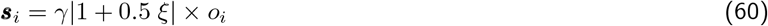

where *ξ* ∼ 𝒩(0, 1), *o*_*i*_ = 1 with probability *p*^stim^ = 0.75, and *o*_*i*_ = 0 otherwise, and *γ* is the stimulation amplitude. It was selected based on cross-validation (Figure Fig. 3c,e) or such that the relative size of the stimulation distance vs. task-space diameter of the network responses approximately match the ratio that was observed empirically (Figure Fig. 3f,g).

We test the model by letting it evolve forward in time without noise, starting from the learned initial condition 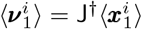 on the matched non-stimulated trial. During the appropriate time window, we introduce the stimulation input ***s*** to the dynamical system according to

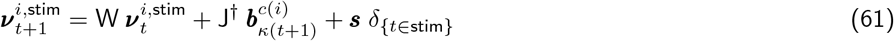

where ***s*** is constant and applied at each time-step of the 200ms stimulation window (during which *δ*_{*t*∈stim}_ = 1 and outside of which *δ*_{*t*∈stim}_ = 0).

In order to relate the stimulation responses in network space to the recorded data, we pick a random subsample of 5 · *K* units from 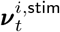 and perform ridge regression to estimate a set of linear weights and offset to predict the recorded data on non-stimulated trials and on stimulated trials from all but one of the conditions. Predictions of neural stimulation responses were made based on network responses to stimulation inputs on the held-out stimulation condition not used for training the regression weights. This step is necessary for establishing a link between network responses and measured neural responses, since there is no a priori relationship between the stimulation-induced translation in neural space and in network space in the model.

### 12 Evaluating training and test performance of models

↪ c.f. Fig. 2e-f

We report the performance in terms of the fraction of signal variance explained on training data and held-out test data. The test dataset contained neural PSTHs that were recorded for reach conditions with a 400ms delay period that were not included in subsequent analyses since they have no analogous stimulation condition. To evaluate the model performance on this dataset, we held all model parameters fixed and iterated posterior inference and updating of the initial conditions of each model until convergence. The model performance was then evaluated by predicting forward from the learned initial condition, holding all other model parameters fixed.

### 13 Comparison with SOC architecture

↪ c.f. Extended Data Fig. 3

We follow the procedure from Hennequin, Vogels, and Gerstner [14] to obtain stability optimized circuits (SOCs). Analogously to our E/I network model, we choose an equal ratio between E and I cells and set the probability of a non-zero synaptic weight in the initial weight matrix to *p* = 0.25. Following [14], the initial spectral radius is chosen to be *R* = 10. We optimize the network until a spectral abscissa of *α*_*abs*_ = 0.5 is reached and maintain E/I balance throughout this optimization. After the training procedure, the network activity is evaluated by solving

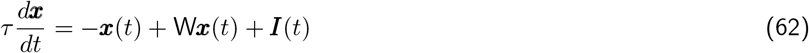

Following [14], we set *τ* = 200 to match the time-scales observed in motor cortical activity patterns.

To obtain a qualitative agreement between the SOC and the recorded PSTHs, we follow the approach in [14] and drive the system to a steady state solution corresponding to a random linear combination of the leading two eigenvectors (which we will denote as ***g***_1_, ***g***_2_) of the Gramian matrix ***G*** satisfying the Lyapunov equation:

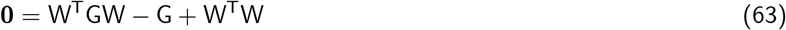

The vectors ***g***_1_ and ***g***_2_ reflect the activity patterns resulting in the largest amount of evoked energy in the system.

At the end of the preparatory period, we would like the network state to settle to the state

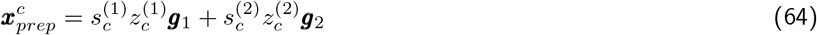

where 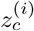∼ Uniform[0.5, 1] and 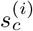 is a random sign (±1) for each target direction *c* = 1, …, 4 and *i* = 1, 2. To reach this steady state, we supply the network with an vector-valued input whose direction is constant and whose magnitude linearly increases to 1 at the time of the go cue:

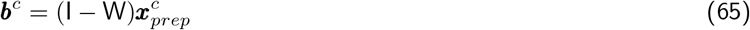

At the time of the go cue, we turn this input off which causes the network to autonomously evolve forward in time starting at the initial state 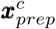. The leading directions of variance of the resulting activity patterns across each of the four target directions define the task-activity space of the SOC network. To evaluate the SOC against input perturbations to excitatory units in the network, we follow the same approach as we did for the self-contained E/I network model. The task-activity space of the SOC is determined by evaluating the task-responses and computing the subspace capturing 95% of variance using principal components analysis. To compute the impulse response of the SOC to a stimulation input ***s*** at *t* = 0, we can solve the differential equation in (62) to obtain

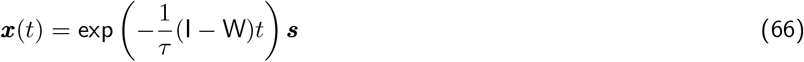

### 14 Theoretical analysis of network responses

↪ c.f. Fig. 4

We can understand the robustness in the network in terms of the random stimulation pattern only having a 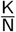 by-chance projection into the K-dimensional subspace J, where the slow, task-relevant dynamics are sensitive to inputs. In the large N limit, any random perturbation will only interact with the unstructured dynamics outside of J, since the self-containment constraint in Equation (43) prevents coupling of responses from outside J into any of the slow modes of the system within J over successive time-steps. Similar to the responses observed under random unstructured E/I connectivity, random inputs decay as a sum of exponentials in the learned E/I network dynamics. Overall, for a low-dimensional, self-contained task-dynamics space embedded in a large network, the probability of a random input perturbing the slow and structured eigenmodes of the network dynamics will vanish.

The other components of the response to stimulation-related input can be understood in terms of the properties of a balanced E/I network. Balanced E/I networks are non-normal dynamical systems in which perturbations along a “differential” mode of contrasting deviations in E and I activity are transiently amplified into a “common” mode of E/I co-activation, which then decays rapidly.^14,50,51^ We found that this pattern of dynamics necessarily dominated the amplification of random perturbations along the differential mode (i.e. targeting only E cells in the population), even for networks optimised to implement systematic long-time-scale dynamics, such as those describing the slow evolution of activity patterns in motor cortex. We can see that this structure is necessitated based on the sign pattern in the dynamics and the stimulation input:

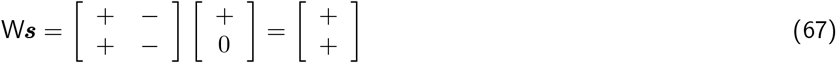

In addition to amplification along a co-activation pattern dominating the stimulation response, we also found that the network produced response variance along this direction during normal task-related activity in the absence of stimulation related inputs. This behavior explained why the network showed a large projection of the stimulation response into the task-activity space, while at the same time being robust to the stimulation. To better understand why this structure arises in the network, we consider the singular value decomposition of a general dynamics matrix W satisfying Dale’s law sign constraints. Using the singular value decomposition, the dynamics matrix can be written as

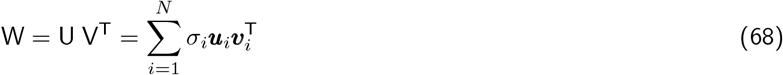

where = diag(*Σ*_1_, …, *Σ*_*N*_) contains the singular values of W along its diagonal, and U = [***u***_1_ … ***u***_*N*_ ] and V = [***v***_1_ … ***v***_*N*_ ] are matrices containing the orthonormal set of left and right singular vectors, respectively. The singular vectors in U and V are the eigenvectors of the matrices WW^⊤^ and W^⊤^W, respectively. The sign-constraints in W imposed by Dale’s law, will also lead particular sign-structure in these matrices. We have

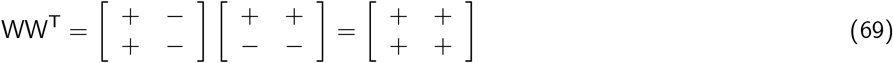

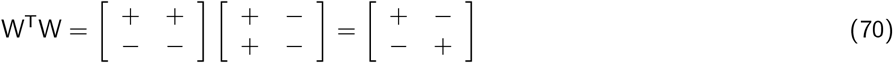

The apparent difference in sign structure in WW^⊤^ and W^⊤^W illustrates the non-normality of W: a non-normal matrix does not commute with its transpose such that WW^⊤^ ≠ W^⊤^W. For any W that obeys Dale’s law WW^⊤^ has strictly non-negative entries, while W^⊤^W has non-negative diagonal blocks, and non-positive off-diagonal blocks. Based on this structure, we can derive sign-constraints on the leading left and right singular vectors of W. The leading left singular vector is chosen as

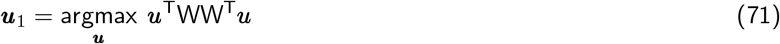

To maximise this quantity, ***u*** will need to have non-negative entries and be of the general form of an E/I co-activation pattern:

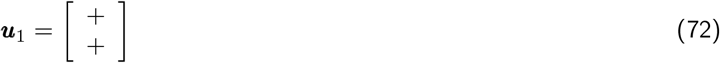

This result also relates to the Perron-Frobenius theorem for non-negative matrices,^117^ which allows one to derive conditions for which ***u***_1_ would have strictly positive entries. Similarly, the leading right singular vector is chosen as

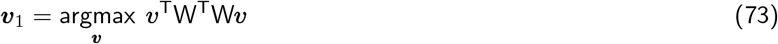

To maximise this quantity, ***v*** will need to have non-negative entries for E-cells and non-positive entries for I-cells, taking the form of a difference pattern:

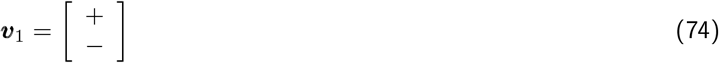

Lastly, the leading singular value is the square-root of the leading eigenvalue of WW^⊤^ or W^⊤^W. We can therefore write

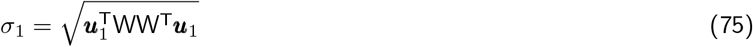

If we assume that each entry in the dynamics matrix W scales according to 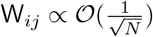 (a common assumption in studying random connectivity matrices linked to the stability of the resulting dynamical system^106,118^), then multiplying ***u***_1_ with each column in W represents a sum over *N* positive terms of 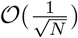. Thus, the leading eigenvalue of WW^⊤^ will scale with 𝒪(*N*), and we arrive at the result

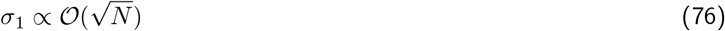

We can use these general properties of dynamics matrices obeying Dale’s law in order to understand the alignment of the network’s task-activity space with the simulation vector. Consider the one-time response of the network, to an arbitrary input pattern ***ξ***:

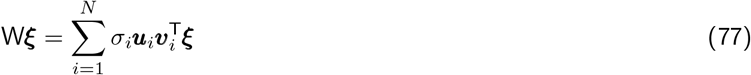

The alignment of the network response to this random input with the co-activation pattern ***u***_1_ can be computed as

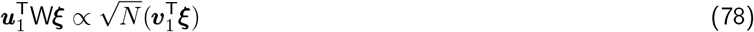

This means that any small projection of an input along the difference pattern ***v***_1_ will get amplified by a factor of 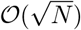 along the co-activation pattern ***u***_1_. For large network sizes, and a randomly chosen ***ξ*** ∼ 𝒪 (0, I), the expectation and variance of projections along the difference pattern are

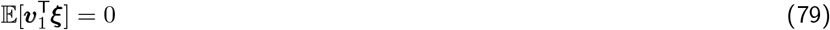

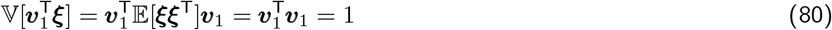

Thus, by chance, we would expect to see variance of 𝒪(*N*) along the E/I co-activation pattern in response to noisy inputs. This shows that activity along ***u***_1_ may also reflect amplified noise. In addition to this, potential coupling of response patterns into the differential mode over multiple time-steps may further contribute towards variance along ***u***_1_.

In summary, the largest singular value (corresponding to the maximum amplification) of any connection matrix satisfying Dale’s law scales with the square-root of the network size, and is always associated with a left (output) row-vector in which all the elements have the same sign, and a right (input) column-vector in which elements corresponding to E-neurons and to I-neurons have opposite signs. The leading singular vectors will depend on the pattern of structured dynamics that the network implements, but the sign constraints we have shown above apply nonetheless. These results can hence explain why we see additional network variance long the E/I co-activation pattern, even when it is not used to reproduce the data, or has any dynamical relevance as it decays rapidly under E/I balance. These theoretical considerations explain why the stimulation has an above-chance alignment with the task-activity space, even when it is misaligned by chance with the low-dimensional task-dynamics space defined through J.

### 15 Evaluating model responses to ICMS

↪ c.f. Fig. 6

To test whether such a description is consistent with the E/I network model, we fit the same model class as before to PSTHs recorded in the absence of stimulation. The peri-stimulation responses appear to follow stimulation-dependent, nonlinear dynamics that depend on unknown circuit-level effects of ICMS and are hard to interpret using our modeling framework. We thus focus our analysis on the period from the last time-point of stimulation onward and investigate whether the observed population responses can be explained using a network model trained on non-stimulated data alone.

Based on previous analyses, we can model the stimulation responses on ICMS trials as a combination of the underlying task-relevant responses on non-stimulated trials, taken to be the posterior mean PSTHs as inferred in our the model 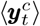, an additive change in firing rates, 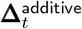, and a residual response component 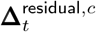 that captures the stimulation-induced distortion of the underlying task-geometry. We thus have

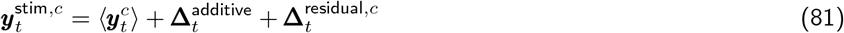

Letting the stimulation delta 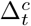 denote the total stimulation effect, taken to be difference between stimulated and expected non-stimulated PSTHs, we have

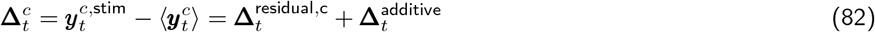

#### 15.1 Additive response component

We hypothesize that the additive response component, analogously to optogenetic stimulation, can be characterized by the response properties of the network to an additive random perturbation. We therefore model it by choosing a random Gaussian vector, ***ν***^rand^ ∼ 𝒪(0, *I*), in network space, and take it as the state at the end of stimulation. This randomly chosen state, by chance, predominantly aligns with dimensions outside of the task-dynamics space. To capture the influence of the resulting activity patterns on data, we need to learn a set of linear weights. To do this, we choose a random size 5 × *K* sample from the network responses evolving forward from the random initial state ***ν***^rand^. We then use this sample from the network to learn a set of linear regression weights that allow to reconstruct the stimulation delta, 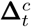, on all but one of the target/timing combination. The set of weights and network responses are then used to obtain a prediction for the additive response component of the held-out condition. We repeat this over 100 random initial vectors ***ν***^rand^ and average the resulting predictions to obtain our final cross-validated estimate of the stimulation response component that can be fully explained as additive and random. Intuitively, our approach aims to “explain away” the condition-invariant response component that can be captured by a random additive perturbation in the network. Based on the results we derived in the context of optogenetic stimulation, we expect any random additive perturbation to miss the task-dynamics space by chance and decay, while also resulting in non-normal amplification along the E/I co-activation pattern. Having an estimate of the additive response component, we can subtract it from the overall stimulation delta to focus our further analyses on structure in the residual response component.

#### 15.2 Residual nonlinear response component

We subtract the cross-validated prediction of the additive component from the stimulation delta to obtain an estimate of the residual response component.

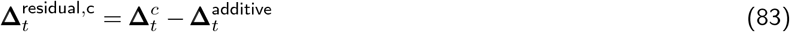

We are interested in investigating whether this response component is consistent with the network dynamics inside the task-dynamics space, and whether these responses can be modelled by finding an initial condition inside the task-dynamics space using the model parameters that were identified from non-stimulated activity alone.

#### 15.3 Correlation of model predictions with residual response component

↪ c.f. Fig. 6f

We next wanted to test whether the dynamics that were estimated from neural activity during normal reaches (in the absence of stimulation) are predictive of the post-stimulation responses. We hypothesize that the residual response component reflects a perturbation inside the task-dynamics space, in which case the post-stimulation responses should reflect signatures of the underlying local population dynamics.

To evaluate the model predicted time-course of recovery from stimulation, we take the residual response component at the last time-point of stimulation and project it into the learned task-dynamics space of the model to obtain a lower-dimensional initial condition inside the task dynamics space

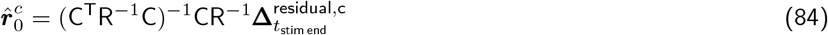

where *t*_stim end_ denotes the time index corresponding to the last time-bin during stimulation. We then use the estimated model dynamics inside the task dynamics space to predict forward in time, starting from 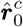:

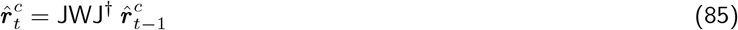

We next project this back out to neural space to obtain our prediction 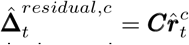 and compute Pearson’s correlation coefficient between the model prediction, and the component of the residual stimulation component that aligns with the task-dynamics space, computed as 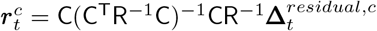. We include data points from the first 200ms after the end of stimulation.

As a control, we repeat the same analysis but evaluate predictions 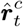 using a dynamics matrix with the same eigenvectors and eigenvalues as those that were learned in the model, but shuffled eigenvalue/vector pairings. This demonstrates that the obtained correlation coefficients are larger than would be expected purely based on a stable system that decays with slow time-constants.

#### 15.4 Upper bound on correlation coefficients with stimulation delta residuals

↪ c.f. Fig. 6f

The correlation coefficient between the empirical residuals ***r***_*i*_ and the model predictions 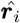 would be maximal if 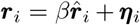. Here, *β* is a scalar and ***η***_*i*_ is the noise covariance in the residual (the same as the noise covariance on stimulation trials since we subtract noiseless quantities from the PSTH to compute the residuals).

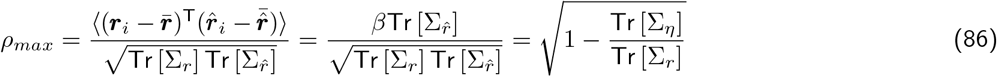

This shows that the maximum correlation we could possibly obtain is small when the residual variance is mostly due to noise, which is relevant for several datasets from Monkey V (Fig. 6f).

#### 15.5 Inferring initial conditions for ICMS responses

↪ c.f. Fig. 6g

The correlation coefficient analysis demonstrates that our model predicts the time-course of the residual response component better than would be expected by chance. Instead of using the raw projection of the residual response component into the task-dynamics space, we next use the model to infer an initial condition at the end of stimulation. This flexibility in finding an initial condition improves the agreement between model predictions and empirical responses even further. We evaluate the fraction of signal variance of the entire post-stimulation responses that can be explained by modeling the stimulation responses as a sum of the model-inferred posterior means on non-stimulated trials, the cross-validated response component to a random additive perturbation and the model prediction of the residual component based on inferring an initial condition inside the task-dynamics space and predicting forward in time using the learned dynamics within this space.

---

**Extended Data Fig. 1.**
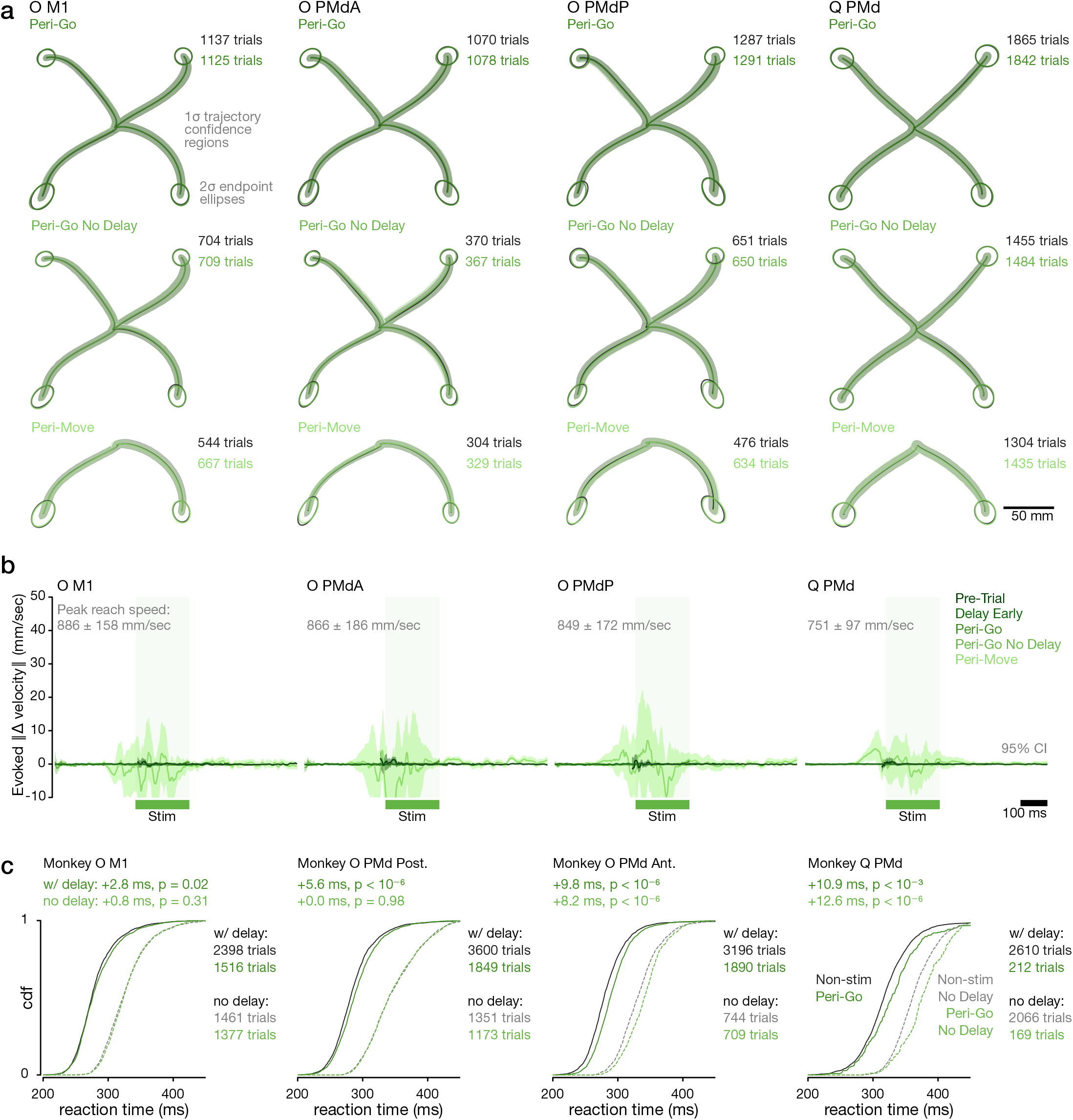
Optogenetic stimulation in motor cortex does not alter reach kinematics. **a:** Mean hand path with ±1*σ* shading for stimulated reaches vs. non-stimulated reaches. Ellipses denote 95% C.I. of reach endpoints. Each row corresponds to a different stimulation timing. Pre-trial and Delay Early stimulation kinematics (not shown) were also identical. Each column corresponds to a simulation site. Trial counts sum across the four reach directions. **b:** Time-courses of the norm of the evoked difference in velocity vector, computed using an unbiased estimate of difference in velocity vectors (which can be negative). For each stimulation condition, we compare the velocities against the same time-points in otherwise equivalent non-stimulation conditions. Shading indicates 95% C.I.s. **c:** Optogenetic stimulation at the go cue slows reaction time (RT). Cumulative distributions of reaction times for non-stimulated (black) vs. stimulated (green) conditions with (solid) and without (dashed) a delay period preceding the go cue. Statistical significance assessed via Mann-Whitney *U* test. The difference between with delay and no delay trials provides a sense of scale of RT benefit incurred through advance preparation.↑ Go back

**Extended Data Fig. 2.**
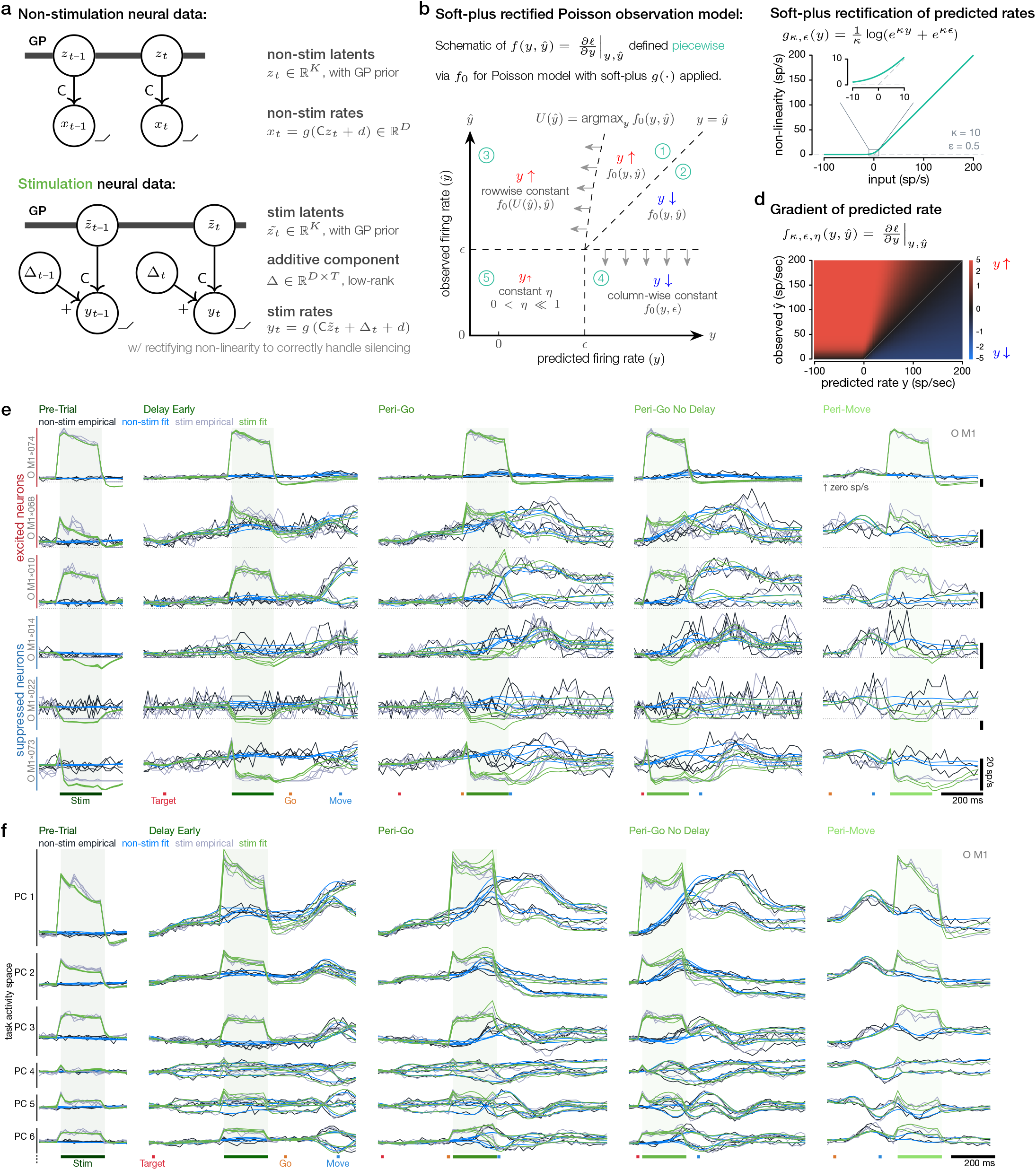
Latent variable model decomposes perturbation effects. **a:** Graphical model of latent variable model (LVM). Circles (nodes) indicate random variables. Arrows indicate a statistical dependence of the recipient node on the source node, with uppercase letter labels indicating matrix multiplication. In non-stimulated trials, the model is similar to Gaussian Process Factor Analysis, as firing rates are constructed from a afine readout of *K* = 6 latent variables with a Gaussian process (GP) prior to ensure temporal smoothness. The linear readout C a matrix with size neurons (*D*) by latents (*K*), and *d* is the offset vector away from the origin. Unlike GPFA, C is fixed and identified using factor analysis, and the autocorrelation timescale of the GP prior is also fixed and constant across latents. For stimulated trials, firing rates are constructed from a distinct set of inferred latents, through the same linear readout and with the same prior, plus a condition-invariant, additive Δ term. The autocorrelation of the GP prior for the stimulation latents is disconnected at the onset and offset of stimulation. The additive Δ, a matrix of size neurons (*D*) × time (*T*), is constrained to be zero before stimulation onset and to be low-rank (rank 5). This constraint ensures that Δ captures the shared timecourses of the additive responses across neurons. Importantly, the neural dimensions addressed by the latent dimensions through C and through Δ (their column spaces) may overlap, depending on the fitted value of Δ. This allows the additive term to capture additive influences on firing rates outside the subspace which explains non-stimulated task variability. In this model, the fitted parameters are the ***z***_*t*_, 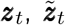, and Δ. **b,c:** Predicted firing rates are computed from the projected latents plus the additive Δ and transformed through a specific rectifying nonlinearity *g*(·). The likelihood of observed spike counts given these rates is evaluated under a Poisson observation model. The rectifying nonlinearity *g*(·) applied to predicted rates is a critical feature of the model to correctly handle neurons whose firing is silenced by stimulation. We chose a soft-plus rectification, plotted in (c), that resembles ReLU but with a softer corner (inset). To gracefully fit the model in light of this rectification, we designed a log-likelihood function *f* (*y, ŷ*) for the observed spike counts *ŷ* (or equivalently 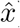) given the predicted rate *y* (or *x*), as illustrated in (b). The design ensures that the model effectively treats neurons silenced by stimulation as censored observations. We construct this function implicitly in terms of its gradient with respect to *y*, and define the gradient piecewise (see Methods §8.4). In (b), each piecewise region is labeled in green; red and blue labels indicate whether the gradient descent acts to increase and decrease the predcited rate. In regions ➀ and ➁, we have a standard Poisson log-likelhood surface where the predicted rate y has been transformed by *g*(·). Region ➂ extends leftwards from the local maximum in each row and simply corrects a non-monotonicity induced by the soft-plus rectification, ensuring that the gradient acts to increase low predicted firing rates when spikes are observed. Region ➃ acts to reduce high predicted rates when the observed neuron fires very few spikes, indicating strong suppression. Lastly, region ➄ aids the optimization by gently penalizing very low or negative predicted firing rates when the neuron is suppressed. **d:** The resulting gradient of the log-likelihood function with respect to predicted rates. Note the logarithmic color scale to aid visibility. **e:** The LVM accurately fits individual neuron responses with both excitatory and suppressive responses to stimulation. Traces show empirical and LVM-fitted trial-averaged firing rates. LVM-fitted rates are not rectified in this visualization to highlight the model’s accurate treatment of suppressed neurons. Dotted horizontal line marks zero spikes/sec. Stimulation timings are separated into columns, with each trace corresponding to a reach direction. **f:** Same as Fig. 2h but with empirical and LVM-fitted trial-averaged firing rates projected into leading six task activity space dimensions. ↑ Go back

**Extended Data Fig. 3.**
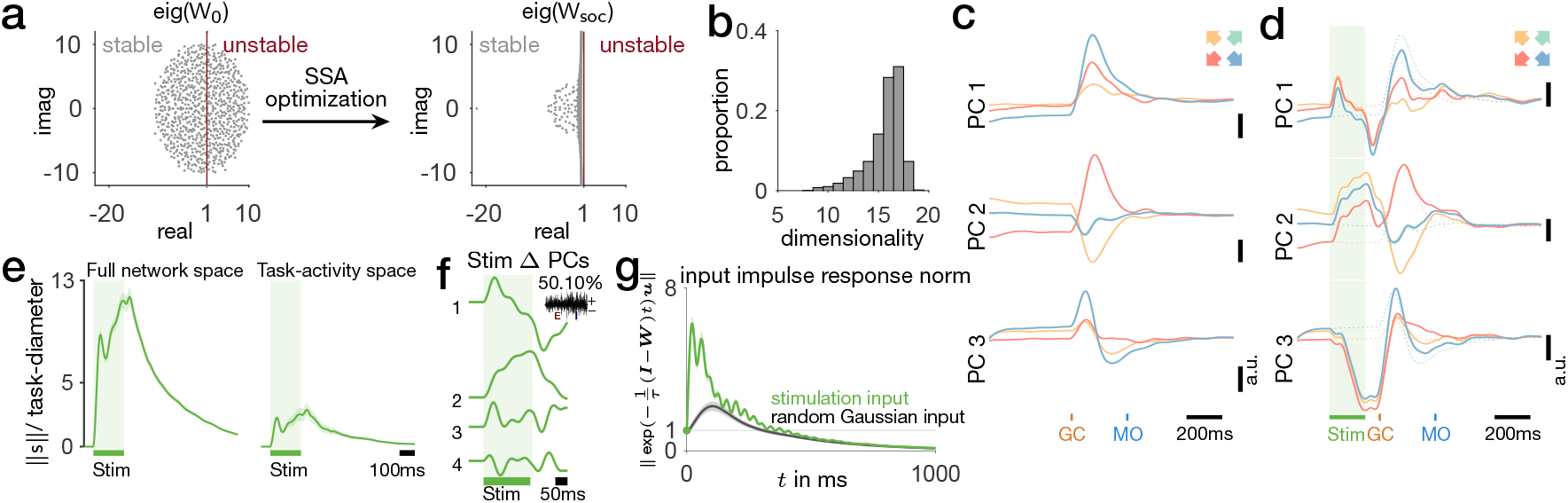
Perturbations trigger long-lasting responses in Stability Optimized Circuits (SOCs). **a:** A highly unstable initial connectivity matrix is optimized by minimizing Smooth Spectral Abscissa, following the approach in [14]. After optimization, the resulting dynamics are stable and can produce complex and slowly varying activity patterns, reflected in the large number of eigenmodes close to the stability line. **b:** The distribution of the number of dimensions needed to capture 95% of the variance produced in response to four initial conditions (see Methods §6), across 20 different realizations of a trained SOC with 400 excitatory and 400 inhibitory units and 100 random samples of initial conditions. **c:** Example projections of SOC activity in response to four initial conditions along its leading three Principal Components (PCs). **d:** The SOC activity in the presence of stimulation-related inputs, projected along the same PCs as in (**c**). **e:** Task-diameter normalized stimulation distance in the full network space and the networks task-activity space (defined by the PC capturing 95% of the variance produced in response to the four initial conditions) for an example SOC. **f:** Projections of the difference between stimulated and non-stimulated SOC activity projected along its leading four PC directions. Inset indicates the loading vector and variance explained by the leading PC. **g:** The average impulse response norm of an example SOC across 100 randomly sampled stimulation vectors (green) and random Gaussian vectors (black). Shaded regions indicate ±2 standard deviations. ↑ Go back

**Extended Data Fig. 4.**
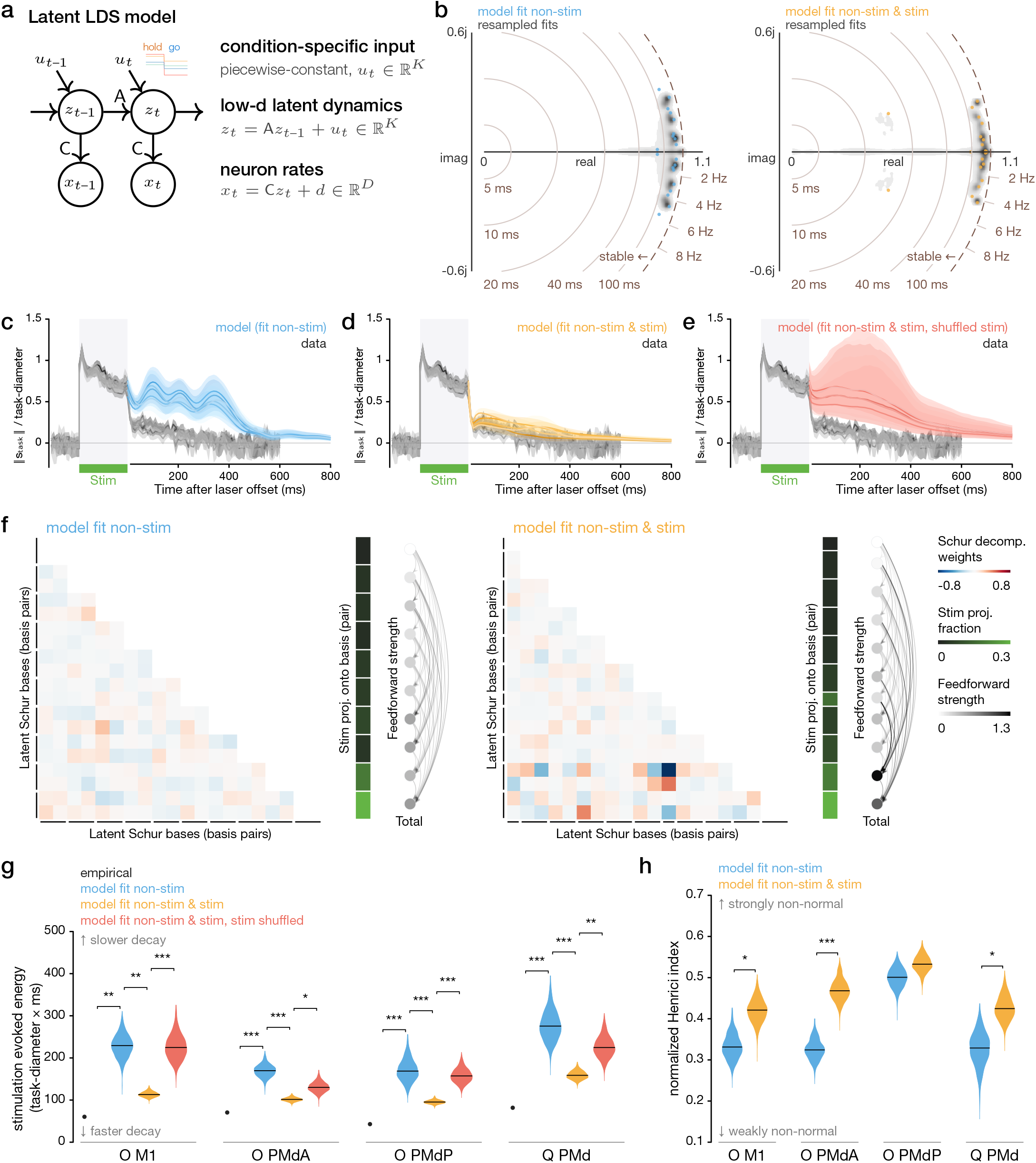
Linear, latent dynamical systems (LDS) models fit to task-related activity predict slow decay of stimulation, but find a fragile, non-normal solution if fit to stimulation responses as well. **a:** LDS graphical model. Circles (nodes) indicate random variables. Arrows indicate a statistical dependence of the recipient node on the source node, with uppercase letter labels indicating matrix multiplication. In the model, low dimensional latent variables ***z*** evolve according to the linear dynamics matrix A and piece-wise constant inputs ***u*** which cause the transition from preparatory to peri-movement neural dynamics. **b:** The timescales of the dynamics can be summarized by the eigenvalues of A. Colored dots indicate eigenvalue locations in the complex plane for the fitted model for the model fit only to non-stimulated activity (left, blue) and the model fit to both non-stimulated and stimulated activity (right, yellow). Gray shading indicates density of eigenvalues for fits to trial-resampled data. Dashed brown circular arc outlines the stable region; light brown circular radii and polar tick marks indicate charateristic timescales and oscillation frequencies, respectively. The model fit to non-stimulated activity learns uniformly slow dynamics, whereas the model fit to both learns a pair of eigenvalues corresponding to a mode which decays more rapidly. **c,d,e:** Time-courses of the length of the perturbation vector (stimulated firing rates minus non-stimulated firing rates at each point in time, for each condition), normalized by the task-diameter. Gray traces indicate the empirical distance measure within the task activity space, as in Fig. 2o, right. Colored traces indicate LDS model predictions, where the initial state was set to best the firing rates at the last time bin of stimulation and the model was run forwards in time. **c:** The model fit to non-stimulated data predicts a slow decay of the perturbation (this is identical to Fig. 2p). **d:** Model fit to both non-stimulated and stimulated data correctly predicts a rapid decay of the perturbation. **e:** However, this solution is fragile, in that it decays quickly only the particular pattern of stimulation used to fit the model. We would expect that the particular pattern of evoked firing rates would depend on the levels of opsin expression in each neuron, which depends stochastically on viral infection. We therefore generated shuffled stimulation vectors by randomly reordering the values of the empirical stimulation vector over neurons. The model fit to both non-stimulated and stimulated data again predicted a slow decay of these shuffled stimulation vectors. **f:** Non-normal dynamics facilitate rapid decay of the stimulation vector in the model fit to both. Heatmaps show the real Schur decomposition of the dynamics matrix (excluding the diagonal). Along each row and column are mutually orthogonal bases within the latent space, either individual bases corresponding to individual eigenvalues, or more commonly, basis pairs corresponding to complex conjugate eigenvalue pairs, as indicated in the margins. The Schur decomposition visualizes the strength of functionally feed-forward influences from the basis in each row onto the basis in each column. We plot this feed-forward coupling weights in the blue-red heatmap and in the the graph edges (grayscale arrows) in the right margin. The total, effective feedforward input for each mode is indicated in the graph nodes (grayscale circles arranged vertically). The Schur decomposition is not unique and depends on the ordering of the eigenvectors used in its contruction. We performed the decomposition so that latent bases/basis pairs on which the stimulation vector had the largest projection (indicated in the vertical green bar) appeared closest to the bottom. The main result is that feedforward coupling weights are uniformly weak in the model fit to non-stimulated activity (left), suggesting approximately normal dynamics. In contrast, the model fit to both exhibits strong feedforward coupling onto the bases with the strongest stimulation responses. This feedforward coupling allows these bases (and the corresponding neural dimensions in the task activity space) to exhibit large variance during non-stimulated conditions, but to rapidly decay with little dynamical impact due to a fast decaying eigenvalue. This is conceptually analagous to internal readout dimensions, driven by the other modes but with minimal influence back onto the network dynamics. **g:** Quantification of the normalized stimulation evoked energy following stimulation offset, which equals the integral under the timecourse curves in (c-e). Black dot indicates empirical evoked energy with 95% confidence intervals (invisible behind dot). **h:** The normalized Henrici index of each model’s dynamics matrix, a measure of non-normality ranging from 0 (normal) to 1 (maximally non-normal). Statistical significance is computed using models fitted to condition-averaged firing rates with trials sampled from replacement. * indicates p < 0.05, ** p < 0.01, *** p < 0.001. ↑ Go back

**Extended Data Fig. 5.**
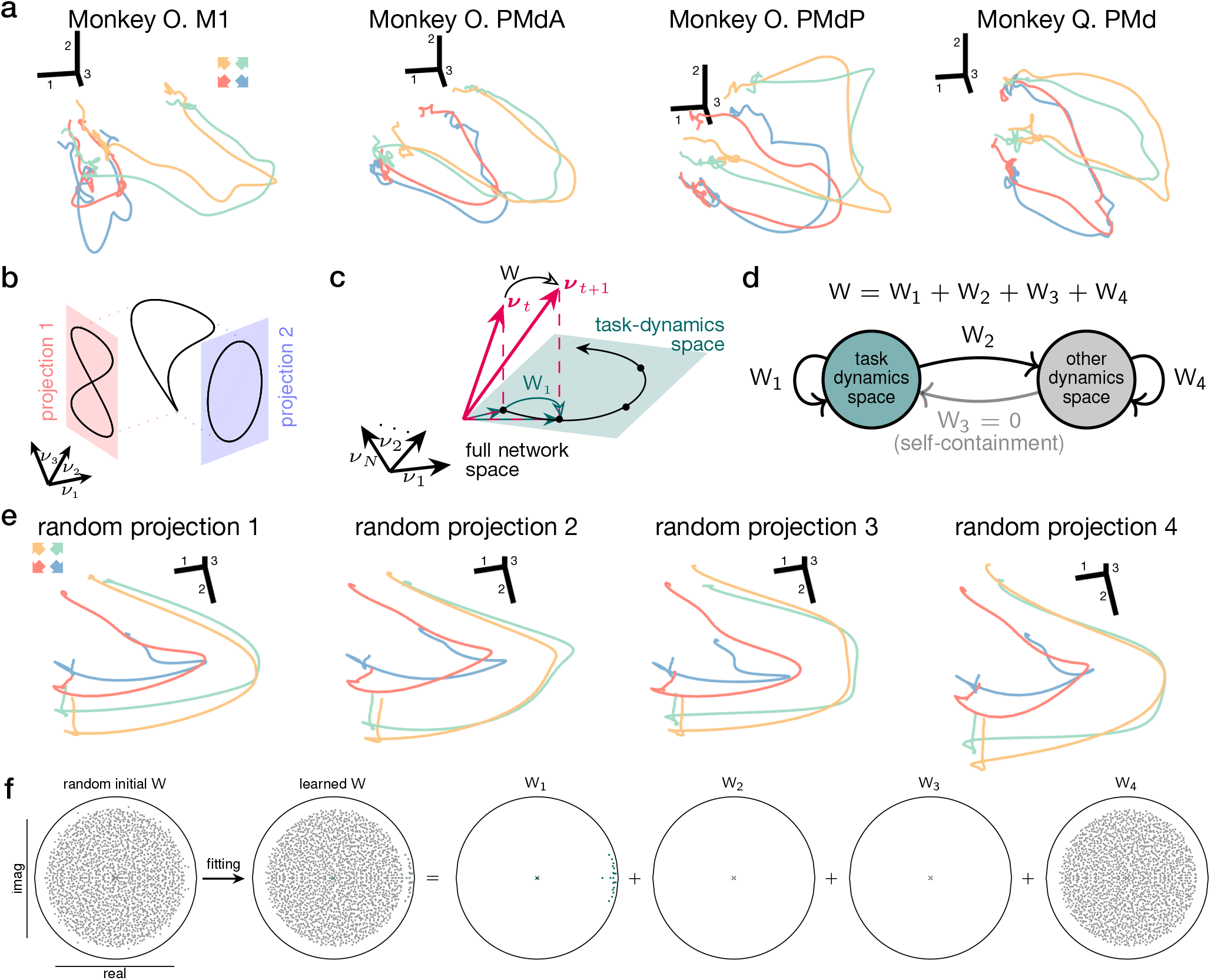
Self-contained low dimensional dynamics. **a:** The leading 30 principal components (PC) of four datasets (different recording sites and monkeys) are rotated using an orthogonal matrix that solves the orthogonal Procrustes problem relative to the PC projections of the PSTHs from Monkey O. PMdP. The first 3 dimensions are shown for each dataset. Projections look similar across recording sites and animals, highlighting the fundamentally low-dimensional and generic structure of the population activity. **b:** Schematic illustration showing that if, rather, population activity was high-dimensional, different random lower dimensional projections of the population would result in different trajectories. This stands in contrast to the similarity of different samples of population activity as shown in (**a**). **c:** Schematic illustration of the self-containment constraint. The temporal evolution of neural activity within the task-dynamics space (shaded green plane) only depends on activity within the task-dynamics space, rather than the full state across all network dimensions. The evolution of neural activity in the full network can be described via the dynamics matrix W (network connectivity), whereas the evolution in the task-dynamics space only depends on low-dimensional dynamics W_1_ (see (**d,f**)). **d:** Decomposition of the network dynamics matrix W into four components, which represent transformations of activity across subspaces. The self-containment constraint sets one of these components to zero, ensuring that there is no transition from outside of the task-dynamics space into the task-dynamics space. This means that the network state strictly outside of the task-dynamics space cannot influence the activity within the task-dynamics space. A low-rank assumption on W would constrain the system to set all components except W_1_ to zero. **e:** Same as (**a**) but using four random projections of the activity of an example network (trained on data from Moneky O. PMdP). Random projections are computed by selecting a random subset of 10 · *K* units from the network. Projections contain similar low-dimensional structure despite sampling different random units from the network, highlighting that the dynamics of the E/I network match the low-dimensional characteristics of recorded neural data. **f:** Eigenspectra within the unit circle in the complex plane. The network connectivity W is initialised randomly and adjusted throughout maximum likelihood learning (see Supplementary Methods). After learning, dynamics within the task-dynamics space are slow and structured (W_1_), but the model largely maintains the random structure of the initialisation outside of the task-dynamics space (W_4_). ↑ Go back

**Extended Data Fig. 6.**
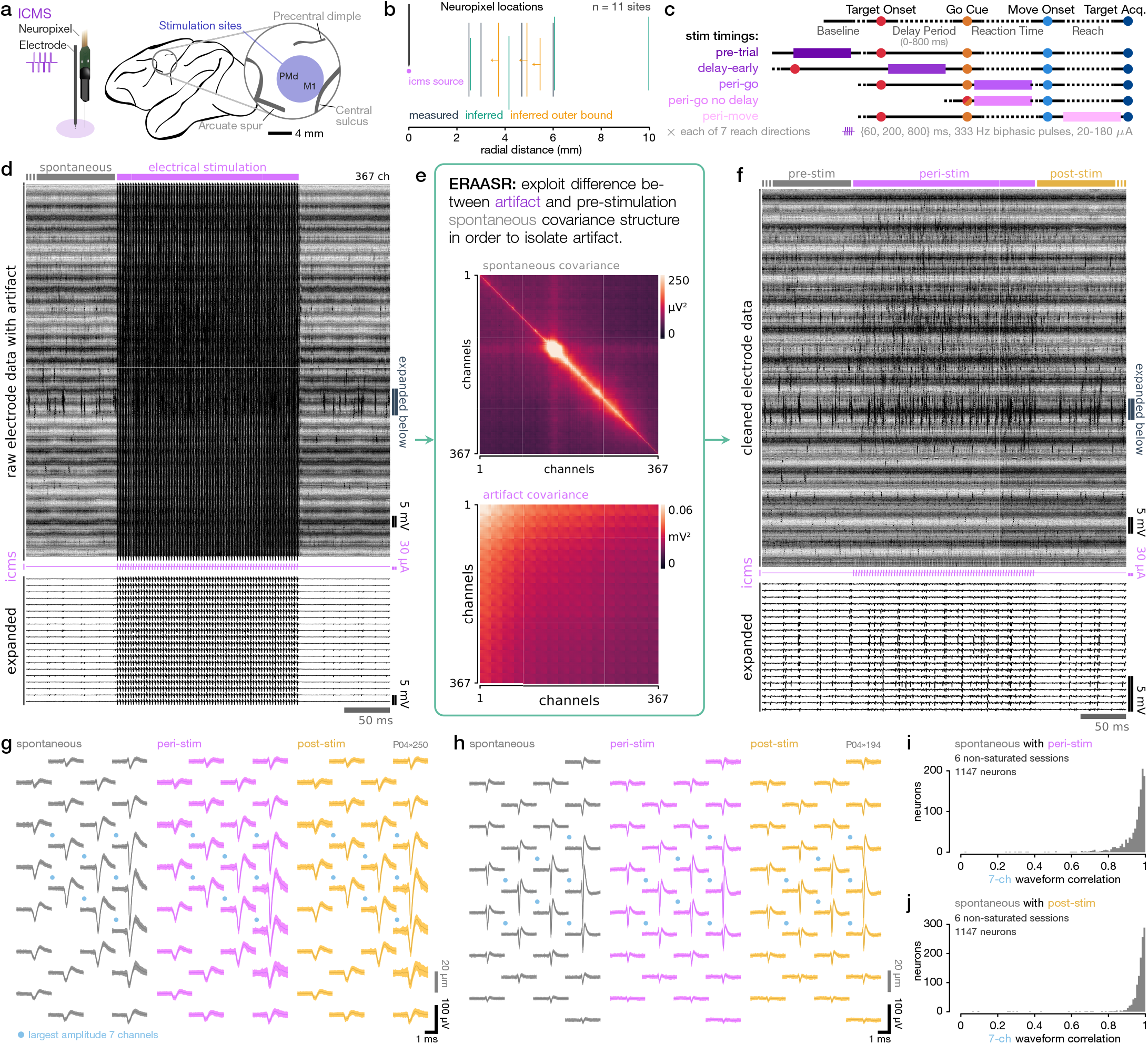
ERAASR_2_ enables recording of neural local activity using Neuropixels probes during and immediately after ICMS. **a:** Schematic of stimulation and recording setup with Neuropixels inserted near tungsten stimulating electrode in M1 or PMd. Anatomical landmarks are approximate based on stereotactic chamber location. A total of 11 Neuropixels penetration sites were recorded, with 21 experimental sessions demarcated by a constant stimulating electrode depth, ICMS amplitude, and ICMS duration. **b:** Approximate Neuropixels probe locations relative to the stimulation electrode. Vertical distances were calculated using micromanipulator depths. For radial distances, *measured* indicates the relative radial distance could be accurately measured on the dural surface based on visible penetration marks. *Inferred* indicates the probe was inferred by fitting a point source voltage model to the electrical stimulation artifact amplitude over channels to recover the relative position (see Methods §3.3). *Inferred outer bound* indicates that the artifact saturated a sufficient number of channels at the amplifier gain settings used for the recording, such that we could only infer an maximal radial distance for the probe, given that the true artifact amplitude was at least as large as the dynamic range. **c:** Diagram of stimulation timings within instructed-delay reaching task; identical to optogenetic stimulation timings, but with seven reach directions instead of four. **d:** Neuropixels recordings through an example 200 ms ICMS stimulation. Each row corresponds to a recording channel, sorted vertically down the Neuropixel, with a subset of channels expanded below. Strong artifact dominates the signal on every channel. **e:** ERAASR_2_ exploits the difference in covariance across channels between spontaneous neural activity (top) and the electrical stimulation artifact (bottom). For each trial, it identifies a subspace capturing maximal artifact with minimal spontaneous variance via a manifold optimization problem, and then tries to reconstruct each channel from a low-dimensional set of artifact basis signals, and then keeps the residual. **f:** Same signals as in (**d**) after cleaning with ERAASR_2_. **g,h:** Multichannel waveforms of two example neurons as detected and sorted by Kilosort2 during the pre-stimulation (spontaneous), peri-stimulation, and post-stimulation time windows. Traces show mean ± s.e.m. **i:** Histogram of Pearson correlation between peri-stimulation and spontaneous waveforms for the 6 non-saturated sessions. Correlation was computed using each neuron’s 7 largest amplitude channels, as marked with blue dots to the top left in (**g,h**). **j:** Histogram of Pearson correlation between post-stimulation and spontaneous waveforms for the non-saturated sessions. Correlations close to 1 indicate that neural spiking signals could be accurately recovered during and after ICMS. ↑ Go back

**Extended Data Fig. 7.**
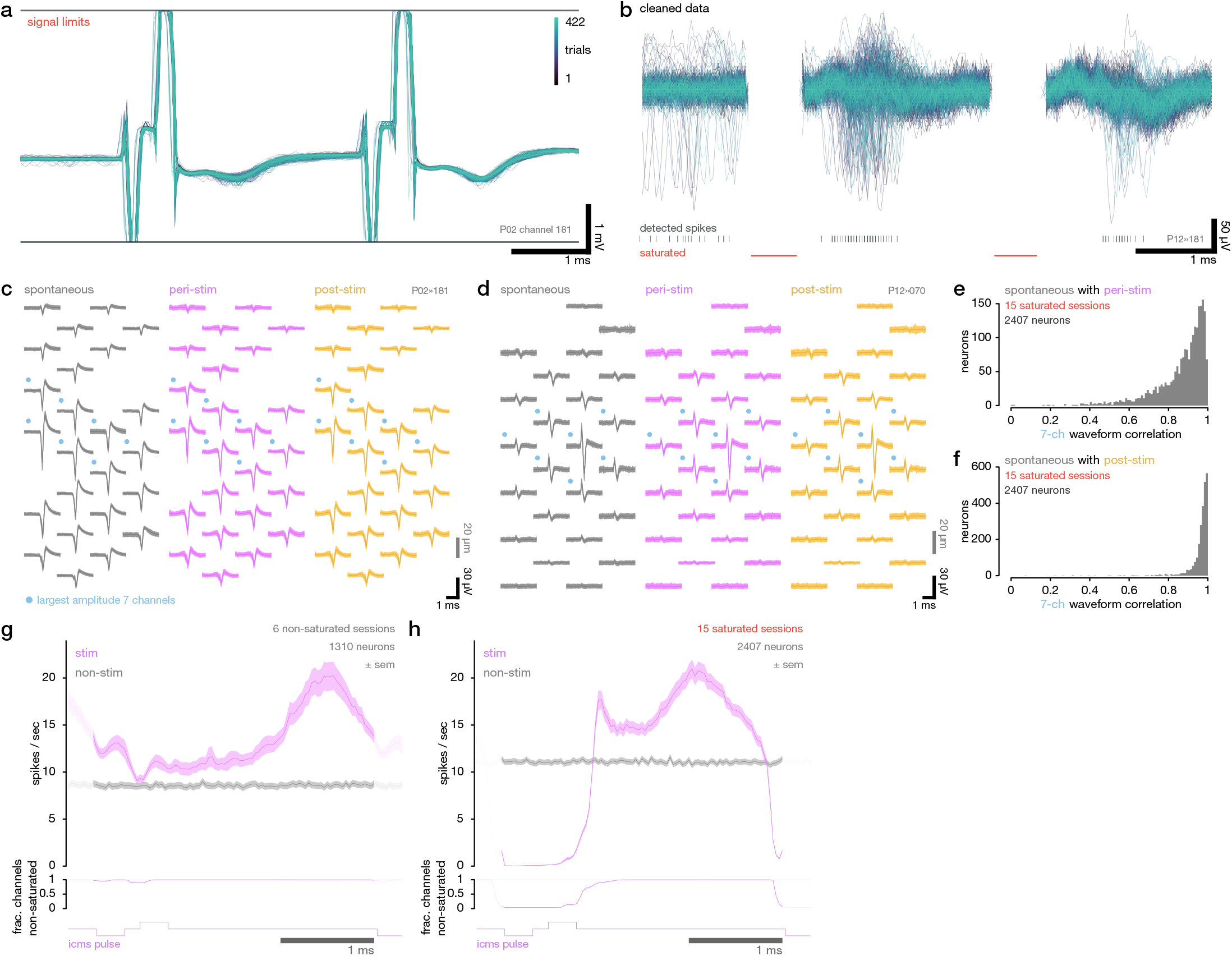
Recovery of spikes in between periods of channel saturation. **a:** Example Neuropixels traces superimposed over all trials for a representatitive saturated channel. **b:** Same traces in (**a**) after cleaning with ERAASR_2_ and blanking the time windows with detected saturation (indicated in red, along with short very intervals between saturation). Detected spikes in this signal for the neuron with the largest spiking waveforms on this channel are plotted below. **c,d:** Multichannel waveforms of two example neurons (from saturated datasets) as detected and sorted by Kilosort2 during the pre-stimulation (spontaneous), peri-stimulation, and post-stimulation time windows. Traces show mean ± s.e.m. **e:** Histogram of Pearson correlation between peri-stimulation and spontaneous waveforms from the saturated sessions. Correlation was computed using each neuron’s 7 largest amplitude channels, as marked with blue dots to the top left in (**c,d**). **f:** Histogram of Pearson correlation between post-stimulation and spontaneous waveforms from the saturated sessions. Correlations close to 1 indicate that neural spiking signals could be accurately recovered during and after ICMS. **g:** (Top) Stimulated vs. non-stimulated firing rates relative to individual ICMS pulses (or the equivalent time in non-stimulated trials), averaged over trials, pulses, and neurons for non-saturated sessions, computed at the time resolution of the original Neuropixels sampling (30 kHz). (Bottom) Fraction of channels with any saturated timepoints. Shaded regions before (after) show the continuation into the previous (subsequent) ICMS pulse. **h:** Same as (**g**) for the saturated-sessions. Note that the timecourse of ICMS-evoked spikes is very similar in the windows between pulses, suggesting that neural states can be accurately estimated despite the saturation during each pulse. ↑ Go back

**Extended Data Fig. 8.**
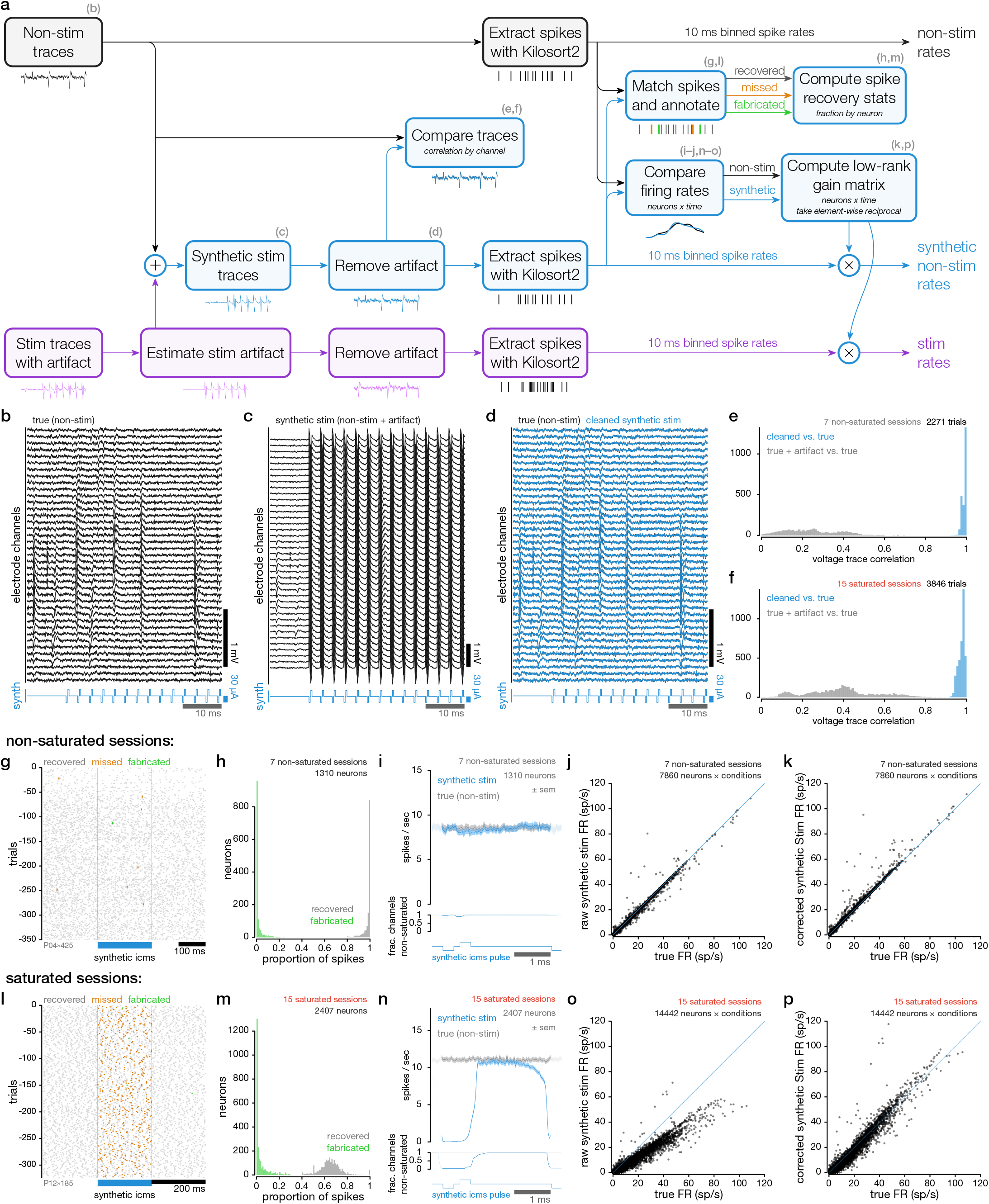
Synthetic stimulation pipeline confirms accurate recovery of spikes throughout stimulation. **a:** Flow-chart schematic of the synthetic stimulation pipeline. Panel letters to the top right of each node indicate the corresponding figure panels related to that step. We use artifact estimated from true stimulation trials, add these artifacts to non-stimulated traces, blank channels where saturation would occur (as is done for the true stimulation traces where saturated occurs), and then perform the identical processing pipeline through ERAASR_2_ and a modified version of Kilosort2 to extract spikes for each neuron (see Methods §3.6). For each neuron, we compare the original non-stimulated spike times to the synthetic pipeline sorted spike times, and annotate each true spike as recovered or missed in the synthetic data. Synthetic pipeline spikes are marked as fabricated if they occur at times with no corresponding spike in the non-stimulated data. We also compute a gain matrix by computing the ratio of the synthetic, condition-averaged firing rates relative to the true, non-stimulated firing rates. We compute a low-rank approximation of this neurons × time matrix, and then use this to compensate for the effects of missing spike times. For both synthetic non-stimulated rates and the true stimulation rates, we divide by this gain matrix element wise to compensate for the effects of missing spike times due to both saturation (if any) and the ERAASR_2_/Kilosort2 pipeline. This process is repeated for each stimulation timing and reach direction, using paired stimulation trials and otherwise equivalent non-stimulation trials. This pipeline achieves two goals. First, we can estimate the effect of the full end-to-end processing pipeline on the Neuropixels recordings due to electrical artifact. Second, we generate synthetic non-stimulated firing rates that simulate the effect of saturation and the end-to-end processing pipeline. We then use the (highly similar) synthetic non-stimulated firing rates in lieu of the non-stimulated firing rates in all subsequent analyses, so that any differences between non-stimulated and stimulated rates are not attributable to saturation or to the cleaning pipeline. Moreover, because spikes evoked by ICMS are preferentially evoked after each individual pulses during the non-saturated timepoints, this gain correction is conservative with respect to the contraction analyses to follow (see note in Methods §3.7). **b:** Example non-stimulation traces for a single ICMS trial. **c:** Synthetic non-stim traces generated by adding electrical artifact from a paired stimulation trial to the traces in (**b**). **d:** Comparison of the ground truth traces in (**b**) (black) against the ERAASR_2_ cleaned traces (blue). **e,f:** Histogram of Pearson correlation coefficients of ground truth vs. ERAASR_2_ cleaned traces, computed on all channels for each trial (blue), versus the correlation coefficients for ground truth vs. the original, artifact corrupted traces vs. ground truth (gray). (**e**) summarizes non-saturated sessions; (**f**) summarizes saturated sessions, excluding the saturated samples. **g:** Synthetic stimulation aligned spike raster, in which each ground truth spike is colored according to whether it was successfully textcolorcRecoveredrecovered (true positive) or missed (false negative) in the synthetic stimulation pipeline. Spikes detected in the synthetic non-stimulation traces with no counterpart in the corresponding non-stimulated trial are marked as fabricated (false positive). **h:** For non-saturated sessions, histogram over neurons of fraction of recovered, true spikes (ideally 1) and fraction of fabricated synthetic pipeline spikes (ideally 0). **i:** (Top) True vs. synthetic firing rates relative to individual ICMS pulses, averaged over trials, pulses, and neurons for non-saturated sessions, computed at the time resolution of the original Neuropixels sampling (30 kHz). (Bottom) Fraction of channels with any saturated timepoints. Shaded regions before (after) show the continuation into the previous (subsequent) ICMS pulse. **j:** Comparison of true vs. synthetic condition-averaged firing rates, for each neuron × condition in non-saturated sessions, *before* the gain correction is performed. **k:** Same comparison *after* gain correction. **l-p:** Identical to (**g-k**) for saturated sessions. ↑ Go back

**Extended Data Fig. 9.**
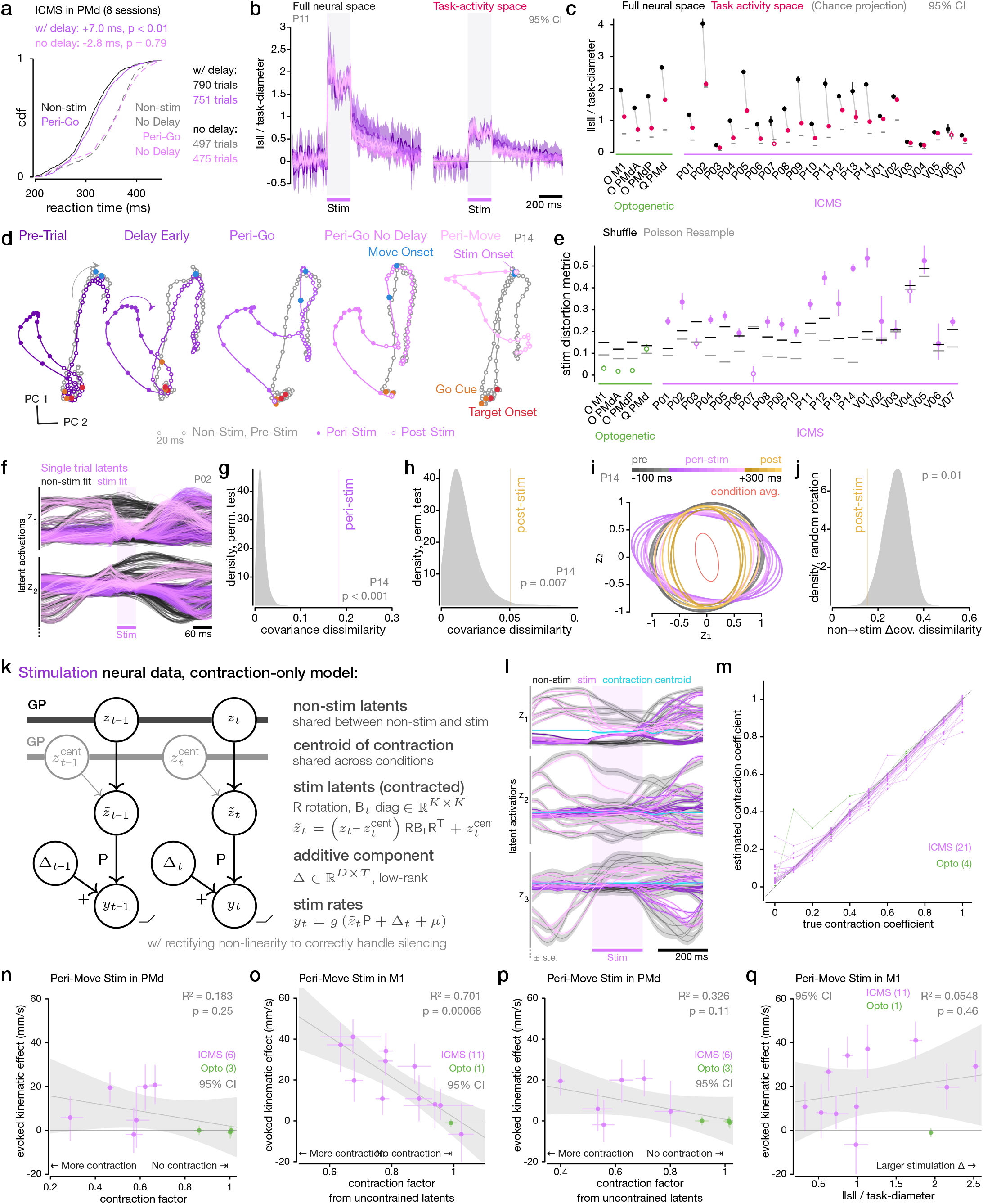
Supplemental ICMS figure. **a:** ICMS in PMd at the Go Cue slows reaction time (RT). Cumulative distributions of reaction times for non-stimulated (black) vs. stimulated (purple) conditions with (solid) and without (dashed) a delay period preceding the go cue. Statistical significance assessed via Mann-Whitney *U* test. The difference between with delay and no delay trials provides a sense of scale of RT benefit incurred through advance preparation. This effect size is smaller than reported in previous studies for subthreshold PMd stimulation^60,62^ as we did not optimize stimlation amplitudes relative to the threshold for evoking movement. **b:** Timecourse of normalized stimulation vector length for O. M1 in the full neural space (left) and projected into the task activity space (right). Each trace represents a stimulation timing colored as in Fig. 5a. **c:** Maximum normalized stimulation vector length in the full neural space (black) and projected into the task activity space (red) for all optogenetic and ICMS sessions. Chance projection of the full stimulation vector into the task-activity space is marked by gray lines. **d:** State space visualization of stimulation evoked displacements of neural trajectories within the task-activity space (PCs of non-stimulated trial-averages) at site P14. Same as Fig. 5f but with all stimulation timings for a single reach direction. **e:** Shape similarity metric quantifying distortion between non-stimulated and stimulated latents for all optogenetic and ICMS sessions. Shuffle and Poisson resample lines indicate significance thresholds (*α* = 0.05) under two null hypotheses where the latents are identical (see Methods §8.3); filled circles indicate statistically significant responses. **f:** Single trial latents inferred by a single trial variant of the latent variable model. **g:** Vertical line indicates normalized dissimilarity metric between the single trial latents’ covariance structure (around their condition means) for stimulated trials compared to non-stimulated trials, computed during stimulation. Gray density indicates the dissimilarity with the trial labels permuted. **h:** Same as (**d**) except for post-stimulation. **i:** Change in single trial latent covariance in the first two latent dimensions as a function of time before, during, and after stimulation. This is computed as stimulation covariance pre-whitened by the non-stimulated covariance, such that no change in variability would appear as the unit circle. Changes in single trial covariance immediately post-stiulation appear similar to the changes in the covariance of the condition means around their centroid (red). **j:** Normalized dissimilarity metric between the change in covariance exhibited by single trials immediately post-stimulation vs. the change in covariance exhibited by the condition means. Density shows the distribution in which the changes in the covariances are randomly rotated within the latent space. This significantly low value indicates that the structure of single trial variability (around their condition-averaged means) in the latents post-stim is reshaped in a similar manner as the distortion that contracts the condition-averaged latents. **k:** Graphical model of the contraction-only LVM. Circles (nodes) indicate random variables. Arrows indicate a statistical dependence of the recipient node on the source node, with uppercase letter labels indicating matrix multiplication. In contrast to the unconstrained model, stimulation latents are calculated from the fitted non-stimulation latents, through a linear operation. This operator takes the form of a pure time-varying contraction along each dimension independently (based on the diagonal matrix B_*t*_) towards a fitted centroid across conditions. The elements of the diagonal matrix B_*t*_ which control the contraction have a Gaussian process prior, with the autocorrelation disconnected at the onset and offset of stimulation. A fitted rotation matrix R allows the axes of the contraction to be oriented appropriately within the latent space. Stimulated rates are computed from the stimulation latents as in the unconstrained model. The fitted parameters of this model are ***z***_*t*_, ***z***^cent^, B_*t*_, R, and Δ. **l:** Fitted latent activations and latent contraction centroid (cyan) for the leading three latent dimensions with the contraction-only LVM. **m:** Accurate recovery of the true effective contraction coefficients. For each dataset, we took the non-stimulated latents inferred by the unconstrained LVM. We generated synthesized stimulation latents by applying with varying amounts of contraction (true contraction coefficient) to the non-stimulation latents. We then regenerated spike counts from both using the projection matrix C and the fitted Δ from the unconstrained LVM. We then fit the contraction only LVM to these synthesized datasets to estimate the contraction coefficient. **n:** Relationship across PMd stimulation sessions between evoked kinematic effect (integral under curve in (c)) vs. stimulation-averaged effective contraction coefficient calculated from the contraction-only LVM. Shading shows 95% CIs for a linear fit. **o,p:** Same as Fig. 5r and (**n**) but with the effective contraction coefficients estimated *post hoc* from unconstrained LVM latents for the (**k**) M1 stimulation sessions and (**l**) PMd stimulation sessions. **q:** Same as Fig. 5r, but with kinematic effect size regressed on the length of the stimulation-induced displacement vector, i.e. value of the black dots in (**c**). ↑ Go back

## Footnotes

a Equivalent definitions with an quasi-upper triangular T are common; the lower-triangular version leads facilitates a more intuitive visualization of the feedforward description of the dynamics. A complex version of the Schur decomposition is also commonly used.

## Notes

### Competing Interest Statement

Prof. Shenoy serves on the Scientific Advisory Boards (SABs) of MIND-X Inc. (acquired by Blackrock Neurotech, Spring 2022), Inscopix Inc. (acquired by Bruker Nano, Fall 2022) and Heal Inc. He also serves as a consultant / advisor (and was on founding SAB) for CTRL-Labs (acquired by Facebook Reality Labs in Fall 2019, and is now a part of Meta Platform's Reality Labs) and serves as a consultant / advisor (and is a co-founder, 2016) for Neuralink.

